# SMA beta bursts reveal distinct mechanisms of anticipatory motor control in children

**DOI:** 10.64898/2026.03.17.712353

**Authors:** Viktoriya Manyukhina, Fanny Barlaam, Judith Vergne, Anaëlle Bain, Oussama Abdoun, Sébastien Daligault, Claude Delpuech, Karim Jerbi, Sandrine Sonié, Mathilde Bonnefond, Christina Schmitz

**Author notes:** Corresponding author: Viktoriya Manyukhina INSERM, INRIA CRNL - CH Le Vinatier - Bâtiment 462 - Neurocampus, 95 Bd Pinel, 69500, Bron, France. both authors equally contributed to the work (Mathilde Bonnefond, Christina Schmitz).

## Abstract

To produce smooth and coordinated movements, the brain generates anticipatory postural adjustments (APA) – feedforward motor commands that counteract predicted postural disturbances before they occur. Although APA emerge early in life, their maturation extends into early adulthood, yet little is known about the underlying neural mechanisms. Using magnetoencephalography during a naturalistic bimanual load-lifting task, we investigated anticipatory postural control in children aged 7-12 years. In this task, lifting a load from the contralateral postural forearm induces upward elbow rotation, which is anticipatorily counteracted by inhibition of *Biceps brachii* postural activity, stabilizing forearm posture. Surprisingly, despite greater variability in inhibition timing and less efficient forearm stabilization in children, the mechanism of anticipatory inhibition was consistent with that recently described in adults: it was mediated by transient oscillatory beta bursts (19-24 Hz), associated with reduced supplementary motor area (SMA) excitability indexed by high-gamma (90-130 Hz) power suppression, and preceded by directed influences from lateral prefrontal and ventral premotor cortex. However, burst frequency was lower in children, and an additional APA mechanism was recruited: while 19-24 Hz bursts were linked to immediate SMA suppression and stronger muscle inhibition, higher-frequency bursts (24-29 Hz) were associated with delayed (∼100 ms) SMA suppression mediated by alpha activity (8 Hz), which attenuated post-inhibition forearm instability. These results reveal that children in middle childhood already engage adult-like neural mechanisms for anticipatory inhibition, while additionally recruiting a compensatory mechanism to counteract postural imprecision, providing new insights into the neural basis of APA maturation and its alteration in neurodevelopmental conditions.

## Introduction

Producing efficient voluntary movement requires the brain to anticipate and compensate for the postural disturbances generated by its own actions through anticipatory postural adjustments (APA) (Massion, 1992). Although the first signs of APA appear within the first months of life, enabling infants to engage more effectively with their surroundings (Reddy et al., 2013), APA remain among the most slowly maturing aspects of motor development, taking over a decade to reach adult levels (Barlaam et al., 2012). Yet despite their critical role in organizing motor control across a wide range of everyday tasks (Lacquaniti & Maioli, 1989; Massion, 1992), the neural mechanisms underlying APA during development remain largely unexplored, particularly whether and how they differ from those in adults, potentially explaining less efficient postural control in children.

The bimanual load-lifting task (BLLT) provides a powerful and naturalistic window into the mechanisms underlying APA and their development (Hugon et al., 1982). Inspired by the ecological scenario of a waiter balancing a tray of glasses in one hand while lifting a glass with the other, the BLLT requires participants to lift a load placed on the contralateral forearm, capturing a common form of posture-movement coordination in which one hand provides postural support while the other performs a voluntary action. To maintain stability of the postural forearm, the brain must anticipate the upward elbow rotation induced by unloading and compensate for it by suppressing activity in the postural elbow flexors – particularly the *Biceps brachii* – shortly before unloading, as demonstrated in adults (Dufossé et al., 1985; Hugon et al., 1982; Paulignan et al., 1989). Notably, although anticipatory inhibition emerges as the predominant APA strategy by age 7-8 (Schmitz et al., 2002), its timing remains highly variable and, on average, later than in adults throughout adolescence, resulting in less precise postural control (Barlaam et al., 2012; Schmitz et al., 2002), suggesting that the neural mechanisms underlying APA in children may differ from those in adults.

In a recent study (Manyukhina et al., 2026), we demonstrated that in adults, anticipatory inhibition of the postural *Biceps brachii* during the BLLT is mediated by transient beta-band (13-30 Hz) activity – beta bursts, associated with suppression of contralateral supplementary motor area (SMA) excitability, potentially reducing its tonic drive to the *Biceps brachii* and thereby leading to muscle inhibition. Here, using MEG during the BLLT in children aged 7-12 years, we investigated whether less efficient anticipatory inhibition during development reflects a qualitatively different neural mechanism that inherently limits the temporal precision and efficacy of feedforward postural control. Specifically, we hypothesized that ongoing APA maturation in children may be reflected in reliance on the primary motor cortex (M1) rather than SMA-mediated control, consistent with the established role of M1 in acquiring novel posture-movement coordination (Ioffe et al., 2003; Kazennikov et al., 2008; Massion et al., 1999). Alternatively, reduced temporal precision may reflect more sustained beta activity implicated in muscle inhibition, or the engagement of slower alpha-band (6-12 Hz) mechanisms, also linked to motor inhibition (Bönstrup et al., 2015; Hummel et al., 2002; Sauseng et al., 2013). Contrary to our hypothesis, we show that children in middle childhood engage a similar SMA-mediated beta burst mechanism for anticipatory inhibition as adults, while additionally relying on compensatory processes that mitigate the postural consequences of imprecise postural control. Together, these findings reveal a two-component proactive adaptive strategy by which the developing brain counteracts its own limitations, shedding new light on the neural basis of predictive motor control during development.

## Methods

### Participants

Twenty-four typically developing (TD) children aged 7.11 to 12.10 years (mean age = 9.69 ± 1.62 years) participated in the study (7 girls). All participants except two were categorized as right-handed based on a manual laterality questionnaire comprising eight items that assess hand preference in daily activities. Participants were additionally assessed using the French adaptation of the M-ABC (Henderson et al., 2019; Soppelsa & Albaret, 2005) by a trained specialist to ensure that none of them presented with motor delay or atypical motor development. Each child scored above the 15th percentile on the M-ABC (mean score ± SD = 3.1 ± 2.1), confirming a typical level of motor functioning. All children attended school and had no history of neurological or psychiatric disorders. Written informed consent was obtained from all children and their parents prior to participation, in accordance with the Declaration of Helsinki. The study was approved by the French Ethics Committee (CPP Sud-Est III) and authorized by the French National Agency for Medicines and Health Products Safety (ANSM; ID RCB 2014-A01953-44; ANSM reference 141587B-31).

### Experimental task and procedure

The experimental setup for the BLLT has been previously described using EEG (Barlaam et al., 2011, 2018) and MEG (Di Rienzo et al., 2019; Manyukhina et al., 2026). Participants were seated comfortably under the MEG helmet in front of a wooden table, on which the right arm rested. The left, postural arm was positioned adjacent to the trunk, with the elbow fixed using a mobile support that allowed only one degree of freedom: upward and downward rotation. A non-magnetic wristband fitted with a vacuum switch system (30 kN/m²) and a strain gauge was worn on the wrist of the postural arm. Each new trial was initiated only when the child was seated still, had returned to the initial posture, and was attentive.

Given morphological differences among children in body weight, height, and forearm length, the load was set at 350 g for children aged 7-9 years and 400 g for those aged 10-12 years. Slight adjustments were also made based on body mass index: within the 7-9 year age group, heavier children received a 400 g load, while within the 10-12 year age group, lighter children received a 350 g load. This procedure was consistent with previous BLLT studies conducted in children of the same age range (Barlaam et al., 2018; Schmitz et al., 2002). No child reported muscular fatigue.

The experiment included two conditions: voluntary unloading and imposed unloading. In the voluntary unloading condition, participants were instructed to lift the load placed on the wristband of the left postural forearm using the right, motor arm. At the beginning of each trial, participants positioned their right hand above the load. Each trial began with a two-second fixation on a light spot projected onto the wrist, to minimize eye movements during the lifting phase. Once the light faded, participants were free to initiate the lifting movement at their own discretion. They lifted the load approximately 10 cm above the wristband and, after a brief pause, returned it to its initial position, completing the trial. In this condition, APA occurred prior to unloading, reducing the forearm destabilization caused by unloading, as evidenced by relatively small postural elbow rotation, consistent with previous findings in children (Barlaam et al., 2018; Schmitz et al., 2002).

In the imposed unloading condition, a load was suspended from the wristband and released by air pulses (3.5 kN/m²) triggered by the experimenter at unpredictable times. This condition served as a control to assess the efficiency of APA observed in the voluntary unloading condition. Since participants could neither anticipate nor prepare for the sudden unloading, load release produced a clearly visible passive upward rotation of the elbow, accompanied by inhibition of the postural forearm flexors – the so-called unloading reflex (Hugon et al., 1982; Schmitz et al., 2002). In both conditions, task demands implicitly required participants to maintain the postural forearm in a horizontal position, as if carrying a tray without spilling its contents. This horizontal posture thus served as the consistent reference frame across conditions.

The session began with a block of 10 imposed unloading trials, followed by a 45 s rest block during which children were instructed to close their eyes and remain still. The imposed unloading condition was limited to 10 trials given its low variability in children (Barlaam et al., 2012; Schmitz et al., 2002) and to limit task demands; data from this condition were used solely as a baseline to assess forearm stabilization achieved during voluntary unloading. The voluntary unloading condition then followed, consisting of seven blocks of eight trials (56 total), with short rest intervals between blocks during which the experimenter presented a picture of a gift the child could choose at the end of the experiment, to sustain motivation and maintain a pleasant atmosphere. A second 45 s rest block with eyes closed was recorded at the end of the voluntary unloading condition. The session then continued with five additional blocks of eight trials in a learning condition (to be reported elsewhere), and concluded with a final 45 s rest block. The entire session typically lasted less than 2 h, including approximately 45 min for equipment setup and 30 min for experimental recordings.

### Behavioural data acquisition

#### Load-lifting onset detection

The onset of load-lifting (unloading; zero reference time point) was automatically detected as the initial deflection in the force signal recorded by the force-plate sensor attached to the non-magnetic wristband, using a threshold-based function in CTF® DataEditor software. The automatically assigned marker was subsequently visually inspected and manually adjusted when necessary.

#### Elbow joint kinematics

Upward elbow-joint rotation following load release or lifting was recorded in all trials and conditions using a copper potentiometer aligned with the elbow joint axis and sampled at 20 kHz. For analysis, the signal was resampled to 600 Hz. Elbow deflection amplitude was expressed in arbitrary units as provided by the recording equipment.

#### Electromyography (EMG)

Electromyography (EMG) data were recorded at 600 Hz using bipolar surface electrodes (2.5 mm²) placed over the left *Brachioradialis*, *Biceps brachii* (flexors), and *Triceps brachii* (extensor) on the postural arm. For comparability with our previous study in adults (Manyukhina et al., 2026), only the activity of the left *Biceps brachii* was analyzed.

### Behavioural data analysis

#### Single-trial detection of EMG inhibition

To identify anticipatory muscle inhibition, we focused our analysis on EMG activity from the postural *Biceps brachii*. In BLLT research, inhibitory events in postural flexors are traditionally identified through manual inspection of preprocessed EMG signals (Barlaam et al., 2012; Schmitz et al., 2002; Viallet et al., 1992), as trial-to-trial variability in muscle activity and signal-to-noise ratio makes automated amplitude-based detection unreliable. Although expert visual inspection can yield reliable estimates, it introduces potential subjective bias. To address this limitation, we implemented both manual and automated procedures for single-trial detection of inhibitory events in the *Biceps brachii*.

##### Manual detection

For the manual approach, the EMG signal was preprocessed following previously established protocols (Barlaam et al., 2011, 2018; Schmitz et al., 2002). The EMG signal was epoched from -1 to 0.5 s relative to unloading onset, mean-centered, band-pass filtered (25-150 Hz) using a finite impulse response (FIR) filter with a Hamming window, and rectified, such that *Biceps brachii* inhibition appeared as a reduction in EMG amplitude (Fig. 1A).

**Figure 1.**
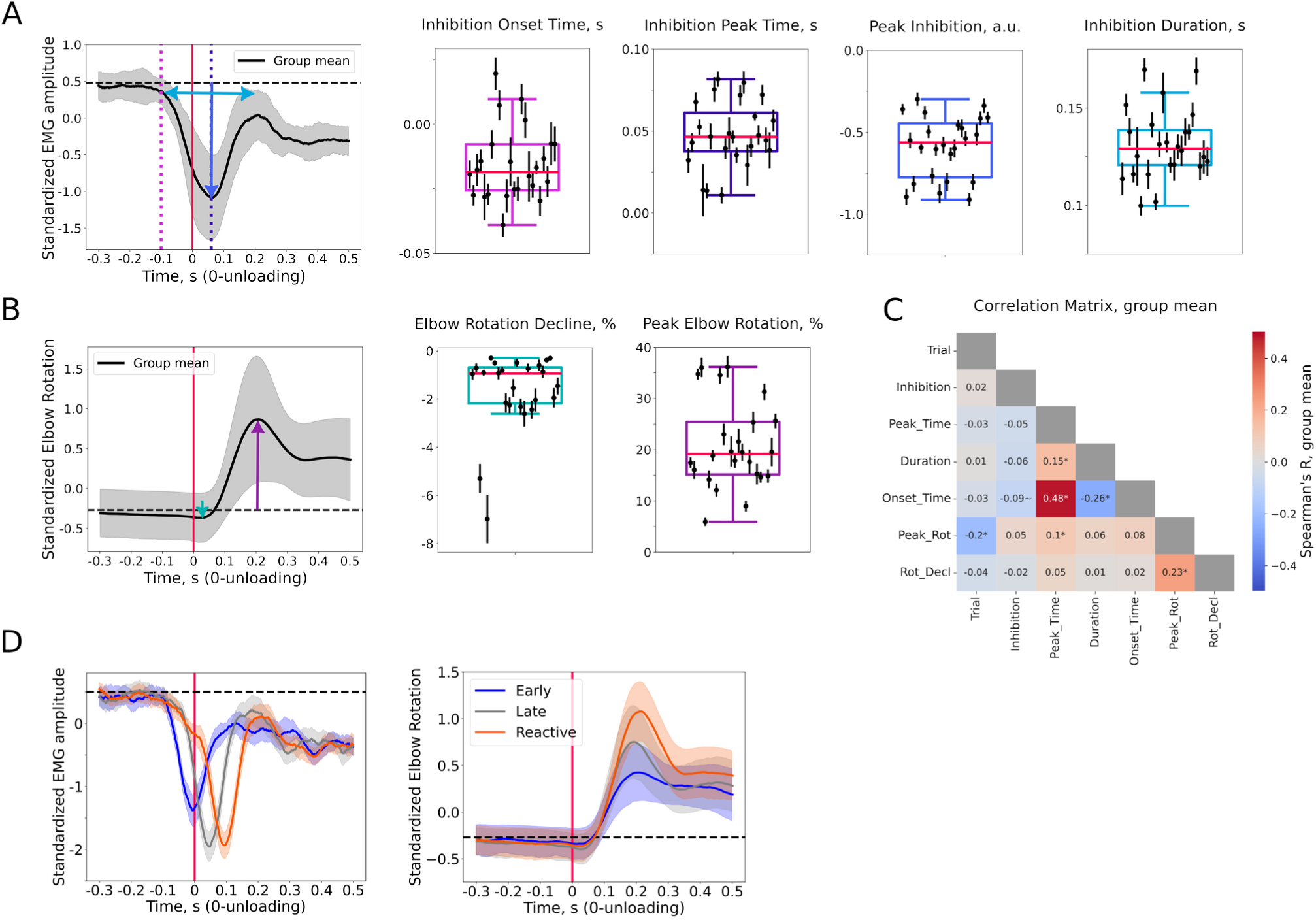
Behavioral analysis results. **A:** Rectified and standardized *Biceps brachii* EMG amplitude modulation over time. **B:** Standardized elbow rotation amplitude modulation over time. In panels **A** and **B**, group means (black line) and standard deviations (gray shading) are shown, with colored arrows and dotted lines indicating the amplitudes of four *Biceps brachii* inhibition-derived measures (**A**) and two elbow rotation-derived measures (**B**). Boxplots for each behavioral measure illustrate between-subject variability. The red line indicates the median, box edges represent the quartiles (Q1 and Q3), and whiskers extend to 1.5 times the interquartile range. Participant-level variability is shown by dots representing the mean values and whiskers indicating the standard error of the mean. **C:** Exploratory correlations between *Biceps brachii* inhibition-derived measures, elbow rotation-derived measures, and trial number, followed by assessment of group-level significance. *p < 0.05; ∼p < 0.15, FDR-corrected. **D:** *Biceps brachii* EMG amplitude and elbow rotation over time, shown for three categories of *Biceps brachii* inhibition: early (anticipatory, peak -70 to 75 ms, onset < 0), late (anticipatory, onset > 0), and reactive (peak 75-150 ms after 0). See Methods for details on anticipatory and reactive inhibition selection.

For each subject, the single-trial EMG was first visually inspected using unloading annotations as a temporal reference to assess the presence and temporal distribution of inhibitory events. Participants differed in the number of identifiable inhibitory events, though reliable inhibitions were observed in all participants. Inhibitory events were characterized by suppression of EMG amplitude to near-zero values, appearing as a fully or partially flattened signal. Inhibitory events were then annotated for each trial when identifiable within a ±250 ms window centered on unloading onset. Consistent with previous studies (Barlaam et al., 2011, 2018; Schmitz et al., 2002), onset and duration were extracted from each identified event. When multiple inhibitions were present, only the first was retained, as its timing is considered most critical for APA efficacy. To improve reliability, annotations were re-examined on a separate day and subsequently reviewed by an experienced specialist. For each examination, subjects were inspected in randomized order.

##### Automated detection

Traditional manual detection assumes that muscle suppression manifests as a global, broadband decrease in activity. However, suppression may be limited to a narrower frequency band, or broadband suppression may be partially obscured by noise, making only the peak detectable.

Manual detection also cannot reliably quantify inhibition strength due to a floor effect: when two inhibitory events both decrease to near-zero amplitude, their relative magnitudes cannot be distinguished.

To address these limitations, we developed an algorithm to automatically identify inhibitory events and estimate their strength using time-frequency EMG analysis. Raw EMG data were epoched from -1.5 to 1 s relative to unloading onset, and single-trial time-frequency analysis was performed using the multitaper method in MNE-Python (tfr_array_multitaper; 30-150 Hz, 1 Hz resolution, 7 cycles, time_bandwidth=10). Power values were log10-transformed and baseline-corrected per frequency using a pre-unloading interval (-1.5 to -0.5 s). The analysis window was then restricted to ±0.5 s around unloading. To avoid detecting random fluctuations, suppression at a given time-frequency point was retained only if the surrounding ±50 ms window contained at least 20 ms of sustained suppression below a predefined threshold (mean - 1.5×SD). The resulting time-frequency matrix was averaged across frequencies and multiplied by -1 to generate a time course of muscle inhibition, with positive peaks reflecting inhibition strength. Peaks were detected using find_peaks() from SciPy (v1.13.1; distance=30, prominence=0.1, width=3), peak amplitude and timing were extracted. Example trials illustrating algorithm performance are provided in Supplementary Materials (Figs. S1A-C).

Although inhibition onset was also estimated, visual inspection revealed that it reflected the moment when suppression became reliably detectable above baseline, rather than the true onset of EMG decrease, resulting in systematically delayed estimates that were correlated with inhibition peak time across trials. The automated algorithm was therefore used to derive inhibition amplitude (Peak EMG Inhibition) and peak time (EMG Inhibition Peak Time), while manual annotations were retained for inhibition onset and duration. The number of trials with identified *Biceps brachii* inhibition ranged from 15 to 56 (mean ± SD: 44.08 ± 9.02) with manual detection and from 22 to 54 (43.5 ± 8.65) with automated detection. Automatic detection identified more inhibition trials than manual detection in 12 participants, fewer in 8, and equal counts in 4, with substantial trial overlap between methods (72 ± 17% relative to total cleaned trials).

#### Single-trial estimation of elbow rotation parameters

Elbow-joint rotation was tracked in the voluntary unloading and imposed conditions to assess forearm stability as an indicator of postural control efficiency, with greater stability reflecting better performance. Following Manyukhina et al. (2026), we estimated two metrics: Peak Elbow Rotation (maximal upward deflection) and Elbow Rotation Decline (maximal downward deflection), representing the two primary directions of forearm instability during voluntary load-lifting in adults. Both measures were estimated at the single-trial level. Fig. 1B shows the grand-average elbow rotation, schematically indicating the amplitudes of Peak Elbow Rotation and Elbow Rotation Decline. On trials with efficient postural control, both measures were expected to approach zero.

The maximal elbow rotation amplitude was manually identified for each trial, following the procedure described in Manyukhina et al. (2026) in adults. Manual annotation was required because small peak elbow rotations could not be reliably identified with automated algorithms. To minimize annotation bias, the procedure was preceded by a training session with an experienced specialist.

Trials from each participant were first previewed to assess within-subject variability, after which peak detection was performed within a 500 ms window following unloading onset. Annotations were re-inspected on a separate day, with participants randomized, and ambiguous trials reviewed with the specialist.

Elbow rotation timecourses were epoched from -1.7 to 1.2 s relative to the annotated peak. Peak Elbow Rotation and Elbow Rotation Decline were estimated automatically as in Manyukhina et al. (2026). Briefly, Peak Elbow Rotation was calculated as the average of the 20 maximal baseline-corrected values within a 165 ms window around the annotated peak, with baseline defined as the 20 minimal values in the -350 to -50 ms interval relative to unloading. Elbow Rotation Decline was computed as the average of the 20 minimal baseline-corrected values from -165 ms relative to unloading to the annotated peak, with baseline defined as the 20 maximal values within the same interval, excluding subsequent positive elbow deflection. Both Peak Elbow Rotation and Elbow Rotation Decline were estimated at the single-trial level, with data and baseline intervals visually inspected. Trials precluding reliable estimation were excluded (0-9 per participant; mean ± SD: 2.64 ± 2.02).

In the imposed condition, Peak Elbow Rotation was estimated as in the voluntary unloading condition, except that the prominent elbow deflection amplitude allowed automatic detection of the peak, as in adults (Manyukhina et al., 2026). Elbow Rotation Decline was not observed in this condition, also consistent with findings in adults, and was therefore not assessed.

#### Relationship between behavioural measures

To assess relationships between elbow rotation and *Biceps brachii* inhibition measures, we performed an exploratory correlation analysis across six metrics, including trial number as an additional variable to evaluate potential changes over time. Within-subject Spearman’s rank correlation coefficients were computed for each variable pair across trials and tested against zero using the Wilcoxon test, with false discovery rate (FDR) correction for multiple comparisons.

Given the well-documented developmental improvement in APA and forearm stability from childhood to adulthood (Barlaam et al., 2012; Schmitz et al., 2002), we also examined whether subject-level metrics varied with age (7-12 years). For this, each of the six elbow rotation and *Biceps brachii* inhibition metrics was averaged across trials per participant, and Spearman correlations with age were computed across subjects, with FDR correction applied.

Previous studies using the same paradigm have shown that earlier anticipatory inhibition onset is critical for efficient forearm stabilization (Barlaam et al., 2012; Massion, 1992; Schmitz et al., 2002). Here, we extended this prediction by testing which *Biceps brachii* inhibition parameters – Inhibition Onset Time, Inhibition Peak Time, Peak Inhibition Strength, and Inhibition Duration, or their interactions – best predicted forearm stabilization, as indexed by Peak Elbow Rotation – the primary source of postural instability in the BLLT (Hugon et al., 1982; Massion, 1992; Viallet et al., 1987). The model was restricted to inhibitions peaking between -70 and 150 ms relative to unloading and included non-linear smooth terms for each predictor and interaction smooths for all predictor pairs. Subject-specific variability was captured using random intercepts and slopes. Model parameters were estimated using restricted maximum likelihood (fREML) via the bam() function from the mgcv package in R (v.1.9.4). The full model specification is provided in Supplementary Materials (Fig. S2).

#### Selection of anticipatory Biceps brachii inhibitions

To investigate mechanisms of anticipatory postural control during development, further MEG analyses focused on anticipatory inhibitory events peaking between -70 and 75 ms relative to unloading (Barlaam et al., 2012; Schmitz et al., 1999, 2002). The early boundary (-70 ms) was guided by previous reports showing that anticipatory elbow flexor inhibition typically occurs within ∼50 ms before unloading (Barlaam et al., 2012; Hugon et al., 1982; Viallet et al., 1987), with a slightly wider margin to account for inter-subject variability. The late boundary (75 ms) was chosen to exclude reactive inhibitory responses, as imposed-condition data showed grand-average *Biceps brachii* amplitude beginning to decrease ∼115 ms after unexpected load release. Although peaks up to 75 ms after unloading may appear late, their onset occurs earlier, and given a mean inhibition duration of 131 ms in this sample, it is unlikely that these inhibitions were triggered by unloading. Such delayed anticipatory inhibitions, with onsets preceding or shortly following unloading, are common in children of this age group (Barlaam et al., 2012; Schmitz et al., 2002). Altogether, this selection likely captures anticipatory responses generated before unloading but delayed due to less efficient anticipatory control in children.

### MRI acquisition

Structural magnetic resonance imaging (MRI) data (voxel size: 0.9 × 0.9 × 0.9 mm; TR = 3500 ms, TE = 2.24 ms) were acquired for all participants using a 3T Siemens Magnetom scanner (MAGNETOM Prisma, CERMEP, France; Siemens HealthCare). T1-weighted images were preprocessed with FreeSurfer’s recon-all pipeline (version 6.0.0; Fischl et al., 2002), which performs motion correction, intensity normalization, removal of non-brain tissue, cortical surface reconstruction, and subcortical segmentation.

### MRI processing

The cortical surface was parcellated into 450 regions using the HCP-MMP1 atlas (Glasser et al., 2016) as implemented in MNE-Python (v.1.7.0; Gramfort et al., 2013). Source-level analysis was restricted to regions previously associated with voluntary unloading in the BLLT and load-lifting in adults (Manyukhina et al., 2026; Ng et al., 2011, 2013a, 2013b; Schmitz et al., 2005): M1, SMA, premotor cortex (PMC), cingulate motor area, prefrontal cortex (PFC), precuneus, supramarginal gyrus, and inferior frontal cortex (IFC). Analyses focused primarily on the right hemisphere, contralateral to the postural arm, to examine pathways specifically implicated in left *Biceps brachii* inhibition, consistent with our adult study (Manyukhina et al., 2026); left-hemisphere contributions were assessed in supplementary analyses. The right cerebellar cortex and basal ganglia (caudate nucleus, putamen, and globus pallidus), previously implicated in the BLLT (Ng et al., 2011, 2013a, 2013b; Schmitz et al., 2005), were also included and parcellated using the FreeSurfer Aseg atlas.

### MEG acquisition

Magnetoencephalography (MEG) data were recorded at CERMEP (France) using a CTF-MEG system (CTF MEG Neuro Innovations, Inc.) equipped with 275 radial gradiometers and 29 reference channels for ambient noise correction. Signals were digitized at 600 Hz and low-pass filtered at 150 Hz. Head position was continuously monitored via three coils placed at the nasion and preauricular points.

### MEG preprocessing

MEG data from the voluntary unloading condition were preprocessed using MNE-Python (v.1.7.0; Gramfort et al., 2013). Raw recordings were aligned to an optimal head position, defined as the average head position across the recording after excluding trials where head movements exceeded 1 cm. Data were high-pass filtered at 1 Hz, and Independent Component Analysis (ICA; 70 components, method="picard", max iterations=1000) was applied to remove biological artifacts (eye blinks, horizontal eye movements, cardiac and muscle activity), identified via automatic selection and visual inspection of component time courses and topographies. Environmental noise was additionally suppressed by removing components detected via reference channels using find_bads_ref() (threshold = 2), validated by visual inspection. The mean number of removed components was 1.96 ± 0.45 (eye movements), 1.96 ± 0.20 (cardiac), 2.50 ± 1.98 (muscle), and 7.88 ± 2.82 (environmental noise).

Following ICA, data from 275 axial gradiometers were epoched from -1.7 to 1.2 s relative to unloading and visually inspected to exclude trials contaminated by instrumental or myogenic artifacts in the -0.5 to 0.5 s window (mean ± SD dropped trials: 1.29 ± 1.40). No baseline correction was applied in subsequent MEG analyses. The mean number of retained trials per participant was 54.25 ± 1.56.

### MEG source-level analysis

#### Forward and inverse solution

To align MEG sensor and head coordinates with each participant’s MRI in a common coordinate system, we used the coregistration procedure implemented in mne.gui.coregistration(). Three fiducial landmarks (nasion, left auricle, and right auricle) were accurately identified using MRI-visible markers placed prior to MRI acquisition at the same positions as the landmarks digitized during MEG recording. These markers were clearly visible on the individual MRI head surface, allowing for an accurate coregistration.

The forward model was computed using a single-layer boundary element model (20,484 vertices across both hemispheres). A mixed source space was defined, combining a surface-based cortical source space (ico4 spacing; 5,124 vertices) and a volume source space covering the basal ganglia and cerebellum (5 mm spacing; 1,300 sources). Sources were constrained to be at least 5 mm from the inner skull surface.

For source reconstruction, ICA-cleaned data were notch-filtered at 50 and 100 Hz, then band-pass filtered using zero-phase FIR filters (‘auto’ length, ‘firwin’ design, ‘reflect_limited’ padding) in two ranges: 25-150 Hz for high-gamma analysis and 1-150 Hz for alpha- and beta-band analysis. Broad filtering ranges were used to enhance beamformer source localization and power estimation, as recommended by Brookes et al. (2008). Data were epoched and bad trials excluded as described above.

The data covariance matrix was computed using the empirical method over the -1 to 0.1 s interval relative to unloading, with rank estimated using a relative singular value threshold of 1×10⁻⁶. No noise covariance matrix was estimated, given the use of a single channel type and beamformer approach. Source localization was performed using a unit-gain linearly constrained minimum variance (LCMV) beamformer (regularization=0.1), which accurately detects both superficial and deep sources (Attal & Schwartz, 2013; Jaiswal et al., 2020) and has been used in previous BLLT MEG studies (Manyukhina et al., 2026; Ng et al., 2011, 2013a, 2013b). For analyses requiring time course extraction for burst detection, a vector LCMV beamformer was used with source orientations maximizing beta-band (13-30 Hz) power. Spatial filters were applied to each epoch individually.

#### Time-frequency analysis

Time-frequency decomposition was performed on the full epoch window (-1.7 to 1.2 s), with 0.5 s symmetric padding; power data were subsequently restricted to the analysis window of interest.

High-gamma power (60-130 Hz) was estimated using the Multitaper method (tfr_array_multitaper(); 7 cycles, time_bandwidth=4, 1 Hz resolution) on single-trial data for each LCMV source. Power was log10-transformed to normalize the distribution and averaged across 90-130 Hz to yield mean high-gamma power as a proxy for cortical excitability (Brazhnik et al., 2021; Lundqvist et al., 2016; Murthy & Fetz, 1996; Ray et al., 2008; Riehle et al., 2018), as in our adult study (Manyukhina et al., 2026).

For alpha- and beta-band analyses, the adaptive Superlets algorithm (Moca et al., 2021; https://github.com/irhum/superlets; base cycles=3, orders=1-20) was applied to compute power in the 1-50 Hz range for each source and trial. Periodic and aperiodic components were then separated using Specparam ("FOOOF"; Donoghue et al., 2020) in "fixed" mode (peak widths: 0.5-12 Hz, detection threshold=2, no minimum peak height, no limit on number of peaks, no regularization).

The model was fitted to trial-averaged spectra in the 1-50 Hz range, selected based on visual inspection of spectral densities, and subsequently applied to single-trial spectra. Periodic power was extracted in the 6-30 Hz range, with the lower boundary set at 6 Hz to account for the lower alpha peak frequency in children (Edgar et al., 2019; Hagne, 1972; Miskovic et al., 2015).

#### Correlational analysis

Correlational analyses examined the relationship between brain activity and anticipatory *Biceps brachii* inhibition by restricting inhibition events to a -70 to 75 ms window relative to unloading (see *Selection of anticipatory Biceps brachii inhibitions*). Five of 24 participants had fewer than 20 clean trials with automatically identified anticipatory inhibitions (14.60 ± 2.57 inhibition trials; 40.68 ± 8.74%), suggesting inconsistent engagement of anticipatory postural control mechanisms. Correlational analyses were therefore restricted to the remaining 19 participants (33.00 ± 9.93 inhibition trials; 71.92 ± 12.73%).

First, we aimed to validate findings from our adult study showing that stronger *Biceps brachii* inhibition was associated with reduced SMA – but not M1 – excitability (Manyukhina et al., 2026). If SMA mediates direct corticospinal control of the *Biceps brachii* during load support, as proposed earlier (Ng et al., 2011, 2013b), this negative correlation would suggest that *Biceps brachii* inhibition reflects cortical-level suppression of the region controlling this muscle.

To test this, we correlated Peak EMG Inhibition strength with high-gamma power (90-130 Hz), used as a proxy for cortical excitability (Ray et al., 2008; Ray & Maunsell, 2011; Yazdan-Shahmorad et al., 2013), across trials for each participant within SMA and M1 labels. The labels were defined as follows: the right SMA label was centered on the peak coordinate identified in adults (MNI305: [8, 3, 52]; Manyukhina et al., 2026), and the right M1 label on the elbow area reported in the literature (MNI305: [32, -20, 66]; Lotze et al., 2000; Plow et al., 2010). Each label comprised the “peak” vertex and its first- and second-order neighbours (37 vertices in total; Fig. S3). Correlations were computed at each time point from -0.1 to 0.05 s, centered on EMG Inhibition Peak Time, to capture neural effects closely aligned with maximal muscle inhibition, and across all cortical sources. A one-tailed permutation cluster test restricted to SMA and M1 labels assessed whether stronger *Biceps brachii* inhibition was significantly associated with reduced high-gamma power in these regions.

Within the SMA label, we further tested whether 6-30 Hz periodic power is associated with anticipatory muscle inhibition by examining whether it is correlated with stronger inhibition or longer inhibition duration. Correlations between periodic 6-30 Hz power and Peak EMG Inhibition or EMG Inhibition Duration were computed at each vertex and time-frequency point within -0.1 to 0.05 s around EMG Inhibition Peak Time. Correlation coefficients were averaged across vertices, and one-tailed permutation cluster tests assessed whether values were significantly less than zero (inhibition strength) or greater than zero (inhibition duration), indicating that increased alpha- or beta-band activity was associated with stronger and/or longer inhibition.

To assess whether alpha- or beta-band activity contributes to the modulation of SMA excitability associated with anticipatory inhibition, we computed, at each vertex of the SMA label, partial correlations between high-gamma power (averaged over 90-130 Hz and the time window correlated with anticipatory inhibition strength) and 6-30 Hz power within -0.1 to 0.05 s around the inhibition peak, controlling for mean 1-130 Hz power. Mean 1-130 Hz power was included as a control variable to control for residual aperiodic fluctuations, since aperiodic subtraction does not fully eliminate broadband aperiodic activity from the spectrum. Coefficients were averaged across vertices, and a one-tailed permutation cluster test assessed whether values were significantly less than zero, indicating that stronger alpha-beta activity was associated with greater high-gamma suppression.

As supplementary analyses, we correlated Peak EMG Inhibition and periodic 6-30 Hz power across right- and left-hemispheric cortical regions, right basal ganglia, and bilateral cerebellum, to test whether regions beyond SMA contribute to anticipatory inhibition via alpha- or beta-band activity. Similarly, we correlated 6-30 Hz power with EMG Inhibition Duration to assess broader network contributions to inhibition duration.

To identify regions potentially involved in the timing of *Biceps brachii* inhibition, we additionally estimated phase-locking values (PLV) in the 6-30 Hz range across the same regions, based on the assumption that regions contributing to inhibition timing should show consistent oscillatory phase across trials when aligned to inhibition onset (Bonnefond & Jensen, 2015; Samaha et al., 2015; Solís-Vivanco et al., 2018).

Methods and results for both analyses are provided in Supplementary Materials (Fig. S12).

#### GAMs on correlational effects

The observed correlational effects on anticipatory *Biceps brachii* inhibitions could be specific to the anticipatory period or persist across a broader range of inhibition timings. To clarify the temporal specificity of observed effects while accounting for subject-level variability, we employed generalized additive models (GAMs), which allow modeling of nonlinear relationships over time (Sóskuthy, 2017; Wieling, 2018). GAMs were fitted on trials with inhibition peak times between -70 and 150 ms relative to unloading in all 24 participants, capturing both anticipatory and reactive inhibitions. Each model included two smooth terms and their interaction, with random intercepts and slopes to account for subject variability. The first smooth term corresponded to the main variable from the correlation analysis, averaged over the significant time-frequency window; the second smooth term represented EMG Inhibition Peak Time. Their interaction assessed whether the relationship between the first term and the outcome was inhibition-time-dependent. To normalize variable distributions, Peak EMG Inhibition was Box-Cox transformed, and total beta power was included in the GAMs as residuals after regressing out broadband contributions to control for aperiodic activity (see Statistical Analysis).

#### Beta burst detection and selection

An increasing body of research suggests that beta-band activity in the brain occurs as transient bursts rather than sustained oscillations (Lundqvist et al., 2024). Burst-related power modulations can confound monotonic or linear estimates, producing misleading results. We therefore reexamined beta-related findings by treating beta activity as discrete, transient, and potentially functionally heterogeneous events – beta bursts – to elucidate the mechanisms underlying anticipatory muscle inhibition.

Burst detection was performed within the SMA label, where high-gamma suppression and beta power were linked to stronger anticipatory inhibition, to reduce computational costs. Bursts were identified using the pipeline of Szul et al. (2023; https://github.com/danclab/burst_detection). Power was computed using the Superlets algorithm as described above, with higher frequency resolution (0.25 Hz) to improve burst detection. Because we aimed to detect all beta bursts around the peak of anticipatory inhibition, bursts overlapping in time but not in frequency were allowed, and evoked-response correction was not applied.

Beta bursts were detected at each vertex of the SMA label within a -1 to 1 s window relative to inhibition peak time. To avoid edge effects, detection was performed over 10-33 Hz, and only bursts with peak frequencies within 13-30 Hz were retained. Burst detection was verified by visual inspection of trial-wise time-frequency representations. For each burst peak amplitude, peak time, duration, and frequency span were extracted. All bursts detected within the label were pooled and treated as individual events.

As in our previous study (Manyukhina et al., 2026), correlation analysis was used to guide the selection of bursts potentially linked to anticipatory inhibition, based on the rationale that if a specific burst type contributes to the process, its temporal dynamics and dominant frequency should show at least a modest association with the behavioural measure. Since two distinct beta-band clusters exhibited relationships with Peak EMG Inhibition with differing temporal dynamics, as indicated by GAM results (see Results, Fig. 2B), two burst types were extracted. As these clusters overlapped in time, bursts were extracted within the 80 ms preceding the inhibition peak and distinguished by frequency: "lower-beta" bursts (peak frequency 19-24 Hz) and "higher-beta" bursts (peak frequency 24-29 Hz). Note that this selection, restricted to a specific time and frequency window, means that such bursts were not necessarily identified in every trial.

**Figure 2.**
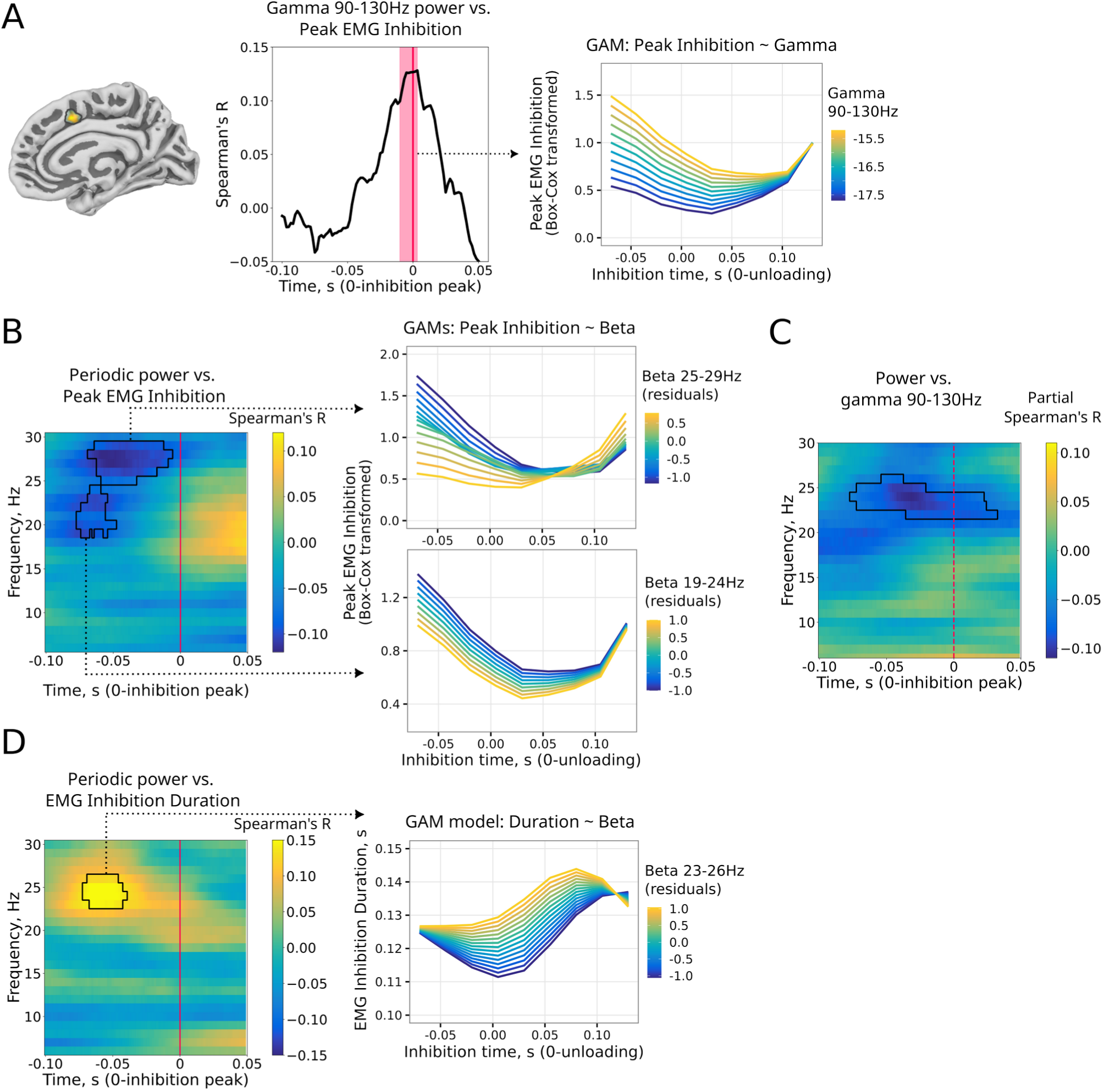
Relationships between *Biceps brachii* inhibition and SMA activity. **A:** Left panel: SMA cluster (four vertices) showing stronger high-gamma suppression (90-130 Hz) linked to greater *Biceps brachii* inhibition (Peak EMG Inhibition). Middle panel: the thick black line shows the group-averaged correlation time course within the cluster, with the red-shaded area indicating the significant time span (p < 0.05, corrected). Right panel: the GAM prediction for the same relationship, with high-gamma averaged over the cluster time window and Inhibition Peak Time included in the model. **B:** Left panel: correlation between 6-30 Hz periodic power and Peak EMG Inhibition in SMA. Right panel shows GAM predictions for the same relationship separately for the two beta clusters, with beta power averaged within each cluster and Inhibition Peak Time included in the models. **C:** Partial correlation between high-gamma power (90-130 Hz, averaged over the gamma cluster in panel A) and 6-30 Hz power, controlling for broadband 1-130 Hz power, in SMA. No GAM is shown, as beta power averaged within the cluster had no significant effect on high-gamma power. **D:** Left panel: correlation between 6-30 Hz periodic power and EMG Inhibition Duration in SMA. Right panel: the GAM prediction for the same relationship with beta power averaged within the cluster and Inhibition Peak Time included in the model. Thin black lines in panels **B-D** indicate significant correlations (p < 0.05, corrected). See Table 2 for GAM results.

**Table 2.** GAM results for smooth terms derived from SMA activity. **A:** Peak EMG Inhibition predicted from high-gamma (90-130 Hz) power and inhibition peak time (Fig. 2A). **B:** Peak EMG Inhibition predicted from beta power (19-24 Hz and 25-29 Hz) and inhibition peak time (Fig. 2B). **C:** High-gamma (90-130 Hz) power predicted from 22-26 Hz beta power and inhibition peak time (Fig. 2C). **D:** EMG Inhibition Duration predicted from 23-26 Hz beta power and inhibition peak time (Fig. 2D).

| Term | edf | Ref.df | F | p |
| --- | --- | --- | --- | --- |
| **A: Peak EMG Inhibition ~ Gamma + Inhibition peak time + Gamma x Inhibition peak time** |  |  |  |  |
| s(Gamma) | 1.00 | 1.00 | 6.26 | 0.013 * |
| s(Inh_peak_time) | 5.45 | 6.58 | 12.52 | < 0.001 *** |
| ti(Gamma, Inh_peak_time) | 1.26 | 1.49 | 1.95 | 0.212 |
| s(Subject, Gamma) | <0.001 | 23.00 | 0.00 | 0.487 |
| s(Subject, Inh_peak_time) | 6.96 | 23.00 | 4.75 | 0.387 |
| s(Subject) | 21.79 | 23.00 | 43.26 | < 0.001 *** |
| **B-1: Peak EMG Inhibition ~ low Beta + Inhibition peak time + low Beta x Inhibition peak time** |  |  |  |  |
| s(Beta_low) | 1.00 | 1.00 | 2.66 | 0.103 |
| s(Inh_peak_time) | 5.46 | 6.58 | 12.74 | < 0.001 *** |
| ti(Beta_low, Inh_peak_time) | 1.00 | 1.00 | 0.70 | 0.402 |
| s(Subject, Beta_low) | 10.93 | 23.00 | 0.86 | 0.015 * |
| s(Subject, Inh_peak_time) | 6.79 | 23.00 | 5.53 | 0.333 |
| s(Subject) | 21.97 | 23.00 | 45.35 | < 0.001 *** |
| **B-2: Peak EMG Inhibition ~ high Beta + Inhibition peak time + high Beta x Inhibition peak time** |  |  |  |  |
| s(Beta_high) | 1.00 | 1.00 | 2.99 | 0.084 |
| s(Inh_peak_time) | 5.27 | 6.38 | 12.35 | < 0.001 *** |
| ti(Beta_high, Inh_peak_time) | 5.09 | 6.98 | 2.47 | 0.017 * |
| s(Subject, Beta_high) | <0.001 | 23.00 | 0.00 | 0.886 |
| s(Subject, Inh_peak_time) | 5.55 | 23.00 | 3.06 | 0.433 |
| s(Subject) | 22.03 | 23.00 | 41.80 | < 0.001 *** |
| **C: Gamma ~ Beta + Inhibition peak time + Beta x Inhibition peak time** |  |  |  |  |
| s(Beta) | 1.000 | 1.000 | 1.196 | 0.274 |
| s(Inh_peak_time) | 1.091 | 1.175 | 0.048 | 0.871 |
| ti(Beta, Inh_peak_time) | 1.000 | 1.000 | 0.382 | 0.537 |
| s(Subject, Beta) | 0.000 | 23.000 | 0.000 | 0.861 |
| s(Subject, Inh_peak_time) | 1.962 | 23.000 | 3.174 | 0.038* |
| s(Subject) | 22.820 | 23.000 | 189.303 | < 0.001*** |
| **D: EMG Inhibition Duration ~ Beta + Inhibition peak time + Beta x Inhibition peak time** |  |  |  |  |
| s(Beta) | 1.00 | 1.00 | 3.93 | 0.048 * |
| s(Inh_peak_time) | 3.32 | 4.15 | 4.36 | 0.0016 ** |
| ti(Beta, Inh_peak_time) | 1.64 | 2.03 | 0.69 | 0.501 |
| s(Subject, Beta) | <0.001 | 23.00 | 0.00 | 0.702 |
| s(Subject, Inh_peak_time) | 5.83 | 23.00 | 1.09 | 0.231 |
| s(Subject) | 18.03 | 23.00 | 6.72 | < 0.001 *** |

ti(Gamma, Inh\_peak\_time): tensor product interaction between smooth terms; s(Subject): subject-based random intercept; s(Subject,Beta), s(Subject,Gamma): subject-specific random slope for beta
or gamma power, respectively. \* $p < 0.05$ ; \*\* $p < 0.01$ ; \*\*\* $p < 0.001$ .

#### Beta burst analyses

All subsequent burst analyses were performed separately for lower- and higher-beta bursts, with data centered on individual burst peak times. For each burst detected at a given vertex, data from that same vertex were centered on the burst peak. Trials without beta bursts (19-30 Hz) at any SMA label vertex served as a control condition. For these trials, data from all vertices were time-locked to the burst peak times observed in burst trials, to ensure comparable temporal alignment across conditions, and then averaged across vertices. Since each burst detected at each label vertex contributed a separate data sample, the total number of samples in burst analyses exceeded the number of burst-containing trials.

##### SMA label

To determine whether lower- and higher-beta bursts exhibit inhibitory properties – specifically, whether their occurrence is associated with suppression of SMA excitability as indexed by high-gamma power decrease – we aligned frequency-resolved high-gamma power (60-130 Hz) to burst peak times. A one-tailed permutation cluster test compared high-gamma power between burst and no-burst trials within -50 to 50 ms around the burst peak, to identify if bursts contribute to cortical inhibition, which should be associated with high-gamma suppression at a short temporal delay, as observed in adults (Manyukhina et al., 2026). As higher-beta bursts appeared associated with delayed high-gamma suppression, an exploratory permutation test was additionally conducted in the 0-150 ms window after higher-beta burst peaks to determine whether this burst type is significantly linked to a delayed decrease in SMA excitability.

To test whether alpha- or beta-band activity mediates the high-gamma suppression associated with each burst type, we computed partial correlations between high-gamma power (averaged over the time and frequency range of the suppression effect) and 6-30 Hz power centered on burst peak times, controlling for broadband fluctuations by regressing out mean 1-130 Hz power from the 6-30 Hz band. A one-tailed permutation cluster test within -50 to 50 ms around high-gamma suppression onset assessed whether stronger alpha- or beta-band power was associated with greater high-gamma suppression during each burst type.

To determine whether beta power increase and high-gamma suppression during each burst type were associated with anticipatory muscle inhibition strength, we computed correlations between periodic 6-30 Hz power and Peak EMG Inhibition, and between high-gamma power (averaged over 90-130 Hz) and Peak EMG Inhibition, centered on burst peak times. One-tailed permutation cluster tests within -50 to 50 ms around high-gamma suppression onset assessed whether increased alpha- or beta-band power and lower high-gamma power were associated with stronger muscle inhibition.

Finally, given the observed correlation between increased beta (23-26 Hz) power and longer muscle inhibition (Fig. 2D), we also estimated correlations between 6-30 Hz and high-gamma power with EMG Inhibition Duration during higher-beta bursts, testing whether increased alpha-beta power and greater high-gamma suppression were associated with longer inhibition.

##### Brain-level analysis

To investigate whether high-gamma suppression during each burst type reflects a local process or is supported by activity in other brain regions, we computed partial correlations between high-gamma suppression, averaged over the time and frequency of the suppression effect, and power in the frequency band that correlated with gamma suppression in the SMA label.

Correlations were estimated at each source across right cerebral cortical regions, right basal ganglia, and left cerebellum. One-tailed permutation cluster tests, performed separately for cortical and subcortical regions within the 50 ms preceding gamma suppression onset, assessed whether alpha-or beta-band power across these regions was associated with high-gamma suppression in the SMA.

##### Connectivity analysis

We further investigated the network supporting anticipatory *Biceps brachii* inhibition during voluntary unloading. Conventional connectivity approaches applied to signals with transient bursts rather than sustained oscillations may produce spurious estimates, as brief high-amplitude activity is interspersed with oscillation-free periods where connectivity cannot be meaningfully estimated (Lundqvist et al., 2024). Connectivity estimation was therefore restricted to time intervals containing beta bursts, as in our adult study (Manyukhina et al., 2026).

MEG signals were aligned to burst peak times extracted from SMA label vertices, and directed connectivity was estimated via the spectral_connectivity_epochs() function (MNE-Python). Only trials containing the selected bursts were therefore included in this analysis. Connectivity was computed between SMA vertices and selected right-hemispheric cortical regions, right basal ganglia, bilateral cerebellum, and left cortical areas to assess the potential involvement of bilateral regions in APA control, separately for the two burst types. Spectral decomposition was performed using Morlet wavelets (4 cycles). Frequency-resolved Granger causality was estimated over 3-33 Hz using 30 lags and the data rank, then restricted to 6-30 Hz to account for edge effects. As a control, Granger causality values from burst trials were compared with values computed in the opposite direction and with time-reversed (TR) Granger causality. The resulting contrast, computed separately for lower- and higher-beta bursts, was defined as follows: (A→B - B→A) - TR(A→B – B→A), where A refers to the vertices of the selected cortical and subcortical regions and B refers to the SMA vertices.

To identify regions communicating with the SMA during beta bursts, one-tailed permutation cluster tests were applied to Granger causality contrasts, separately for right- and left-hemispheric cortical regions, the right basal ganglia, and both cerebellar hemispheres. Tests focused on the 50 ms preceding burst peak within burst frequency ranges (19-24 Hz for lower-beta; 24-29 Hz for higher-beta) to capture directed influences preceding and potentially contributing to SMA bursts.

For regions identified through this analysis, Granger causality was further computed between them to assess inter-regional communication during SMA bursts, with connectivity estimated across all vertices within defined clusters.

##### Behavioural effects

Finally, we aimed to clarify which behavioral effects are associated with lower- and higher-beta bursts by contrasting trials containing each burst type with no-burst trials. As an additional EMG-derived measure, we estimated EMG Rebound, an increase in EMG amplitude ∼200 ms after unloading, closely time-locked to peak elbow rotation and thus indexing forearm instability at the muscle level. In each trial, EMG Rebound was quantified as mean EMG amplitude over a 30 ms window centered on the post-inhibition peak identified individually for each subject.

Elbow rotation and muscle inhibition measures were averaged across burst and no-burst trials for each participant and compared using paired Student’s t-tests or Wilcoxon signed-rank tests. Since beta bursts were selected based on anticipatory muscle inhibitions, no-burst trials for this comparison were also drawn from trials with anticipatory inhibitions, to avoid confounding by reactive inhibitions in EMG-based measures. These comparisons were therefore performed in 19 participants with a sufficient number of automatically defined anticipatory inhibitions (see *Selection of anticipatory Biceps brachii inhibitions* section above). For inhibition onset and duration comparisons, two additional participants with fewer than four higher-beta burst trials were excluded.

Since behavioral effects of beta bursts may be mediated by burst-associated excitability modulation (Manyukhina et al., 2026), we also assessed the relationship between burst-related high-gamma suppression and behavioral measures using GAMs in 19 participants. Individual models were fitted separately for lower- and higher-beta bursts, and for three sets of behavioral measures: (1) automatically defined EMG inhibition measures, (2) manually defined EMG inhibition measures, and (3) elbow rotation measures. Each model predicted high-gamma suppression from the corresponding behavioral measures, with random intercepts and random slopes for all fixed effects to account for subject-level variability. The first model included Peak EMG Inhibition, EMG Inhibition Peak Time, and EMG Rebound as linear fixed effects; the second included EMG Inhibition Onset and EMG Inhibition Duration; and the third included Peak Elbow Rotation and Elbow Rotation Decline (Table S4). GAM results were additionally validated using Spearman’s correlations; however, given the limited number of trials per participant, this approach is less suitable than GAMs and should be considered supplementary (Fig. S17).

### Statistical analysis

Statistical analyses were performed in Python using paired t-tests, Shapiro-Wilk tests, and Wilcoxon tests. Spearman’s rank correlations (spearmanr(), scipy.stats v. 1.7.3) were used to assess relationships between variables. For correlations involving 6-30 Hz power, periodic activity was used to reduce aperiodic contributions. For alpha-beta and high-gamma power correlations specifically, broadband power (1-130 Hz) was additionally regressed out of the 6-30 Hz band, to account for residual shared aperiodic variance that could otherwise confound power correlations (Manyukhina et al., 2026; Spaak et al., 2012).

Multiple comparisons were corrected using FDR (Benjamini-Hochberg, α = 0.05; mne.stats). For group-level analyses, individual LCMV signals were morphed to fsaverage using compute_source_morph() in MNE-Python. Non-parametric permutation cluster one-sample t-tests (5000 permutations, "hat" variance regularization) were applied to all cluster-level analyses.

Threshold-free cluster enhancement (TFCE) was applied for one- or two-dimensional data (step = 0.2); for higher-dimensional data, a cluster-forming threshold of p = 0.001 was used.

GAMs were fitted and visualized in R (v. 4.5.2), with diagnostics via gam.check() to verify residual distribution, variance homogeneity, and term smoothness. To ensure normality of variables fitted in the GAMs, a Box-Cox transformation was applied to Peak EMG Inhibition prior to model fitting (PowerTransformer(), scikit-learn v. 1.5.0). As this transformation requires non-negative values, the variable was sign-flipped before and after transformation. As periodic beta power values were non-normally distributed, total beta power was used for GAM fitting; to reduce aperiodic contributions, variance explained by broadband power (1-130 Hz) was regressed out of total beta power, and residuals were used for fitting.

## Results

### Behavioural data

#### EMG analysis

The group-averaged *Biceps brachii* EMG amplitude in the voluntary unloading condition is shown in Figure 1A, revealing a pronounced inhibition beginning prior to unloading and peaking shortly after. This suppression is followed by a transient increase in EMG amplitude around 200 ms after unloading, which we term the EMG Rebound (by analogy with the post-movement beta rebound).

Peak EMG Inhibition, expressed in arbitrary units relative to baseline, quantified inhibition strength, with more negative values reflecting stronger suppression. The inhibition peak occurred 10.82-81.67 ms after unloading (mean ± SD: 47.75 ± 20.21 ms). Inhibition onset ranged from - 39.12 to 19.64 ms (mean ± SD: -15.70 ± 13.80 ms), consistent with values reported in children (Barlaam et al., 2012; Schmitz et al., 2002). Inhibition duration ranged from 99.80 to 169.76 ms (mean ± SD: 131.40 ± 18.23 ms). Group-level distributions are shown in Figure 1A; individual-subject distributions are provided in the Supplementary Materials (Figs S4, S5).

#### Elbow rotation analysis

Figure 1B shows the group-averaged elbow rotation in the voluntary unloading condition. Following a relative flattening before unloading, elbow rotation increases progressively, quantified as Peak Elbow Rotation and expressed as a percentage of the trial-averaged Peak Elbow Rotation in the imposed condition, with lower values indicating greater forearm stability. Peak Elbow Rotation ranged from 5.9% to 36.2% (mean ± SD: 20.86 ± 8.49%), consistent with values reported in children (Barlaam et al., 2012; Schmitz et al., 2002). Although negligible in the group average, negative elbow deflection before unloading occurred in some subjects, quantified as Elbow Rotation Decline (Manyukhina et al., 2026), similarly expressed as a percentage of maximal elbow rotation in the imposed condition, ranging from -6.99% to -0.29% (mean ± SD: -1.65 ± 1.59%).

Figure 1D illustrates how EMG amplitude and elbow rotation vary with inhibition timing, classified as anticipatory (-70 to 75 ms) or reactive (75 to 150 ms): earlier muscle inhibitions were associated with better forearm stabilization and were lower in amplitude.

#### Inhibition timing and strength predict forearm stabilization

To assess relationships between elbow deflection and muscle inhibition measures, Spearman’s R correlations were estimated for each pair (Fig. 1C). Among inhibition measures, the strongest correlation was between Inhibition Onset Time and Inhibition Peak Time, as expected given that earlier onset tends to produce earlier peak, which also validates the estimates, as the two measures were derived from different approaches. Furthermore, longer inhibition was associated with earlier onset but later peak, consistent with the expected properties of muscle inhibition.

Lower Peak Elbow Rotation was associated with earlier Inhibition Peak Time, confirming that earlier inhibition is associated with better forearm stabilization (Fig. 1D). Peak Elbow Rotation also decreased over trials, likely reflecting practice-related improvement. However, forearm stabilization continues to mature over a much longer timescale across childhood and adolescence (Barlaam et al., 2012; Schmitz et al., 2002), suggesting the observed within-session change reflects only a minor contribution to this broader process. Importantly, it also suggests participants were not fatigued by the end of training, as fatigue would produce the opposite pattern. A positive relationship between Peak Elbow Rotation and Elbow Rotation Decline suggested that greater post-unloading elbow rotation tended to co-occur with a smaller pre-unloading decline, and vice versa. This relationship may reflect opposing contributions of muscle inhibition timing, with later inhibition resulting in greater peak rotation, whereas premature inhibition is associated with elbow lowering, as proposed by Manyukhina et al. (2026). However, given the absence of an association between Elbow Rotation Decline and inhibition parameters, this interpretation remains speculative.

Assessment of the relationships between the six behavioral measures and participants’ age revealed no significant correlation after FDR correction (all p’s>0.91; maximum R=0.30).

By fitting a GAM, we further examined which parameters of *Biceps brachii* inhibition had the strongest impact on forearm stabilization (Table S1; Fig. S2). Only EMG Inhibition Peak Time demonstrated a significant main effect, with earlier inhibition peaks associated with better elbow stabilization and optimal stabilization occurring around 33 ms after unloading (95% confidence interval (CI): -62 to 63 ms). A significant interaction with Peak EMG Inhibition suggested that stronger muscle inhibition during this interval further improved stabilization. Significant random intercepts and slopes indicated between-subject variability in both Peak Elbow Rotation and its relationship with Inhibition Peak Time.

### MEG data results

#### Anticipatory inhibition correlates with high-gamma suppression and beta power in the SMA

To determine whether *Biceps brachii* inhibition is associated with suppression of the SMA or M1, we estimated the correlation between Peak EMG Inhibition and high-gamma (90-130 Hz) power, as a proxy for local excitability (Brazhnik et al., 2021; Lundqvist et al., 2016; Murthy & Fetz, 1996; Ray et al., 2008; Riehle et al., 2018). A significant positive correlation was found in the medial SMA (cluster of four vertices, spanning -10 to 5 ms around inhibition peak; t_max_ = 5.29, p = 0.033; Fig. 2A), but not in M1, indicating that anticipatory *Biceps brachii* inhibition is linked to a decrease in SMA high-gamma activity, replicating prior observations in adults (Manyukhina et al., 2026). A comparison of the spatial localization of this effect with adults, along with correlation distributions across hemispheres, is provided in the Supplementary Materials (Figs. S3, S6).

In the SMA label, we further correlated alpha-beta power with Peak EMG Inhibition to test whether anticipatory muscle inhibition relates to alpha- or beta-band activity, consistent with their proposed inhibitory role. Beta power showed a significant negative correlation with Peak EMG Inhibition, spanning 19-29 Hz, -77 to -6 ms around inhibition peak (t_min_ = -3.8, p = 0.017; Fig. 2B), indicating that stronger beta power was associated with more pronounced anticipatory muscle inhibition. To test whether beta activity also linked to the SMA excitability decrease associated with anticipatory inhibition, we estimated the relationship between alpha-beta power and the high-gamma power that correlated with anticipatory inhibition. The analysis revealed a significant negative correlation in the beta band (22-26 Hz), with a cluster spanning -78 to 33 ms (t_min_ = -3.65, p = 0.015; Fig. 2C), indicating that stronger beta activity was associated with reduced high-gamma power. However, the peak frequency of this effect did not overlap with that of the beta-band correlation with anticipatory inhibition. Additionally, we assessed whether alpha- or beta-band activity predicts the duration of muscle inhibition, finding a significant positive correlation between beta power (23-26 Hz) and EMG Inhibition Duration, with a cluster spanning -72 to -39 ms (t_max_ = 3.49, p = 0.041; Fig. 2D).

As a supplementary analysis, we aimed to identify whether other cortical or subcortical regions contribute to *Biceps brachii* inhibition through alpha- or beta-band activity, by estimating the correlation between Peak EMG Inhibition and alpha-beta power across the selected brain areas. We found that stronger muscle inhibition was correlated with 14-19 Hz beta power in the right IFC (t_min_ = -6.42, p = 0.040; Fig. S12A); no correlation was observed with Inhibition Duration.

To identify regions potentially contributing to the timing of *Biceps brachii* inhibition during APA, we assessed PLV in the alpha-beta band across selected cortical and subcortical regions aligned to inhibition onset, hypothesizing that regions controlling APA timing would show phase consistency shortly before inhibition onset. Several clusters in the right hemisphere, including the SMA, PMC, M1, precuneus, basal ganglia, and cerebellum, showed a significant PLV increase in both alpha and beta ranges in the 100 ms preceding inhibition onset (Fig. S12B).

#### GAM analysis reveals temporal dynamics of inhibition timing-related neural effects

To explore the temporal dynamics of the SMA correlational effects and assess their specificity to anticipatory inhibitions, GAMs were fitted across a broader range of inhibition times in all 24 participants (Table 2; see also Supplementary Table S3 for the IFC analysis).

For high-gamma power predicting Peak EMG Inhibition, we confirmed a significant main effect of gamma, generalizing the correlation results across all participants. The largest portion of variability was explained during the anticipatory period (Fig. 2A, right panel), indicating that the effect is primarily driven by anticipatory inhibitions. Besides, a significant main effect of inhibition peak time indicated an increase in inhibition strength for later inhibitions (Fig. S7).

For beta power and inhibition strength, separate GAMs were fitted for lower- (19-24 Hz) and higher-beta (25-29 Hz) clusters (Fig. 2B). The lower-beta model showed only a tendency for a main effect of beta power (Fig. 2B, right lower panel), while significant subject-specific random intercepts and slopes indicated between-subject variability in both muscle inhibition strength and its relationship with lower-beta power. In contrast, the higher-beta model revealed a significant interaction between beta power and inhibition peak time, with beta power explaining the largest variability in inhibition strength specifically during the early anticipatory period (Fig. 2B, right upper panel). Together, these results suggest that lower- and higher-beta effects exhibit distinct inhibition-related dynamics, with only higher-beta showing a reliable link to anticipatory muscle inhibition strength.

Furthermore, the GAM confirmed a significant main effect of beta power (23-26 Hz) on EMG Inhibition Duration, with the largest variability explained during the late anticipatory period(Fig. 2D, right panel); no interaction with inhibition peak time was found. Inhibition peak time, however, showed a significant main effect on duration, with the longer inhibitions occurring during reactive inhibition trials (Fig. S8).

Finally, the GAM using 22-26 Hz beta power to predict high-gamma activity showed no significant main effect of beta or its interaction with inhibition peak time, combined with significant subject-specific random intercepts and slopes. However, a supplementary GAM predicting high-gamma power from frequency-resolved 6-30 Hz power confirmed a significant main effect in a similar frequency range (21-25 Hz), even after accounting for inhibition timing and inter-subject variability (see Supplementary Materials for details; Table S2, Fig. S9). Together, these results suggest this relationship, while likely present, is not fully robust to inter-subject variability and a broader range of inhibition times.

#### Lower- and higher-beta bursts in the SMA serve distinct functional roles

The correlational analysis confirmed the involvement of the SMA and beta-band activity in anticipatory *Biceps brachii* inhibition, replicating our findings in adults (Manyukhina et al., 2026). However, it also yielded inconclusive evidence: the frequency ranges of beta effects on Peak EMG Inhibition and on high-gamma suppression did not correspond, the link between beta and high-gamma power lacked robustness, and the two beta clusters showed different dynamics. We hypothesized that these mixed results reflect the burst-like nature of beta activity, which can bias monotonic correlation estimates toward transient high-amplitude events (Lundqvist et al., 2024), thereby obscuring the dissociation between burst occurrence and behavior-related effects.

Following our previous study (Manyukhina et al., 2026), we used the correlation clusters (Fig. 2B) to guide burst selection, reasoning that distinct peaks in the correlation time-frequency plot may reflect burst types with different time-frequency characteristics and functional contributions to anticipatory inhibition. Accordingly, we extracted two burst types corresponding to the two clusters: 19-24 Hz ("lower-beta") and 24-29 Hz ("higher-beta") bursts. Higher-beta bursts were also intended to capture the 23-26 Hz beta effect linked to inhibition duration (Fig. 2D). To avoid overlap, trials containing both burst types were therefore assigned to the higher-frequency category.

Time-frequency plots averaged across trials for each burst type are presented in Figure 3A for illustration. The average number of burst trials per subject was 21.79 ± 10.67 (lower-beta bursts) and 19.68 ± 8.37 (higher-beta bursts; mean ± SD). The distribution of anticipatory inhibition trials across categories per subject is provided in the Supplementary Materials (Fig. S10). Compared to lower-beta bursts, higher-beta bursts occurred around 10 ms earlier (lower-beta: -35 ± 11 ms relative to inhibition peak; higher-beta: -44 ± 8 ms; t = 2.27, p = 0.035), had a broader frequency span (4.36 ± 0.43 vs. 3.75 ± 0.32 Hz; W = 3, p = 1.9×10⁻⁵), and were shorter in duration (76 ± 9 vs. 85 ± 10 ms; t = -3.31, p = 0.004), with a tendency toward lower amplitude (W = 47, p = 0.055).

**Figure 3.**
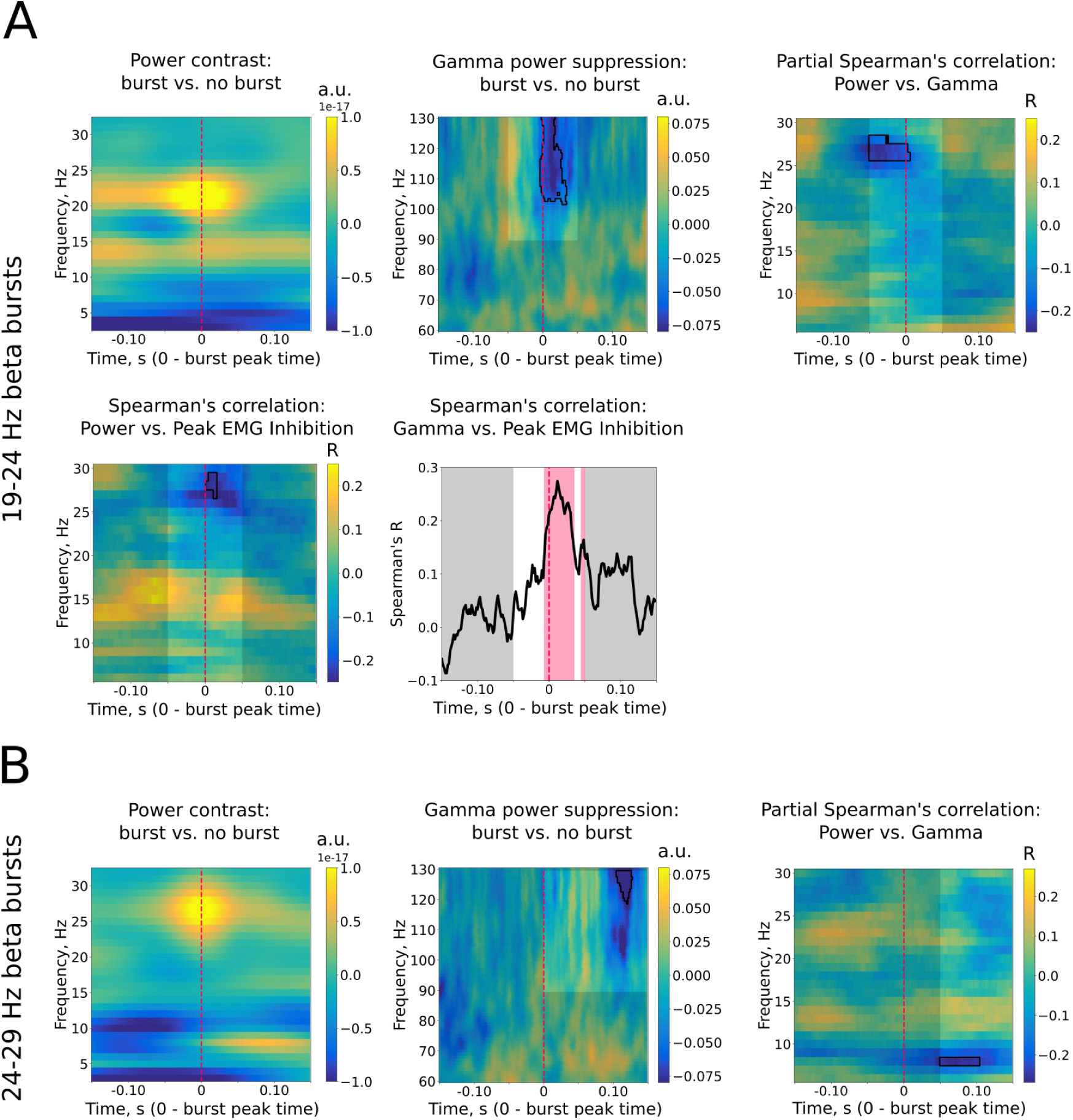
Burst analysis results in the SMA label for 19-24 Hz bursts (“lower-beta” bursts; **A**) and 24-29 Hz bursts (“higher-beta” bursts; **B**). The upper panels in **A** and **B** show analogous analyses for the two burst types: the left plot illustrates time-frequency power averaged over bursts; the middle plot shows high-gamma power suppression time-locked to burst occurrence; and the right plot depicts the correlation between high-gamma suppression (averaged over the time-frequency interval of gamma cluster) and 6-30 Hz power. The lower panels in **A** further show the correlation between *Biceps brachii* inhibition strength and 6-30 Hz power (left) and high-gamma suppression (right). No such relationships are shown for 24-29 Hz bursts as no significant correlations were observed. Bright rectangles in panels **A** and **B** indicate the analysis window, with the shaded area included for demonstration purposes. The red-shaded area in **A** highlights the significant time span, and thin black lines in **A** and **B** mark significant correlation or contrast effects (p < 0.05, corrected).

To test whether the selected burst types show inhibitory properties, we assessed whether their occurrence in the SMA was associated with a temporally proximate reduction in excitability, as reflected in high-gamma activity. The occurrence of lower-beta bursts was associated with reduced high-gamma power (103-130 Hz), with a cluster spanning -4 to 26 ms around the burst peak (t_min_ = -3.94, p_min_ = 0.024; Fig. 3A), suggesting that these bursts may reflect direct transmission of an inhibitory signal to the SMA. In contrast, higher-beta bursts showed no immediate high-gamma suppression (within 50 ms around burst peak; t_min_ = -1.54, p_min_ = 0.57), but a delayed suppression was observed (118-130 Hz), with a cluster spanning 104-128 ms after the burst peak (t_min_ = -5.08, p_min_ = 0.029; Fig. 3B), suggesting that these bursts are linked to a delayed inhibitory influence on SMA excitability.

Stronger high-gamma suppression during lower-beta bursts correlated with stronger anticipatory *Biceps brachii* inhibition, spanning -5 to 51 ms around burst peak (t_max_ = 4.04, p_min_ = 0.005; Fig. 3A), while no such relationship was found for higher-beta bursts (t_max_ = 1.27, p_min_ = 0.58) or in burst-free trials centered on burst times (t_max_ = 1.32, p_min_ = 0.58). Lower-beta bursts also showed a negative correlation between anticipatory inhibition strength and 27-29 Hz beta power, spanning 0 to 16 ms around burst peak (t_min_ = -3.53, p_min_ = 0.043; Fig. 3A), with no such effect found for higher-beta bursts (t_min_ = -2.68, p_min_ = 0.18). Altogether, these results demonstrate that the correlations between high-gamma power, beta power, and anticipatory muscle inhibition observed across all trials (see Fig. A,B) 2) are likely explained by lower-beta bursts.

Furthermore, high-gamma suppression during lower-beta bursts negatively correlated with 26-28 Hz beta power (-50 to 3 ms; t_min_ = -3.88, p_min_ = 0.029; Fig. 3A) – a frequency range that also correlated with anticipatory inhibition during these bursts, thereby establishing a three-way relationship between anticipatory inhibition, high-gamma suppression, and 26-29 Hz beta power in lower-beta burst trials. This pattern thus replicates findings in adults (Manyukhina et al., 2026), though here it is specific to trials containing a particular burst type. In contrast, high-gamma suppression during higher-beta bursts did not correlate with anticipatory inhibition strength and, similarly, showed no correlation with inhibition duration (t_min_ = -1.17, p_min_ = 0.72). However, it correlated with 8 Hz alpha power (t_min_ = -5.73, p_min_ = 0.029; Fig. 3B), with no effect in the beta range, suggesting a different underlying mechanism from lower-beta bursts.

#### Lateral PFC and PMC/M1 exhibit directed influence on the SMA during lower-beta bursts

To further clarify the functional significance of lower- and higher-beta bursts, we investigated their potential cortical and subcortical sources. To this end, we estimated Granger causality between candidate brain regions and the SMA to identify which regions exert directed influence on the SMA during bursts, using data centered on burst peaks, following our previous approach (Manyukhina et al., 2026).

For lower-beta bursts, a significant increase in directed influence toward the SMA was found in the right cortex, localized to the lateral PFC (lPFC), with a maximal cluster spanning -50 to -21 ms around burst peak across the full SMA burst frequency range (19-24 Hz; t_max_ = 4.54, p = 0.004; Fig. 4A). A trend toward increased directed connectivity from ventral PMC/M1 was also observed, with a maximal cluster spanning -38 to -16 ms and 19-22 Hz (t_max_ = 4.01, p = 0.052; Fig. 4A). Together, these two clusters replicate the brain regions found to exert directed influence on the SMA during anticipatory inhibition-related bursts in adults (Manyukhina et al., 2026). Unlike the previous study, both cerebellar hemispheres also showed directed influence on the SMA (Fig. S13). No directed influence from the right basal ganglia was found (t_max_ = 3.32, no clusters).

**Figure 4.**
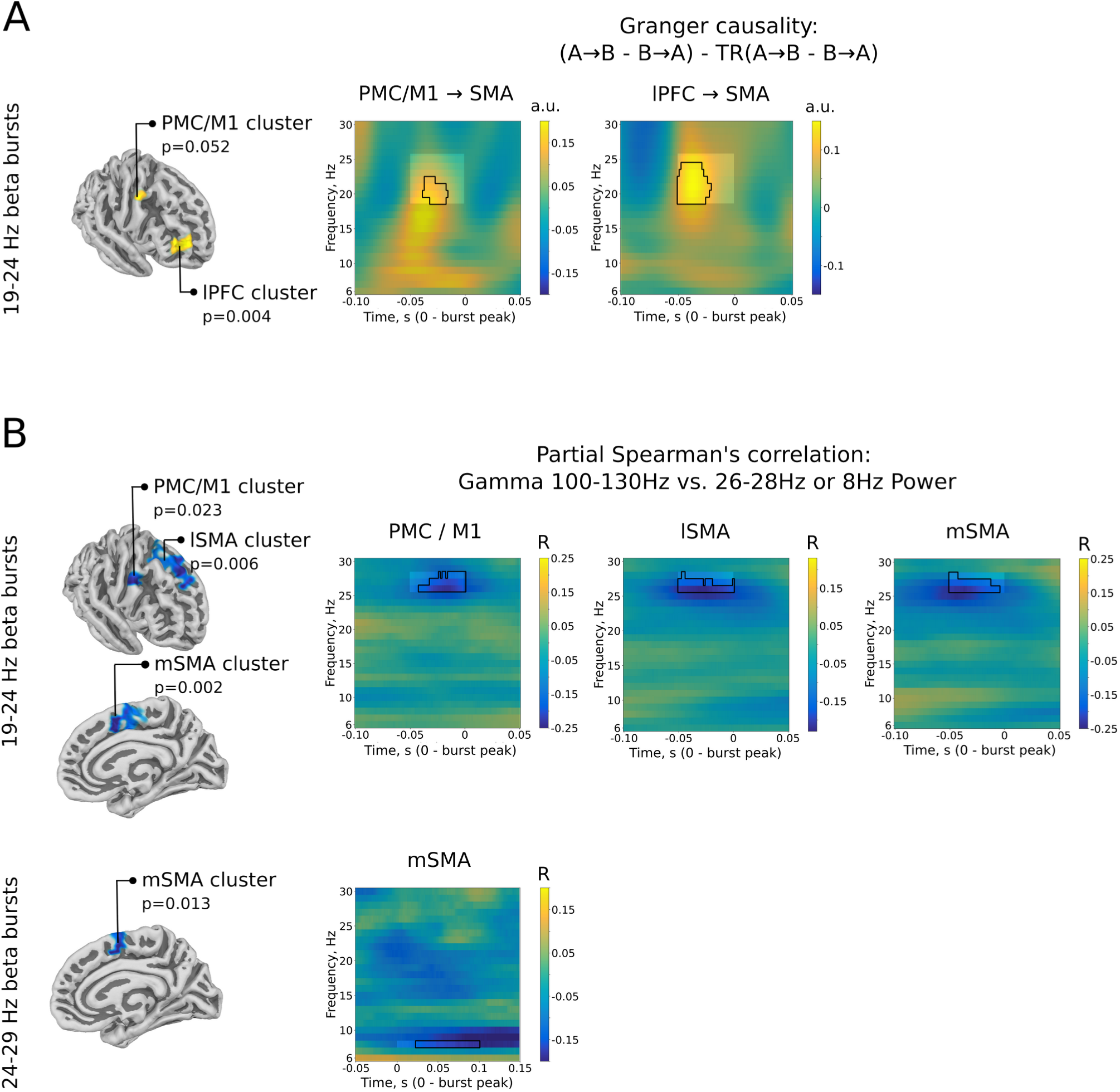
Connectivity analysis and correlations with SMA high-gamma suppression during beta bursts. **A:** Granger causality results showing right cortical regions with significant directed influence on the SMA during 19-24 Hz (“lower-beta”) bursts. No results are shown for 24-29 Hz (“higher-beta”) bursts, as no significant effects were observed. **B:** Right cortical regions showing significant negative correlations between SMA higher-gamma suppression and 26-28 Hz power for lower-beta bursts, and 8 Hz power for higher-beta bursts. Thin black lines indicate significant correlations (p < 0.05, corrected). Bright rectangles indicate the analysis window, with shaded areas included for illustration.

In contrast to lower-beta bursts, no significant directed influence on the SMA during higher-beta bursts was observed in any candidate cortical or subcortical region (right cortex: t_max_ = 3.4; left cortex: t_max_ = 3.36; right basal ganglia: t_max_ = 2.96; left cerebellum: t_max_ = 3.16; all no clusters; right cerebellum: t_max_ = 4.09, p = 0.12).

#### PMC/M1 beta power correlates with SMA high-gamma suppression during lower-beta bursts

To identify brain regions contributing to SMA excitability modulation during bursts, we correlated burst-associated SMA high-gamma suppression with the frequency-band activity that correlated with it in the SMA – 26-28 Hz beta for lower-beta bursts and 8 Hz alpha for higher-beta bursts (Fig. 3, right panels) – across selected cortical and subcortical regions.

For lower-beta bursts, three clusters of significant negative correlation were identified (Fig. 4B): the largest in the lateral SMA (-50 to 0 ms; t_min_ = -6.20, p = 0.006), a second in the medial SMA (-50 to -5 ms; t_min_ = -5.64, p = 0.002), partially overlapping the original SMA label, and a third in ventral PMC/M1 (-50 to 0 ms; t_min_ = -5.63, p = 0.023), indicating that stronger high-gamma suppression is associated with increased beta activity in these regions. No significant correlations were observed in the left cerebellum (t_min_ = -3.03, no clusters) or right basal ganglia (t_min_ = -3.23, no clusters). Notably, among these regions, only PMC/M1 also showed significant directed connectivity to the SMA during this burst type (Fig. 4A).

For higher-beta bursts, a significant negative correlation between SMA high-gamma suppression and 8 Hz alpha power was localized to the medial SMA adjacent to the original SMA label (23-100 ms after burst peak; t_min_ = -5.77, p = 0.013; Fig. 4B), indicating that stronger high-gamma suppression is associated with increased alpha activity in the surrounding region. No significant correlations were found in the right basal ganglia, though a trend was observed in the left cerebellum, with a maximal cluster spanning 3-42 ms after burst peak (t_min_ = -4.85, p = 0.059).

#### Lower- and higher-beta bursts have distinct effects on anticipatory inhibition and forearm stabilization

To clarify the behavioural effects associated with lower- and higher-beta bursts, we examined their effects on EMG- and elbow rotation-derived behavioural measures.

First, we contrasted burst and no-burst trials on EMG- and elbow rotation-derived measures.

Lower-beta burst trials showed stronger anticipatory inhibition, reflected in more negative Peak EMG Inhibition (t(18) = -2.52, p = 0.021; Fig. S17), with no differences in inhibition timing, duration, EMG rebound, or elbow rotation measures (all p’s > 0.18). In contrast, higher-beta burst trials were characterized by earlier inhibition onset (t(16) = -2.27, p = 0.037) and longer duration (t(16) = 3.12, p = 0.007; Fig. S17), with no differences in inhibition strength, peak time, EMG rebound, or elbow rotation parameters.

Second, considering that burst effects may be mediated by burst-associated excitability modulation (Manyukhina et al., 2026), we assessed whether burst-associated high-gamma suppression is linked to behavioral measures using GAMs. Stronger high-gamma suppression during lower-beta bursts was significantly predicted only by more pronounced anticipatory inhibition strength (t = 1.98, p = 0.048). In contrast, stronger high-gamma suppression during higher-beta bursts was associated with earlier inhibition onset (t = 2.15, p = 0.032), lower EMG Rebound (t = 2.85, p = 0.005), and decreased Peak Elbow Rotation (t = 2.80, p = 0.005). Full model details are provided in the Supplementary Materials (Table S4, Fig. S16).

Figure 5 summarizes the core findings of this study on the functional effects of two beta burst types and proposes two hypothetical mechanisms, indexed by lower- and higher-beta bursts (19-24 Hz and 24-29 Hz, respectively), through which the SMA supports anticipatory postural control in children.

**Figure 5.**
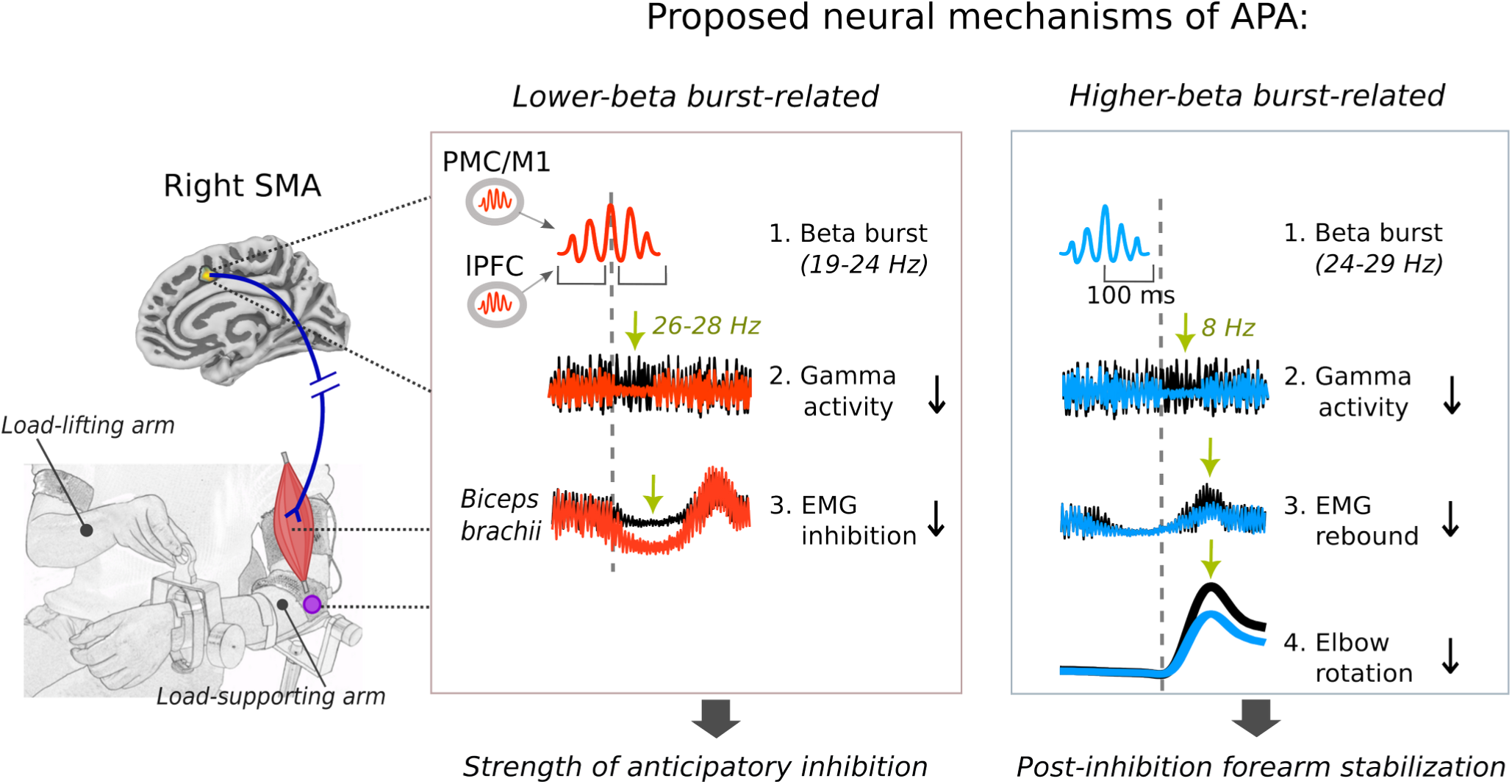
The role of beta bursts in anticipatory postural control: summary of findings and proposed models. This schematic summarizes the main findings on two beta burst types differently linked to anticipatory postural control, establishing a link between SMA burst occurrence, high-gamma suppression, *Biceps brachii* inhibition, and elbow rotation, and proposes two complementary mechanisms associated with each burst type. First, a lower-beta burst (19-24 Hz) in the contralateral SMA (1) reduces local excitability, as reflected in high-gamma (90-130 Hz) suppression, mediated by 26-28-Hz activity (green arrow) (2), resulting in *Biceps brachii* inhibition reflected in EMG amplitude decrease (green arrow) (3), thereby replicating the mechanism proposed for anticipatory inhibition in adults (Manyukhina et al., 2026). Second, a higher-beta burst (24-29 Hz) (1) does not produce immediate SMA suppression but is time-locked to a delayed (∼100 ms) high-gamma suppression mediated by 8-Hz activity (green arrow) (2), which attenuates post-inhibition increases in EMG (3) and elbow rotation (4) (green arrows), thereby compensating for unloading-induced forearm instability. Notably, this mechanism was not observed in adults, suggesting it is recruited during development to compensate for imprecise anticipatory inhibition. Together, the two mechanisms may ensure forearm stabilization during voluntary load-lifting in children: one by supporting anticipatory inhibition and the other by additionally compensating for the postural disturbance arising from imprecise anticipatory postural control.

## Discussion

To our knowledge, this study provides the first characterization of the neurodynamical mechanisms underlying anticipatory muscle inhibition – a core component of APA – in children. Using MEG during voluntary bimanual load-lifting, we investigated the neural basis of anticipatory *Biceps brachii* inhibition during ongoing APA maturation, to determine whether the underlying mechanisms are similar to or distinct from those in adults. A core finding is the replication of core results from adults (Manyukhina et al., 2026): stronger anticipatory muscle inhibition was associated with reduced high-gamma activity (90-130 Hz) in the SMA rather than M1, time-locked to 19-24 Hz beta bursts, and correlated with increased beta power. These effects were accompanied by a similar pattern in connectivity: SMA beta bursts were preceded by directed influences from lateral PFC and ventral PMC/M1, while only PMC/M1 beta activity correlated with the magnitude of SMA high-gamma suppression. Importantly, children additionally exhibited a second beta burst type not observed in adults, associated with delayed (∼100 ms) SMA high-gamma suppression mediated by 8 Hz alpha activity and attenuation of post-inhibition EMG and elbow rotation increases, thereby ensuring forearm stabilization. The two burst types thus likely reflect distinct functional components of the APA mechanism: the lower-frequency (19-24 Hz) bursts supporting anticipatory muscle inhibition, and the higher-frequency (24-29 Hz) bursts compensating for the postural disturbance arising from its imprecision, together providing mechanistic insight into how the developing brain proactively supports and compensates for immature anticipatory postural control.

### Forearm stabilization is predicted by the timing and strength of anticipatory muscle inhibition

In the BLLT, anticipatory inhibition of the postural forearm flexors counteracts the upward elbow rotation caused by unloading, ensuring postural stability. Previous BLLT studies proposed that inhibition timing is a crucial determinant of APA efficiency, with inhibition time-locked to unloading onset associated with better stabilization (Hugon et al., 1982; Massion, 1992) – a relationship confirmed in the present study (Fig. 1C, D). The substantial variability in inhibition timing in children observed here (Fig. 1A) and in previous studies (Barlaam et al., 2012; Schmitz et al., 2002) may thus account for their less efficient forearm stabilization relative to adults: elbow rotation during voluntary unloading reached 21% of that in the imposed condition, consistent with earlier reports in children of similar age (≈20%: Barlaam et al., 2012; Schmitz et al., 2002) – markedly higher than in adults (5-8%: Barlaam et al., 2012; Manyukhina et al., 2026). However, whether inhibition onset, peak time, or other inhibition parameters such as strength and duration are the primary determinants of forearm stabilization efficiency was not directly addressed.

We found that among the inhibition parameters examined, only inhibition peak time significantly predicted maximal elbow rotation, indicating that efficient forearm stabilization primarily depends on when muscle inhibition peaks during the anticipatory period. A significant interaction effect further revealed that forearm stabilization was most effective when stronger muscle inhibition peaked during the late anticipatory period, approximately 0-50 ms after unloading. This result is consistent with adult findings, where forearm stabilization similarly depended on inhibition strength during the anticipatory period, though the optimal interval preceded unloading onset (Manyukhina et al., 2026). The later optimal window in 7–12-year-old children may reflect an adaptive shift to greater task demands: in adults, earlier inhibition onset and later peak were observed lying (Ng et al., 2011) versus sitting (Dufossé et al., 1985; Hugon et al., 1982; Viallet et al., 1992), suggesting that optimal inhibition timing adapts to postural demands – a question that remains to be directly addressed in future studies.

Overall, the results of this analysis showed that the strength of anticipatory inhibition is a primary determinant of APA efficiency, which therefore became the focus of subsequent analyses.

### SMA beta activity mediates anticipatory postural muscle inhibition

Previous investigations of APA mechanisms during the BLLT have yielded mixed findings.

It was initially proposed that anticipatory muscle inhibition reflects suppressed activity in the cortical region controlling the muscle – specifically M1 (Kazennikov et al., 2005) – consistent with lesion and EEG studies implicating M1 in APA (Barlaam et al., 2011, 2018; Martineau et al., 2004; Viallet et al., 1992). However, TMS and MEG studies failed to find APA-specific M1 involvement (Kazennikov et al., 2005; Ng et al., 2011, 2013b), leading to the alternative proposal that the SMA, rather than M1, mediates postural muscle control during APA (Ng et al., 2011, 2013b). This interpretation was supported by our recent study in adults (Manyukhina et al., 2026), in which stronger anticipatory *Biceps brachii* inhibition was associated with reduced high-gamma power (90-130 Hz) in the medial SMA but not M1 – a finding we replicate here in children using a more sensitive automated approach to quantify inhibition strength on a single-trial basis. In both studies, we used high-gamma (90-130 Hz) power as an index of regional excitability, following studies linking broadband high-gamma activity to neuronal spiking and local excitability (Brazhnik et al., 2021; Lundqvist et al., 2016; Ray et al., 2008; Riehle et al., 2018; Sohal & Rubenstein, 2019).

While high-gamma activity is susceptible to myogenic artifacts (Muthukumaraswamy, 2013; Whitham et al., 2008), such artifacts are unlikely to account for the observed gamma suppression effect, as stronger muscle activity would increase rather than decrease high-gamma power.

Additional artifact controls, including ICA-based removal of muscle components and beamforming-based source localization, further minimized potential muscle contributions.

The medial SMA has been consistently implicated in APA during the BLLT (Barlaam et al., 2011; Ng et al., 2013a, 2013b; Viallet et al., 1992), with contralateral medial frontal lesions selectively impairing APA without affecting postural control per se (Massion, 1992; Viallet et al., 1992). While it was proposed that the SMA contributes to APA by gating postural circuits (Viallet et al., 1992), our findings clarify its role, suggesting that the SMA may instead directly regulate postural muscle activity during anticipatory motor control. Although muscle control is classically attributed to M1 corticospinal projections, converging evidence from monkey and human studies demonstrates that somatotopically organized corticospinal pathways also originate from the SMA, directly mediating muscle-specific activations with latencies and amplitudes comparable to those of M1 (Dum & Strick, 1991; Entakli et al., 2014; Teitti et al., 2008). While evidence remains limited, these SMA-originating projections have been linked to tasks requiring precise force regulation and anticipation of upcoming task demands (Chen et al., 2013; Spieser et al., 2013), consistent with the demands of the BLLT. In this context, our findings offer novel evidence for a specific role of the SMA in regulating muscle activity under conditions requiring predictive motor control. This role appears particularly relevant during APA, where the SMA may mediate anticipatory control of tonic postural muscle activity.

Anticipatory *Biceps brachii* inhibition was also associated with increased beta- rather than alpha-band activity in the SMA, replicating previous results in adults (Manyukhina et al., 2026) and suggesting that inhibition of ongoing tonic muscle activity is mediated by beta-band dynamics, consistent with evidence linking beta activity to sustained muscle contractions (Bräcklein et al., 2022; Echeverria-Altuna et al., 2022; Graef et al., 2026). However, this beta activity did not correlate with concurrent high-gamma suppression, which instead emerged later and within a narrower frequency range (Fig. 2), in contrast to adults (Manyukhina et al., 2026), suggesting that this dissociation may reflect complex dynamics associated with beta bursts (Lundqvist et al., 2024). To disentangle burst-related effects, we separately analyzed trials with beta bursts in two frequency clusters: lower-beta (19-24 Hz) and higher-beta (24-29 Hz), which showed distinct temporal dynamics in their relationship with anticipatory inhibition (Fig. 2B).

### Lower-beta bursts are linked to muscle inhibition and supported by lPFC and PMC/M1

We observed that only lower-beta bursts were associated with high-gamma (90-130 Hz) suppression time-locked to the second half of the burst (Fig. 3A), consistent with findings in adults (Manyukhina et al., 2026). This burst-locked suppression correlated with anticipatory muscle inhibition strength, suggesting that lower-beta bursts primarily account for the relationship between high-gamma power and muscle inhibition observed across all trials (Fig. 2A). Notably, both high-gamma suppression and inhibition strength during these burst trials correlated with beta power in the 26-29 Hz range, thereby replicating the anticipatory inhibition–high-gamma–beta relationships previously observed in adults (Manyukhina et al., 2026), suggesting that these bursts are functionally analogous to those found in that study. This interpretation is further supported by replication of the directed connectivity pattern associated with these bursts (Manyukhina et al., 2026; Supplementary Materials): the lPFC and ventral PMC/M1 exerted directed influence on the SMA time-locked to the first half of the burst. Together, these findings suggest that adult-like neural mechanisms for anticipatory inhibition are already established by middle childhood.

One difference from adults was a mismatch between burst frequency (19-24 Hz) and the beta frequency correlating with high-gamma suppression and anticipatory inhibition (26-29 Hz). This may reflect a reduced capacity for anticipatory inhibition-related SMA synchronization at higher beta frequencies in children, consistent with the well-documented age-related acceleration of neural rhythms across development in both the alpha (Miskovic et al., 2015; Smith, 1941; Stroganova et al., 1999) and beta band (Rayson et al., 2023; Wilkinson et al., 2024). Such reduced entrainment at higher beta frequencies may in turn compromise burst efficacy, resulting in less effective anticipatory muscle inhibition in children. This discrepancy may also explain both the presence of two beta clusters in the correlation with anticipatory inhibition (Fig. 2B) and the GAM results for the same relationship (Table 2B), which showed that only the higher-frequency cluster (25-29 Hz) remained significant after accounting for inter-individual variability, suggesting that the apparent 19-24 Hz effect in Figure 2B is driven by lower-beta (19-24 Hz) bursts time-locked to the higher-frequency oscillation. In turn, the relationship between high-gamma power and beta (22-26 Hz) activity (Fig. 2C) may thus reflect the dependence of gamma suppression on lower-beta (19-24 Hz) burst occurrence (Fig. 3A) rather than beta power per se.

Further insight into the functional role of these bursts comes from brain-level analyses.

High-gamma suppression in the SMA correlated with 26-28 Hz beta not only in the medial SMA, but also in the lateral SMA and ventral PMC/M1 (Fig. 4B), suggesting that these regions may contribute to SMA suppression during lower-beta bursts. Importantly, ventral PMC/M1 also showed a trend toward directed influence on the SMA during the same interval (p = 0.052), raising the possibility that PMC/M1 drives the 26-28 Hz beta effect in the SMA. Although the directed influence appeared only at lower-beta frequencies (19-22 Hz) with a peak below the SMA burst frequency range (Fig. 4A) and was not observed at higher frequencies correlating with SMA suppression (26-28 Hz), the two effects overlapped in time, suggesting they may reflect a single underlying process in which a greater presence of higher-frequency activity is associated with stronger inhibitory effects on the SMA.

Beyond PMC/M1, the lPFC also exerted significant directed influence on the SMA during lower-beta (19-24 Hz) bursts (Fig. 4A). As in adults, the frequency of this lPFC effect more closely matched SMA burst frequency than that of PMC/M1 (Manyukhina et al., 2026; Supplementary Materials), suggesting that SMA synchronization in the higher-beta range may be partly constrained by lPFC input. One notable difference from adults was the absence of directed lPFC-to-PMC/M1 influence during SMA bursts, with connectivity tending toward the opposite direction (Fig. S14).

This suggests reduced prefrontal control over premotor areas in children, potentially reflecting the continued maturation of prefrontal circuitry through adolescence (Barnea-Goraly et al., 2005; Gogtay et al., 2004; Sowell et al., 2004), which may further limit the efficacy of bursts underlying anticipatory inhibition.

Although the specific contributions of lPFC and PMC/M1 to SMA bursts and anticipatory inhibition remain to be clarified, their roles likely differ. The lPFC (Brodmann area 46) has been consistently implicated in response inhibition tasks, modulation of inhibitory tone according to task demands, and task-rule selection (Khan et al., 2024; McNab et al., 2008; Rubia et al., 2001; Zheng et al., 2008), with beta synchrony in lPFC reflecting engagement of task-relevant ensembles (Buschman et al., 2012). During APA, the lPFC may thus contribute to selecting and activating muscle-specific SMA ensembles associated with load support, thereby shaping SMA burst characteristics; suppression of these ensembles in anticipation of unloading may then facilitate a smooth transition to the unloaded state.

In contrast to lPFC, beta activity in the PMC/M1 correlated with SMA high-gamma suppression, suggesting that this region contributes to the strength of inhibitory drive onto the SMA. Although the spatial resolution of MEG does not allow disentangling the ventral PMC (vPMC) from M1, the vPMC is the more plausible source, given that connectivity typically runs from the SMA to M1 rather than in the reverse direction (Arai et al., 2012; Pagge et al., 2024) and that the vPMC has previously been implicated in APA (Ng et al., 2013a). The vPMC is involved in muscle-specific hand control and in encoding object-related manipulations (Bonini et al., 2014; Davare et al., 2009; Gallese et al., 1996), suggesting that it may maintain internal representations of manipulated objects. During the BLLT, the vPMC may therefore use internal representations of the load to guide suppression of lPFC-preactivated SMA ensembles, thereby contributing to postural stability upon load removal.

The replication of concurrent lPFC and PMC/M1 influences on the SMA across both adults and children suggests that their cooperative input may be essential for SMA burst generation and smooth postural transitions following load release. How concurrent inputs at different frequencies induce bursts with specific characteristics in the receiving region remains to be clarified, and will likely require novel computational models of beta burst generation. A potential cerebellar contribution to anticipatory inhibition bursts, observed in children (Fig. S13) but not adults, also warrants further investigation.

### Higher-beta bursts are linked to post-inhibition forearm stabilization

Trials containing higher-beta bursts (24-29 Hz), despite falling within the frequency range linked to inhibition strength during lower-beta bursts (27-29 Hz), did not share the same properties. They occurred on average 10 ms earlier than lower-beta bursts, with a broader frequency span and shorter duration. Importantly, they were not associated with high-gamma suppression time-locked to the burst peak, but rather to a delayed (∼100 ms) suppression (Fig. 3B), likely reflecting an indirect relationship. Neither high-gamma suppression nor power modulation during these bursts correlated with anticipatory muscle inhibition strength. Stronger high-gamma suppression during higher-beta (24-29 Hz) bursts was associated with increased 8 Hz alpha activity, visible in trial-averaged power during this burst type (Fig. 3B), suggesting a delayed, alpha-dependent suppression mechanism distinct from the beta-mediated inhibition during lower-beta (19-24 Hz) bursts. The source of this 8 Hz activity remains unclear: as this relationship was confined only to the SMA in the brain-level analysis (Fig. 4B), alpha activity may be locally generated; alternatively, higher-beta bursts may engage subcortical regions, less visible to MEG, which in turn give rise to an inhibitory alpha response in the SMA. Such closed-loop interactions are particularly characteristic of the cerebellum (Kang et al., 2021; Purves et al., 2018). Notably, a similar correlation between SMA high-gamma suppression and ∼9 Hz alpha activity was previously observed in adults (Manyukhina et al., 2026; Supplementary Materials), which also did not correlate with anticipatory inhibition strength; the present findings thus provide a potential explanation for this effect.

In contrast to lower-beta (19-24 Hz) bursts, no regions showed significant directed influence on the SMA preceding higher-beta (24-29 Hz) bursts. This lack of connectivity may reflect either the presence of cross-frequency interactions to which Granger causality is insensitive (Bressler & Seth, 2011), 2011), dependence of higher-beta bursts on non-oscillatory inputs, as proposed in a model of burst generation (Sherman et al., 2016), or their local generation within the SMA. Indeed, a recent study distinguished local bursts, confined to cortex, from global bursts synchronizing across cortical and subcortical networks (Khanna et al., 2025). Within this framework, higher-beta (24-29 Hz) bursts may reflect local SMA dynamics, supported by the confinement of the high-gamma–alpha relationship to the SMA (Fig. 4B). Lower-beta bursts, by contrast, showed reproducible connectivity profiles across two independent datasets, consistent with global burst properties. Notably, global bursts have been associated with inhibitory effects on neuronal firing and movement slowing (Khanna et al., 2025), in line with their inhibitory role reported here.

While the link between lower-beta (19-24 Hz) bursts and anticipatory inhibition replicated adult findings (Manyukhina et al., 2026), the presence of a functionally distinct higher-beta (24-29 Hz) burst type, not associated with anticipatory inhibition, is a novel finding, raising questions about its functional role. Although higher-beta bursts were not associated with anticipatory inhibition strength, they were linked to earlier inhibition onset and longer duration, potentially explaining the beta-inhibition duration relationship across all trials (Fig. 2D). Furthermore, stronger high-gamma suppression during these bursts correlated with earlier inhibition onset, lower EMG Rebound, and lower Peak Elbow Rotation. As this high-gamma suppression occurred on average 55 ms after EMG inhibition peak, its link to inhibition onset is likely indirect; instead, this timing aligns with the post-inhibition rise in EMG and elbow rotation – reflected in EMG Rebound and Peak Elbow Rotation – suggesting that SMA suppression during higher-beta bursts may attenuate this raising phase, thereby contributing to forearm stabilization. Since higher-beta bursts arise no later than lower-beta bursts, they are unlikely to reflect a feedback response to inefficient inhibition, and instead may represent a distinct APA mechanism engaged to compensate for predicted forearm instability. These bursts may be driven by higher-level processes that predict APA errors in parallel with anticipatory inhibition commands and proactively engage a compensatory mechanism, though the specific nature of this signal remains to be clarified. We propose that higher-beta bursts thus constitute a second, complementary APA mechanism that compensates for immature anticipatory inhibition through a distinct, alpha-mediated process, thereby ensuring more effective postural control. Together, these two burst-associated mechanisms may contribute to more efficient APA during development. This interpretation aligns with evidence linking beta activity to prediction and improved motor performance (Arnal et al., 2015; Van Pelt et al., 2016), while extending it by suggesting that beta bursts may not only carry predictive signals but also index proactive corrective mechanisms that support predictive motor control.

An open question is why the brain implements alpha- rather than beta-mediated inhibition for post-inhibition forearm stabilization. Since alpha oscillations mediate cortical excitability across visual and sensorimotor networks (Bonnefond & Jensen, 2015; Haegens et al., 2011; Kelley et al., 2025), an 8 Hz alpha increase may establish a broad inhibitory influence over SMA area controlling postural muscles, globally suppressing information gating (Jensen & Mazaheri, 2010; Klimesch et al., 2007) and thereby cancelling residual muscle activation – a mechanism well-suited to complement the precise, muscle-specific inhibition mediated by beta-band activity.

### Implications for APA maturation and its prolonged development

The replication of adult-like mechanisms for anticipatory inhibition in children aged 7-12 years indicates that the core neural pathways and processes are already in place by middle childhood, although they remain far from adult levels of efficiency. The presence of an additional anticipatory compensatory mechanism suggests that the developing brain actively mitigates its own limitations – a process that may also exist in adults but is likely less prominent during self-initiated unloading due to more efficient APA. This raises the question: what accounts for the persistent imprecision of APA throughout childhood and adolescence, reflected in substantial variability in anticipatory inhibition timing and poorer postural stability extending into early adulthood (Barlaam et al., 2012)?

Several answers can be proposed. One potential explanation lies in ongoing musculoskeletal development, which continuously alters body proportions and force requirements, thereby demanding constant motor re-adaptation (Barlaam et al., 2012; Schmitz et al., 2002). Consistent with this, we observed no improvement in forearm stabilization or inhibition parameters across the 7-12 year age range, likely reflecting the non-linear trajectory of APA maturation driven by the interplay between age and changing body parameters. Further evidence is provided by the association between anticipatory inhibition strength and IFC beta activity (Fig. S12A) – a relationship specific to children, as it was not observed in adults (Manyukhina et al., 2026). IFC has been associated with response inhibition (Suda et al., 2020), updating sensorimotor representations during unpredictable load-lifting (Schmitz et al., 2005), and error processing (Aron et al., 2003), suggesting that in children, IFC may modulate inhibitory tone according to the predicted consequences of unloading, a process that likely becomes more automated and thus less IFC-dependent as APA matures.

At the neurodynamic level, APA inefficiency may reflect immature transient beta dynamics, particularly evident in the mismatch between SMA burst frequency and the frequency of the associated high-beta effect (Fig. 3A), suggesting reduced neuronal entrainment in the high-beta range in the SMA, potentially linked to the ongoing maturation of lPFC pathways discussed above.

The mismatch with higher-frequency PMC/M1 input may in turn affect SMA burst characteristics, reducing the precision and strength of anticipatory inhibition. The lack of lPFC-to-PMC/M1 directed influence in children further implicates immature prefrontal circuitry as a potential source of APA inefficiency.

Finally, although not the focus of this study, imprecise APA timing may reflect inefficient temporal alignment between the voluntary motor and the anticipatory postural commands, potentially arising from weaker coupling within relevant motor networks or immature interhemispheric communication. Although our data showed no evidence of ipsilateral hemispheric involvement in anticipatory inhibition across correlation, connectivity, or phase-locking analyses (Figs. 4, S12), consistent with lesion evidence showing that contralateral, but not ipsilateral, damage disrupts APA (Massion, 1992; Viallet et al., 1992), APA occurs exclusively in the context of voluntary movement, making efficient interhemispheric coordination a plausible contributor to timing variability. The cerebellum plays a well-established role in timing neural processes (Ivry & Keele, 1989; Salman, 2002) and has been proposed as a key relay transmitting movement-related signals to the contralateral hemisphere to enable APA (Ng et al., 2013b); its protracted maturation through middle and late childhood (Tiemeier et al., 2010) therefore makes it a plausible source of inhibition timing variability in children. Consistent with this, we observed cerebellar involvement in both connectivity analysis and phase-locking to inhibition onset (Figs. S12, S13), supporting a potential role in APA timing and suggesting that that cerebellar-mediated interhemispheric communication may contribute to timing variability in children.

Several limitations should be noted. First, variability in inhibition timing resulted in a limited number of anticipatory inhibitions per participant and burst type, constraining trial-level analyses. Second, the limited sensitivity of MEG to deep sources (Attal & Schwartz, 2013) means that subcortical contributions, particularly from the basal ganglia, may have gone undetected, and cerebellar effects should be interpreted with caution. Third, burst selection based on correlation analyses, though yielding replicated results across two independent datasets (Manyukhina et al., 2026), remains suboptimal and may not fully separate distinct burst-related effects, highlighting the need for methods linking burst dynamics directly to behaviour. Finally, Granger causality is inherently limited to linear information transfer (Bressler & Seth, 2011) and therefore cannot capture cross-frequency burst communication; developing reliable measures for such communication remains a critical gap in understanding transient neural interactions.

In conclusion, this is the first neuroimaging study to provide a mechanistic characterization of anticipatory postural control in children, replicating a central role for SMA and beta bursts established in adults alongside a children-specific contribution of the right IFC. Crucially, we demonstrate that the developing brain does not merely respond to postural disturbance arising from imprecise anticipatory control, but proactively recruits a complementary alpha-mediated compensatory mechanisms. Burst-based analysis was key to revealing the two distinct APA mechanisms, indexed by lower- and higher- (19-24 Hz and 24-29 Hz) beta bursts, serving complementary components of anticipatory postural control in children – mechanisms obscured by conventional power analyses. Together, these findings establish that core adult-like mechanisms of anticipatory postural control are already in place by middle childhood, while persistent developmental limitations may reflect ongoing musculoskeletal growth and maturation of prefrontal circuits and cortico-cerebellar coordination. More broadly, this work advances our understanding of the developmental mechanisms underlying predictive motor control and provides a framework for investigating their alteration in neurodevelopmental conditions. It further demonstrates that burst-based analyses can reveal temporally overlapping yet functionally distinct neural mechanisms invisible to conventional spectral approaches, underscoring their value for probing the dynamics of brain computation.

## Acknowledgments

The authors thank all the participants and their families.

## Author contributions

Viktoriya Manyukhina (Data curation, Formal analysis, Methodology, Software, Validation, Visualization, Writing—original draft, Writing—review & editing), Fanny Barlaam (Data curation, Investigation, Writing—review & editing), Judith Vergne (Data curation, Investigation, Writing—review & editing), Anaëlle Bain (Data curation, Investigation, Writing—review & editing), Oussama Abdoun (Methodology, Validation, Writing—review & editing), Sébastien Daligault (Data curation, Investigation, Writing—review & editing), Claude Delpuech (Methodology, Resources, Writing—review & editing), Karim Jerbi (Conceptualization, Writing—review & editing), Sandrine Sonié (Investigation, Writing—review & editing), Mathilde Bonnefond (Conceptualization, Methodology, Project administration, Supervision, Writing—review & editing), and Christina Schmitz (Conceptualization, Funding acquisition, Methodology, Project administration, Supervision, Writing—review & editing)

## References

1. Arai, N., Lu, M.-K., Ugawa, Y., & Ziemann, U. (2012). Effective connectivity between human supplementary motor area and primary motor cortex: A paired-coil TMS study. Experimental Brain Research, 220(1), 79–87. 10.1007/s00221-012-3117-5

2. Arnal, L. H., Doelling, K. B., & Poeppel, D. (2015). Delta–Beta Coupled Oscillations Underlie Temporal Prediction Accuracy. Cerebral Cortex, 25(9), 3077–3085. 10.1093/cercor/bhu103

3. Aron, A. R., Fletcher, P. C., Bullmore, E. T., Sahakian, B. J., & Robbins, T. W. (2003). Stop-signal inhibition disrupted by damage to right inferior frontal gyrus in humans. Nature Neuroscience, 6(2), 115–116. 10.1038/nn1003

4. Attal, Y., & Schwartz, D. (2013). Assessment of Subcortical Source Localization Using Deep Brain Activity Imaging Model with Minimum Norm Operators: A MEG Study. PLoS ONE, 8(3), e59856. 10.1371/journal.pone.0059856

5. Barlaam, F., Descoins, M., Bertrand, O., Hasbroucq, T., Vidal, F., Assaiante, C., & Schmitz, C. (2011). Time–Frequency and ERP Analyses of EEG to Characterize Anticipatory Postural Adjustments in a Bimanual Load-Lifting Task. Frontiers in Human Neuroscience, 5(163). 10.3389/fnhum.2011.00163

6. Barlaam, F., Fortin, C., Vaugoyeau, M., Schmitz, C., & Assaiante, C. (2012). Development of action representation during adolescence as assessed from anticipatory control in a bimanual load-lifting task. Neuroscience, 221, 56–68. 10.1016/j.neuroscience.2012.06.062

7. Barlaam, F., Fortin, C., Vaugoyeau, M., Schmitz, C., & Assaiante, C. (2018). Mu-oscillation changes related to the development of anticipatory postural control in children and adolescents. Journal of Neurophysiology, 120(1), 129–138. 10.1152/jn.00637.2017

8. Barnea-Goraly, N., Menon, V., Eckert, M., Tamm, L., Bammer, R., Karchemskiy, A., Dant, C. C., & Reiss, A. L. (2005). White Matter Development During Childhood and Adolescence: A Cross-sectional Diffusion Tensor Imaging Study. Cerebral Cortex, 15(12), 1848–1854. 10.1093/cercor/bhi062

9. Bonini, L., Maranesi, M., Livi, A., Fogassi, L., & Rizzolatti, G. (2014). Space-Dependent Representation of Objects and Other’s Action in Monkey Ventral Premotor Grasping Neurons. The Journal of Neuroscience, 34(11), 4108–4119. 10.1523/JNEUROSCI.4187-13.2014

10. Bonnefond, M., & Jensen, O. (2015). Gamma Activity Coupled to Alpha Phase as a Mechanism for Top-Down Controlled Gating. PLOS ONE, 10(6), e0128667. 10.1371/journal.pone.0128667

11. Bönstrup, M., Hagemann, J., Gerloff, C., Sauseng, P., & Hummel, F. C. (2015). Alpha oscillatory correlates of motor inhibition in the aged brain. Frontiers in Aging Neuroscience, 7. 10.3389/fnagi.2015.00193

12. Bräcklein, M., Barsakcioglu, D. Y., Del Vecchio, A., Ibáñez, J., & Farina, D. (2022). Reading and Modulating Cortical β Bursts from Motor Unit Spiking Activity. The Journal of Neuroscience, 42(17), 3611–3621. 10.1523/JNEUROSCI.1885-21.2022

13. Brazhnik, E., Novikov, N., McCoy, A. J., Ilieva, N. M., Ghraib, M. W., & Walters, J. R. (2021). Early decreases in cortical mid-gamma peaks coincide with the onset of motor deficits and precede exaggerated beta build-up in rat models for Parkinson’s disease. Neurobiology of Disease, 155, 105393. 10.1016/j.nbd.2021.105393

14. Bressler, S. L., & Seth, A. K. (2011). Wiener–Granger Causality: A well established methodology. NeuroImage, 58(2), 323–329. 10.1016/j.neuroimage.2010.02.059

15. Brookes, M. J., Vrba, J., Robinson, S. E., Stevenson, C. M., Peters, A. M., Barnes, G. R., Hillebrand, A., & Morris, P. G. (2008). Optimising experimental design for MEG beamformer imaging. NeuroImage, 39(4), 1788–1802. 10.1016/j.neuroimage.2007.09.050

16. Buschman, T. J., Denovellis, E. L., Diogo, C., Bullock, D., & Miller, E. K. (2012). Synchronous Oscillatory Neural Ensembles for Rules in the Prefrontal Cortex. Neuron, 76(4), 838–846. 10.1016/j.neuron.2012.09.029

17. Chen, S., Entakli, J., Bonnard, M., Berton, E., & De Graaf, J. B. (2013). Functional Corticospinal Projections from Human Supplementary Motor Area Revealed by Corticomuscular Coherence during Precise Grip Force Control. PLoS ONE, 8(3), e60291. 10.1371/journal.pone.0060291

18. Davare, M., Montague, K., Olivier, E., Rothwell, J. C., & Lemon, R. N. (2009). Ventral premotor to primary motor cortical interactions during object-driven grasp in humans. Cortex, 45(9), 1050–1057. 10.1016/j.cortex.2009.02.011

19. Di Rienzo, F., Barlaam, F., Daligault, S., Delpuech, C., Roy, A. C., Bertrand, O., Jerbi, K., & Schmitz, C. (2019). Tracking the acquisition of anticipatory postural adjustments during a bimanual load-lifting task: A MEG study. Human Brain Mapping, 40(10), 2955–2966. 10.1002/hbm.24571

20. Donoghue, T., Haller, M., Peterson, E. J., Varma, P., Sebastian, P., Gao, R., Noto, T., Lara, A. H., Wallis, J. D., Knight, R. T., Shestyuk, A., & Voytek, B. (2020). Parameterizing neural power spectra into periodic and aperiodic components. Nature Neuroscience, 23(12), 1655–1665. 10.1038/s41593-020-00744-x

21. Dufossé, M., Hugon, M., & Massion, J. (1985). Postural forearm changes induced by predictable in time or voluntary triggered unloading in man. Experimental Brain Research, 60(2). 10.1007/BF00235928

22. Dum, R., & Strick, P. (1991). The origin of corticospinal projections from the premotor areas in the frontal lobe. The Journal of Neuroscience, 11(3), 667–689. 10.1523/JNEUROSCI.11-03-00667.1991

23. Echeverria-Altuna, I., Quinn, A. J., Zokaei, N., Woolrich, M. W., Nobre, A. C., & Van Ede, F. (2022). Transient beta activity and cortico-muscular connectivity during sustained motor behaviour. Progress in Neurobiology, 214, 102281. 10.1016/j.pneurobio.2022.102281

24. Edgar, J. C., Dipiero, M., McBride, E., Green, H. L., Berman, J., Ku, M., Liu, S., Blaskey, L., Kuschner, E., Airey, M., Ross, J. L., Bloy, L., Kim, M., Koppers, S., Gaetz, W., Schultz, R. T., & Roberts, T. P. L. (2019). Abnormal maturation of the resting-state peak alpha frequency in children with autism spectrum disorder. Human Brain Mapping, 40(11), 3288–3298. 10.1002/hbm.24598

25. Entakli, J., Bonnard, M., Chen, S., Berton, E., & De Graaf, J. B. (2014). TMS reveals a direct influence of spinal projections from human SMAp on precise force production. European Journal of Neuroscience, 39(1), 132–140. 10.1111/ejn.12392

26. Fischl, B., Salat, D. H., Busa, E., Albert, M., Dieterich, M., Haselgrove, C., Van Der Kouwe, A., Killiany, R., Kennedy, D., Klaveness, S., Montillo, A., Makris, N., Rosen, B., & Dale, A. M. (2002). Whole Brain Segmentation. Neuron, 33(3), 341–355. 10.1016/S0896-6273(02)00569-X

27. Gallese, V., Fadiga, L., Fogassi, L., & Rizzolatti, G. (1996). Action recognition in the premotor cortex. Brain, 119(2), 593–609. 10.1093/brain/119.2.593

28. Glasser, M. F., Coalson, T. S., Robinson, E. C., Hacker, C. D., Harwell, J., Yacoub, E., Ugurbil, K., Andersson, J., Beckmann, C. F., Jenkinson, M., Smith, S. M., & Van Essen, D. C. (2016). A multi-modal parcellation of human cerebral cortex. Nature, 536(7615), 171–178. 10.1038/nature18933

29. Gogtay, N., Giedd, J. N., Lusk, L., Hayashi, K. M., Greenstein, D., Vaituzis, A. C., Nugent, T. F., Herman, D. H., Clasen, L. S., Toga, A. W., Rapoport, J. L., & Thompson, P. M. (2004). Dynamic mapping of human cortical development during childhood through early adulthood. Proceedings of the National Academy of Sciences, 101(21), 8174–8179. 10.1073/pnas.0402680101

30. Graef, C., Pascual Valdunciel, A., Farina, D., Vaidyanathan, R., Tai, Y. F., & Haar, S. (2026). Propagation of beta bursts from the motor cortex to the motor units of multiple upper-limb muscles. Journal of Neurophysiology, 135(1), 97–109. 10.1152/jn.00336.2025

31. Gramfort, A., Luessi, M., Larson, E., Engemann, D. A., Strohmeier, D., Brodbeck, C., Goj, R., Jas, M., Brooks, T., Parkkonen, L., & Hämäläinen, M. (2013). MEG and EEG data analysis with MNE-Python. Frontiers in Neuroscience, 7, 267. 10.3389/fnins.2013.00267

32. Haegens, S., Nácher, V., Luna, R., Romo, R., & Jensen, O. (2011). α-Oscillations in the monkey sensorimotor network influence discrimination performance by rhythmical inhibition of neuronal spiking. Proceedings of the National Academy of Sciences, 108(48), 19377–19382. 10.1073/pnas.1117190108

33. Hagne, I. (1972). Development of the Sleep EEG in Normal Infants During the First Year of Life. Acta Paediatrica, 61(S232), 25–53. 10.1111/j.1651-2227.1972.tb08271.x

34. Henderson, S. E., Sugden, D., & Barnett, A. L. (2019). *Movement Assessment Battery for Children-2* [Dataset]. 10.1037/t55281-000

35. Hugon, M., Massion, J., & Wiesendanger, M. (1982). Anticipatory postural changes induced by active unloading and comparison with passive unloading in man. Pflügers Archiv European Journal of Physiology, 393(4), 292–296. 10.1007/BF00581412

36. Hummel, F., Andres, F., Altenmüller, E., Dichgans, J., & Gerloff, C. (2002). Inhibitory control of acquired motor programmes in the human brain. Brain, 125(2), 404–420. 10.1093/brain/awf030

37. Ioffe, M. E., Massion, J., Schmitz, C., Viallet, F., & Gancheva, R. (2003). Specific Functions of the Motor Cortex in Reorganizing Coordinations during Motor Training in Animals and Humans. Neuroscience and Behavioral Physiology, 33(2), 143–150. 10.1023/A:1021717829999

38. Ivry, R. B., & Keele, S. W. (1989). Timing Functions of The Cerebellum. Journal of Cognitive Neuroscience, 1(2), 136–152. 10.1162/jocn.1989.1.2.136

39. Jaiswal, A., Nenonen, J., Stenroos, M., Gramfort, A., Dalal, S. S., Westner, B. U., Litvak, V., Mosher, J. C., Schoffelen, J.-M., Witton, C., Oostenveld, R., & Parkkonen, L. (2020). Comparison of beamformer implementations for MEG source localization. NeuroImage, 216, 116797. 10.1016/j.neuroimage.2020.116797

40. Jensen, O., & Mazaheri, A. (2010). Shaping Functional Architecture by Oscillatory Alpha Activity: Gating by Inhibition. Frontiers in Human Neuroscience, 4. 10.3389/fnhum.2010.00186

41. Kang, S., Jun, S., Baek, S. J., Park, H., Yamamoto, Y., & Tanaka-Yamamoto, K. (2021). Recent Advances in the Understanding of Specific Efferent Pathways Emerging From the Cerebellum. Frontiers in Neuroanatomy, 15, 759948. 10.3389/fnana.2021.759948

42. Kazennikov, O., Solopova, I., Talis, V., Grishin, A., & Ioffe, M. (2005). TMS-responses during anticipatory postural adjustment in bimanual unloading in humans. Neuroscience Letters, 383(3), 246–250. 10.1016/j.neulet.2005.04.005

43. Kazennikov, O., Solopova, I., Talis, V., & Ioffe, M. (2008). Anticipatory postural adjustment: The role of motor cortex in the natural and learned bimanual unloading. Experimental Brain Research, 186(2), 215–223. 10.1007/s00221-007-1224-5

44. Kelley, C., Slater, C., Sorrentino, M., Noone, D., Hung, J., Sajda, P., & Wang, Q. (2025). Alpha modulation of spiking activity across multiple brain regions in mice performing a tactile selective detection task. Neuroscience. 10.1101/2025.03.07.642074

45. Khan, A. U., Irwin, Z., Mahavadi, A., Roller, A., Goodman, A. M., Guthrie, B. L., Visscher, K., Knight, R. T., Walker, H. C., & Bentley, J. N. (2024). Low-Frequency Oscillations in Mid-rostral Dorsolateral Prefrontal Cortex Support Response Inhibition. The Journal of Neuroscience, 44(40), e0122242024. 10.1523/JNEUROSCI.0122-24.2024

46. Khanna, P., Farrokhi, B., Choi, H., Griffin, S., Heimbuch, I., Novik, L., Thiesen, K., Morrison, J., Morecraft, R. J., & Ganguly, K. (2025). Separable global and local beta burst dynamics in motor cortex of primates. 10.7554/eLife.107473.1

47. Klimesch, W., Sauseng, P., & Hanslmayr, S. (2007). EEG alpha oscillations: The inhibition–timing hypothesis. Brain Research Reviews, 53(1), 63–88. 10.1016/j.brainresrev.2006.06.003

48. Lacquaniti, F., & Maioli, C. (1989). The role of preparation in tuning anticipatory and reflex responses during catching. The Journal of Neuroscience, 9(1), 134–148. 10.1523/JNEUROSCI.09-01-00134.1989

49. Lotze, M., Erb, M., Flor, H., Huelsmann, E., Godde, B., & Grodd, W. (2000). fMRI Evaluation of Somatotopic Representation in Human Primary Motor Cortex. NeuroImage, 11(5), 473–481. 10.1006/nimg.2000.0556

50. Lundqvist, M., Miller, E. K., Nordmark, J., Liljefors, J., & Herman, P. (2024). Beta: Bursts of cognition. Trends in Cognitive Sciences, 28(7), 662–676. 10.1016/j.tics.2024.03.010

51. Lundqvist, M., Rose, J., Herman, P., Brincat, S. L., Buschman, T. J., & Miller, E. K. (2016). Gamma and Beta Bursts Underlie Working Memory. Neuron, 90(1), 152–164. 10.1016/j.neuron.2016.02.028

52. Manyukhina, V., Abdoun, O., Di Rienzo, F., Barlaam, F., Daligault, S., Delpuech, C., Szul, M., Bonaiuto, J., Bonnefond, M., & Schmitz, C. (2026). Beta bursts in SMA mediate anticipatory muscle inhibition. Cerebral Cortex, 36(6), bhag054. 10.1093/cercor/bhag054

53. Martineau, J., Schmitz, C., Assaiante, C., Blanc, R., & Barthélémy, C. (2004). Impairment of a cortical event-related desynchronisation during a bimanual load-lifting task in children with autistic disorder. Neuroscience Letters, 367(3), 298–303. 10.1016/j.neulet.2004.06.018

54. Massion, J. (1992). Movement, posture and equilibrium: Interaction and coordination. Progress in Neurobiology, 38(1), 35–56. 10.1016/0301-0082(92)90034-C

55. Massion, J., Ioffe, M., Schmitz, C., Viallet, F., & Gantcheva, R. (1999). Acquisition of anticipatory postural adjustments in a bimanual load-lifting task: Normal and pathological aspects. Experimental Brain Research, 128(1–2), 229–235. 10.1007/s002210050842

56. McNab, F., Leroux, G., Strand, F., Thorell, L., Bergman, S., & Klingberg, T. (2008). Common and unique components of inhibition and working memory: An fMRI, within-subjects investigation. Neuropsychologia, 46(11), 2668–2682. 10.1016/j.neuropsychologia.2008.04.023

58. Miskovic, V., Ma, X., Chou, C.-A., Fan, M., Owens, M., Sayama, H., & Gibb, B. E. (2015). Developmental changes in spontaneous electrocortical activity and network organization from early to late childhood. NeuroImage, 118, 237–247. 10.1016/j.neuroimage.2015.06.013

59. Moca, V. V., Bârzan, H., Nagy-Dăbâcan, A., & Mureșan, R. C. (2021). Time-frequency super-resolution with superlets. Nature Communications, 12(1), 337. 10.1038/s41467-020-20539-9

60. Murthy, V. N., & Fetz, E. E. (1996). Oscillatory activity in sensorimotor cortex of awake monkeys: Synchronization of local field potentials and relation to behavior. Journal of Neurophysiology, 76(6), 3949–3967. 10.1152/jn.1996.76.6.3949

61. Muthukumaraswamy, S. D. (2013). High-frequency brain activity and muscle artifacts in MEG/EEG: A review and recommendations. Frontiers in Human Neuroscience, 7. 10.3389/fnhum.2013.00138

62. Ng, T. H. B., Sowman, P. F., Brock, J., & Johnson, B. W. (2011). Premovement brain activity in a bimanual load-lifting task. Experimental Brain Research, 208(2), 189–201. 10.1007/s00221-010-2470-5

63. Ng, T. H. B., Sowman, P. F., Brock, J., & Johnson, B. W. (2013a). Neuromagnetic brain activity associated with anticipatory postural adjustments for bimanual load lifting. NeuroImage, 66, 343–352. 10.1016/j.neuroimage.2012.10.042

64. Ng, T. H. B., Sowman, P. F., Brock, J., & Johnson, B. W. (2013b). Neuromagnetic imaging reveals timing of volitional and anticipatory motor control in bimanual load lifting. Behavioural Brain Research, 247, 182–192. 10.1016/j.bbr.2013.03.020

65. Pagge, C., Caballero-Insaurriaga, J., Oliviero, A., Foffani, G., & Ammann, C. (2024). Transcranial static magnetic field stimulation of the supplementary motor area decreases corticospinal excitability in the motor cortex: A pilot study. Scientific Reports, 14(1), 6597. 10.1038/s41598-024-57030-0

66. Paulignan, Y., Dufossé, M., Hugon, M., & Massion, J. (1989). Acquisition of co-ordination between posture and movement in a bimanual task. Experimental Brain Research, 77(2), 337–348. 10.1007/BF00274991

67. Plow, E. B., Arora, P., Pline, M. A., Binenstock, M. T., & Carey, J. R. (2010). Within-limb somatotopy in primary motor cortex – revealed using fMRI. Cortex, 46(3), 310–321. 10.1016/j.cortex.2009.02.024

68. Purves, D., Augustine, G. J., & Fitzpatrick, D. (Eds.). (2018). Neuroscience (Sixth edition). Oxford University Press, Sinauer Associates is an imprint of Oxford Universitiy Press.

69. Ray, S., Crone, N. E., Niebur, E., Franaszczuk, P. J., & Hsiao, S. S. (2008). Neural correlates of high-gamma oscillations (60–200 Hz) in macaque local field potentials and their potential implications in electrocorticography. Journal of Neuroscience, 28(45), 11526–11536.

70. Ray, S., & Maunsell, J. H. R. (2011). Different Origins of Gamma Rhythm and High-Gamma Activity in Macaque Visual Cortex. PLoS Biology, 9(4), e1000610. 10.1371/journal.pbio.1000610

71. Rayson, H., Szul, M. J., El-Khoueiry, P., Debnath, R., Gautier-Martins, M., Ferrari, P. F., Fox, N., & Bonaiuto, J. J. (2023). Bursting with Potential: How Sensorimotor Beta Bursts Develop from Infancy to Adulthood. The Journal of Neuroscience, 43(49), 8487–8503. 10.1523/JNEUROSCI.0886-23.2023

72. Reddy, V., Markova, G., & Wallot, S. (2013). Anticipatory Adjustments to Being Picked Up in Infancy. PLoS ONE, 8(6), e65289. 10.1371/journal.pone.0065289

73. Riehle, A., Brochier, T., Nawrot, M., & Grün, S. (2018). Behavioral Context Determines Network State and Variability Dynamics in Monkey Motor Cortex. Frontiers in Neural Circuits, 12, 52. 10.3389/fncir.2018.00052

74. Rubia, K., Russell, T., Overmeyer, S., Brammer, M. J., Bullmore, E. T., Sharma, T., Simmons, A., Williams, S. C. R., Giampietro, V., Andrew, C. M., & Taylor, E. (2001). Mapping Motor Inhibition: Conjunctive Brain Activations across Different Versions of Go/No-Go and Stop Tasks. NeuroImage, 13(2), 250–261. 10.1006/nimg.2000.0685

75. Salman, M. S. (2002). Topical Review: The Cerebellum: It’s About Time! But Timing Is Not Everything-New Insights Into the Role of the Cerebellum in Timing Motor and Cognitive Tasks. Journal of Child Neurology, 17(1), 1–9. 10.1177/088307380201700101

76. Samaha, J., Bauer, P., Cimaroli, S., & Postle, B. R. (2015). Top-down control of the phase of alpha-band oscillations as a mechanism for temporal prediction. Proceedings of the National Academy of Sciences, 112(27), 8439–8444. 10.1073/pnas.1503686112

77. Sauseng, P., Gerloff, C., & Hummel, F. C. (2013). Two brakes are better than one: The neural bases of inhibitory control of motor memory traces. NeuroImage, 65, 52–58. 10.1016/j.neuroimage.2012.09.048

78. Schmitz, C., Jenmalm, P., Ehrsson, H. H., & Forssberg, H. (2005). Brain Activity During Predictable and Unpredictable Weight Changes When Lifting Objects. Journal of Neurophysiology, 93(3), 1498–1509. 10.1152/jn.00230.2004

79. Schmitz, C., Martin, N., & Assaiante, C. (1999). Development of anticipatory postural adjustments in a bimanual load-lifting task in children. Experimental Brain Research, 126(2), 200–204. 10.1007/s002210050729

80. Schmitz, C., Martin, N., & Assaiante, C. (2002). Building anticipatory postural adjustment during childhood: A kinematic and electromyographic analysis of unloading in children from 4 to 8 years of age. Experimental Brain Research, 142(3), 354–364. 10.1007/s00221-001-0910-REFy

81. Sherman, M. A., Lee, S., Law, R., Haegens, S., Thorn, C. A., Hämäläinen, M. S., Moore, C. I., & Jones, S. R. (2016). Neural mechanisms of transient neocortical beta rhythms: Converging evidence from humans, computational modeling, monkeys, and mice. Proceedings of the National Academy of Sciences, 113(33), E4885–E4894. 10.1073/pnas.1604135113

82. Smith, J. R. (1941). The Frequency Growth of the Human Alpha Rhythms During Normal Infancy and Childhood. The Journal of Psychology, 11(1), 177–198. 10.1080/00223980.1941.9917028

83. Sohal, V. S., & Rubenstein, J. L. R. (2019). Excitation-inhibition balance as a framework for investigating mechanisms in neuropsychiatric disorders. Molecular Psychiatry, 24(9), 1248–1257. 10.1038/s41380-019-0426-0

84. Solís-Vivanco, R., Jensen, O., & Bonnefond, M. (2018). Top–Down Control of Alpha Phase Adjustment in Anticipation of Temporally Predictable Visual Stimuli. Journal of Cognitive Neuroscience, 30(8), 1157–1169. 10.1162/jocn_a_01280

85. Soppelsa, R., & Albaret, J. M. (2005). La batterie d’évaluation du mouvement chez l’enfant (M-ABC): Étalonnage sur une population d’enfants de 4 à 12 ans.

86. Sóskuthy, M. (2017). *Generalised additive mixed models for dynamic analysis in linguistics: A practical introduction* (No. arXiv:1703.05339). arXiv. 10.48550/arXiv.1703.05339

87. Sowell, E. R., Thompson, P. M., Leonard, C. M., Welcome, S. E., Kan, E., & Toga, A. W. (2004). Longitudinal Mapping of Cortical Thickness and Brain Growth in Normal Children. The Journal of Neuroscience, 24(38), 8223–8231. 10.1523/JNEUROSCI.1798-04.2004

88. Spaak, E., Bonnefond, M., Maier, A., Leopold, D. A., & Jensen, O. (2012). Layer-Specific Entrainment of Gamma-Band Neural Activity by the Alpha Rhythm in Monkey Visual Cortex. Current Biology, 22(24), 2313–2318. 10.1016/j.cub.2012.10.020

89. Spieser, L., Aubert, S., & Bonnard, M. (2013). Involvement of SMAp in the intention-related long latency stretch reflex modulation: A TMS study. Neuroscience, 246, 329–341. 10.1016/j.neuroscience.2013.05.005

90. Stroganova, T. A., Orekhova, E. V., & Posikera, I. N. (1999). EEG alpha rhythm in infants. Clinical Neurophysiology, 110(6), 997–1012. 10.1016/S1388-2457(98)00009-1

91. Suda, A., Osada, T., Ogawa, A., Tanaka, M., Kamagata, K., Aoki, S., Hattori, N., & Konishi, S. (2020). Functional Organization for Response Inhibition in the Right Inferior Frontal Cortex of Individual Human Brains. Cerebral Cortex, 30(12), 6325–6335. 10.1093/cercor/bhaa188

92. Szul, M. J., Papadopoulos, S., Alavizadeh, S., Daligaut, S., Schwartz, D., Mattout, J., & Bonaiuto, J. J. (2023). Diverse beta burst waveform motifs characterize movement-related cortical dynamics. Progress in Neurobiology, 228, 102490. 10.1016/j.pneurobio.2023.102490

93. Teitti, S., Määttä, S., Säisänen, L., Könönen, M., Vanninen, R., Hannula, H., Mervaala, E., & Karhu, J. (2008). Non-primary motor areas in the human frontal lobe are connected directly to hand muscles. NeuroImage, 40(3), 1243–1250. 10.1016/j.neuroimage.2007.12.065

94. Tiemeier, H., Lenroot, R. K., Greenstein, D. K., Tran, L., Pierson, R., & Giedd, J. N. (2010). Cerebellum development during childhood and adolescence: A longitudinal morphometric MRI study. NeuroImage, 49(1), 63–70. 10.1016/j.neuroimage.2009.08.016

95. Van Pelt, S., Heil, L., Kwisthout, J., Ondobaka, S., Van Rooij, I., & Bekkering, H. (2016). Beta- and gamma-band activity reflect predictive coding in the processing of causal events. Social Cognitive and Affective Neuroscience, 11(6), 973–980. 10.1093/scan/nsw017

96. Viallet, F., Massion, J., Massarino, R., & Khalil, R. (1987). Performance of a bimanual load-lifting task by parkinsonian patients. *Journal of Neurology*, Neurosurgery & Psychiatry, 50(10), 1274–1283. 10.1136/jnnp.50.10.1274

97. Viallet, F., Massion, J., Massarino, R., & Khalil, R. (1992). Coordination between posture and movement in a bimanual load lifting task: Putative role of a medial frontal region including the supplementary motor area. Experimental Brain Research, 88(3). 10.1007/BF00228197

98. Whitham, E. M., Lewis, T., Pope, K. J., Fitzgibbon, S. P., Clark, C. R., Loveless, S., DeLosAngeles, D., Wallace, A. K., Broberg, M., & Willoughby, J. O. (2008). Thinking activates EMG in scalp electrical recordings. Clinical Neurophysiology, 119(5), 1166–1175. 10.1016/j.clinph.2008.01.024

99. Wieling, M. (2018). Analyzing dynamic phonetic data using generalized additive mixed modeling: A tutorial focusing on articulatory differences between L1 and L2 speakers of English. Journal of Phonetics, 70, 86–116. 10.1016/j.wocn.2018.03.002

100. Wilkinson, C. L., Yankowitz, L. D., Chao, J. Y., Gutiérrez, R., Rhoades, J. L., Shinnar, S., Purdon, P. L., & Nelson, C. A. (2024). Developmental trajectories of EEG aperiodic and periodic components in children 2–44 months of age. Nature Communications, 15(1), 5788. 10.1038/s41467-024-50204-4

101. Yazdan-Shahmorad, A., Kipke, D. R., & Lehmkuhle, M. J. (2013). High gamma power in ECoG reflects cortical electrical stimulation effects on unit activity in layers V/VI. Journal of Neural Engineering, 10(6), 066002. 10.1088/1741-2560/10/6/066002

102. Zheng, D., Oka, T., Bokura, H., & Yamaguchi, S. (2008). The Key Locus of Common Response Inhibition Network for No-go and Stop Signals. Journal of Cognitive Neuroscience, 20(8), 1434–1442. 10.1162/jocn.2008.20100

