## Supplementary Materials for "SMA beta bursts reveal distinct mechanisms of anticipatory motor control in children"

*corresponding author

### Single-trial illustration of the automated *Biceps brachii* EMG inhibition detection algorithm

The three panels below provide single-trial examples illustrating the automated *Biceps brachii* EMG inhibition detection algorithm. Figure S1A shows examples in which inhibition events were clearly identified both manually and automatically. Figure S1B presents cases in which the automated detection identified inhibitory events that were not detected manually. Figure S1C shows two examples in which the automated approach failed to detect long inhibition events identified manually, alongside one example of a long inhibition that was successfully detected, indicating that the automated method was also capable of detecting such events.

Each trial illustrates the five steps of the *Biceps brachii* inhibition peak detection algorithm:

1) Rectified EMG signal. Blue rectangles (when present) indicate manually annotated inhibitions, shown for comparison with the automatic detection. Violet lines mark the unloading time, around which inhibition was identified both manually and automatically. Grey and pink lines represent additional annotations not relevant for the present plots.

2) Single-trial time-frequency representation (TFR) of the EMG signal (0 – unloading).

3) Thresholded TFR, obtained using a threshold of mean - 1.5 × SD.

4) TFRs after thresholding and exclusion of non-persistent EMG decreases (assessed separately for each frequency bin), ensuring that only reliable power reductions were used for inhibition peak estimation.

5) Time course used to detect the inhibition peak (red line) and its boundaries (red dashed lines), obtained by averaging across frequencies from the previous step and multiplying by -1 so that inhibition appears as a positive peak.

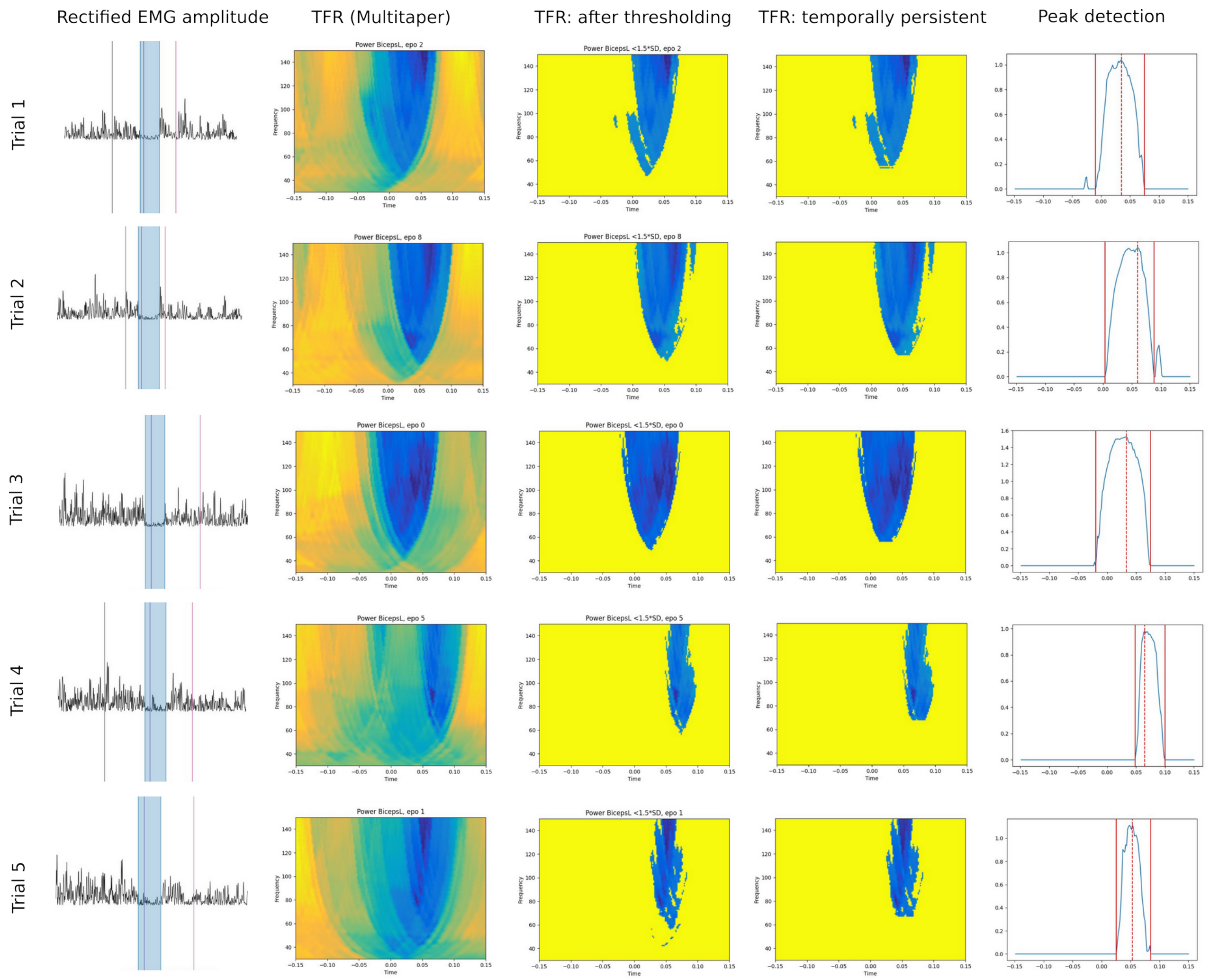


**Figure S1A.** *Biceps brachii* inhibition events clearly identified both manually and automatically.

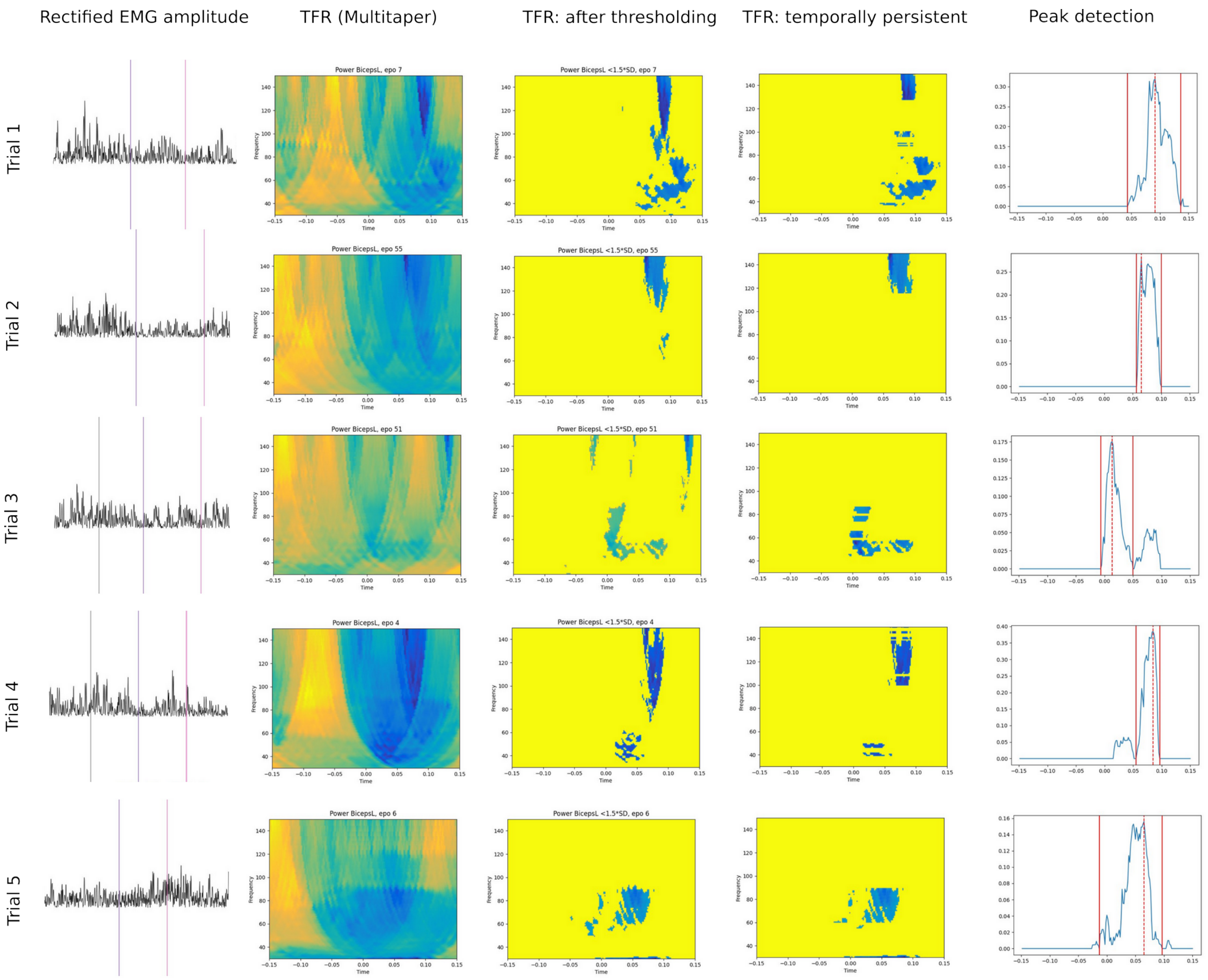


**Figure S1B.** *Biceps brachii* inhibition events identified by the automatic detection method but not detected manually.

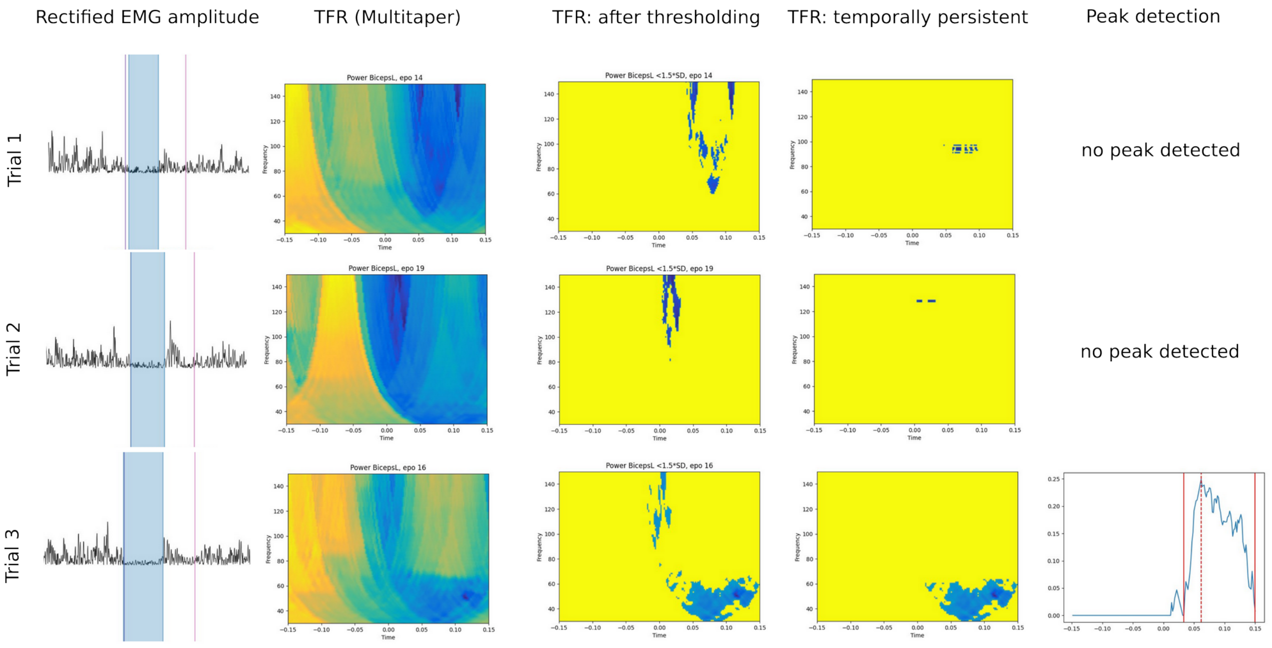

**Figure S1C.** Trials 1-2: Examples of *Biceps brachii* inhibition in which the automatic approach failed to detect long inhibition events that were identified manually. Trial 3 illustrates an example of a long inhibition that was also successfully detected by the automatic approach.

### GAM: Prediction of forearm stabilization from *Biceps brachii* inhibition parameters

We fitted a GAM to determine which *Biceps brachii* inhibition parameters – Inhibition Onset Time, Inhibition Peak Time, Peak EMG Inhibition (inhibition strength), and Inhibition Duration, and their interactions – best predicted forearm stabilization, indexed by Peak Elbow Rotation.

Schematic model fit representation:

*Peak Elbow Rotation ~ Peak Inhibition + Inhibition Peak Time + Inhibition Duration + Inhibition Onset + paired interactions*

Exact model specification:

*PeakElbRot ~ s(Inh_ampl) + s(Inh_peak_time) + s(Inh_onset_time) + s(Inh_duration) +*
*ti(Inh_duration, Inh_onset_time) + ti(Inh_ampl, Inh_onset_time) +*
*ti(Inh_ampl, Inh_duration) + ti(Inh_duration, Inh_peak_time) +*
*ti(Inh_ampl, Inh_peak_time) + ti(Inh_onset_time, Inh_peak_time) +*
*s(Subject, Inh_ampl, bs='re') + s(Subject, Inh_duration, bs='re') +*
*s(Subject, Inh_onset_time, bs='re') + s(Subject, Inh_peak_time, bs='re') +*
*s(Subject, bs='re')*

Full model results are provided in Table S1. The GAM revealed a significant main effect of Inhibition Peak Time ($F = 3.52$, $p = 0.01$) and a significant interaction between Peak EMG Inhibition and Inhibition Peak Time ($F = 3.17$, $p = 0.02$), illustrated in Figure S2.

**Table S1.** GAM smooth-term results predicting Peak Elbow Rotation from *Biceps brachii* inhibition measures and their interactions.

| Term | edf | Ref.df | F | p-value |
|---|---|---|---|---|
| s(Inh_ampl) | 1.00 | 1.00 | 1.14 | 0.286 |
| s(Inh_peak_time) | 2.84 | 3.61 | 3.52 | 0.010* |
| s(Inh_onset_time) | 2.66 | 3.41 | 1.69 | 0.156 |
| s(Inh_duration) | 1.00 | 1.00 | 0.98 | 0.322 |
| ti(Inh_duration, Inh_onset_time) | 1.00 | 1.00 | 2.59 | 0.108 |
| ti(Inh_ampl, Inh_onset_time) | 5.26 | 7.03 | 1.36 | 0.233 |
| ti(Inh_ampl, Inh_duration) | 2.97 | 4.19 | 0.91 | 0.449 |
| ti(Inh_duration, Inh_peak_time) | 1.00 | 1.00 | 2.24 | 0.135 |
| ti(Inh_ampl, Inh_peak_time) | 2.69 | 3.18 | 3.17 | 0.020* |
| ti(Inh_onset_time, Inh_peak_time) | 1.00 | 1.00 | 0.36 | 0.547 |
| s(Subject, Inh_ampl) | <0.001 | 23.00 | 0.00 | 0.792 |
| s(Subject, Inh_duration) | <0.001 | 23.00 | 0.00 | 0.924 |
| s(Subject, Inh_onset_time) | 0.002 | 23.00 | 0.00 | 0.442 |
| s(Subject, Inh_peak_time) | 9.59 | 23.00 | 7.10 | 0.023* |
| s(Subject) | 19.95 | 23.00 | 29.13 | < 0.001*** |

PeakElbRot: Peak Elbow Rotation; Inh_ampl: Peak EMG Inhibition; Inh_peak_time: EMG Inhibition Peak time; Inh_onset_time: EMG Inhibition Onset time; Inh_duration: EMG Inhibition duration; ti(Inh_duration, Inh_onset_time): tensor product interaction between smooth terms; s(Subject, Inh_ampl, bs = "re"): subject-based random slope; s(Subject, bs = "re"): subject-based random intercept.

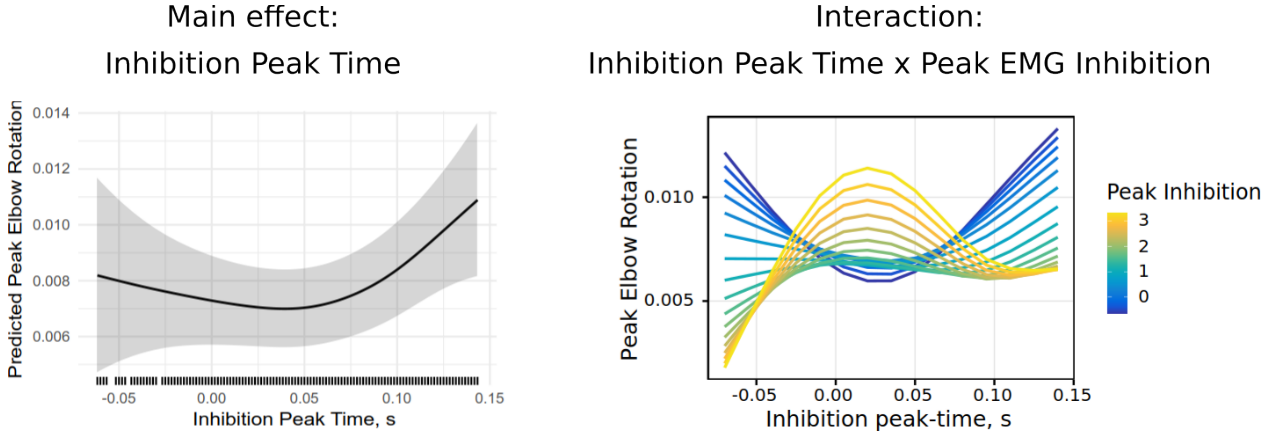


**Figure S2.** Significant main effect of peak *Biceps brachii* inhibition time and its interaction with inhibition strength on maximal elbow rotation (see Table 1). More negative Peak Inhibition indicates stronger inhibition.

### SMA and M1 regions for correlation analysis between high-gamma power and *Biceps brachii* inhibition strength

To test whether stronger *Biceps brachii* inhibition is associated with reduced excitability in SMA or M1, we performed cluster-based analyses within predefined SMA and M1 labels (Fig. 2A).

The SMA was defined based on peak effects observed in adults in our previous study (Manyukhina et al., 2026), and the M1 label was based on the hand/arm representation of M1 reported in the literature (Lotze et al., 2000; Plow et al., 2010). The SMA and M1 regions are illustrated in Figure S3.

For comparison with adults (Manyukhina et al., 2026), the correlation effect observed in children (Fig. 2A) is shown alongside the peak adult effect (blue marker) in Figure S3.

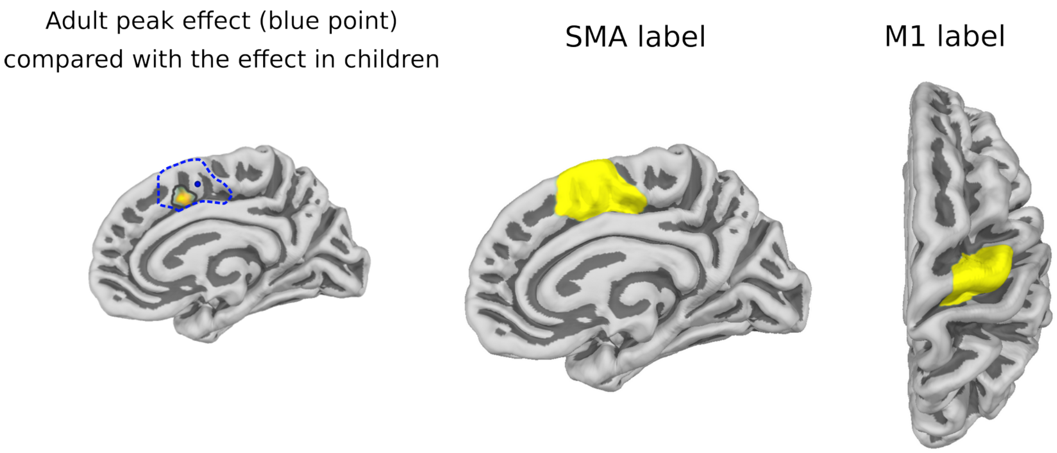


**Figure S3.** SMA and M1 labels used to test whether stronger *Biceps brachii* inhibition is associated with reduced high-gamma (90-130 Hz) power in SMA or M1 (see Fig. 2A, main text). The left panel reproduces the significant SMA correlation effect observed in the present sample and compares its localization with the peak effect reported in adults (blue point; Manyukhina et al., 2026). The dashed blue line indicates the boundaries of the SMA label used in the analysis in children.

**Distribution of elbow rotation-derived and *Biceps brachii* inhibition-derived behavioral measures across subjects**

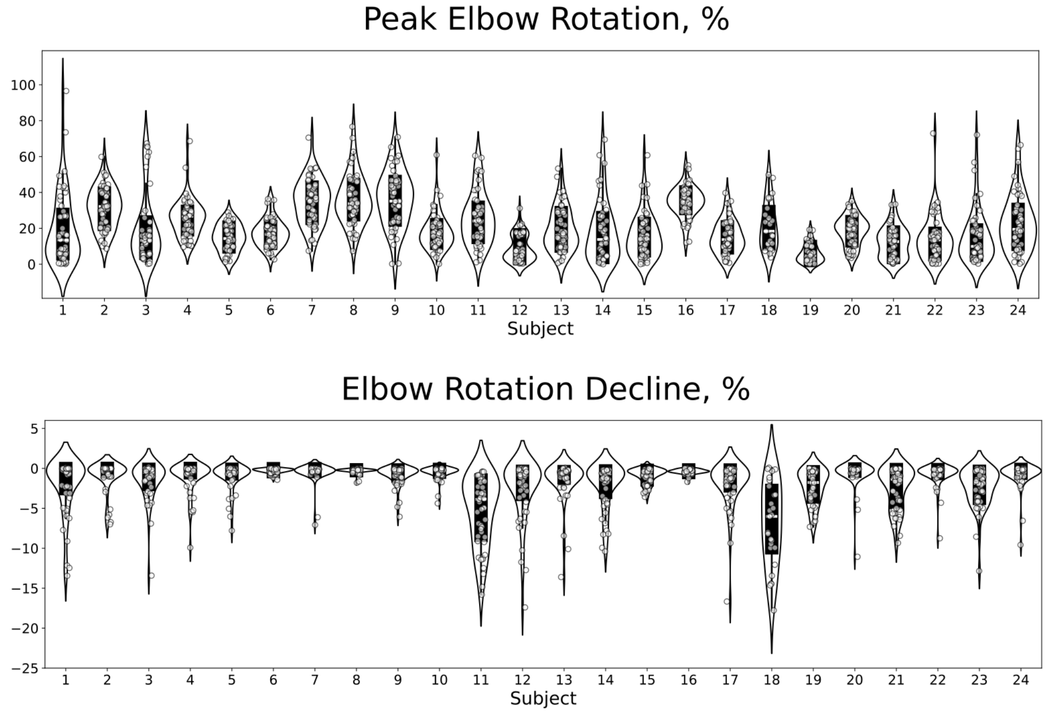


**Figure S4.** Violin plots of elbow rotation-derived parameters (Elbow Rotation Decline, Peak Elbow Rotation) for each subject, showing the distribution of values and individual data points.

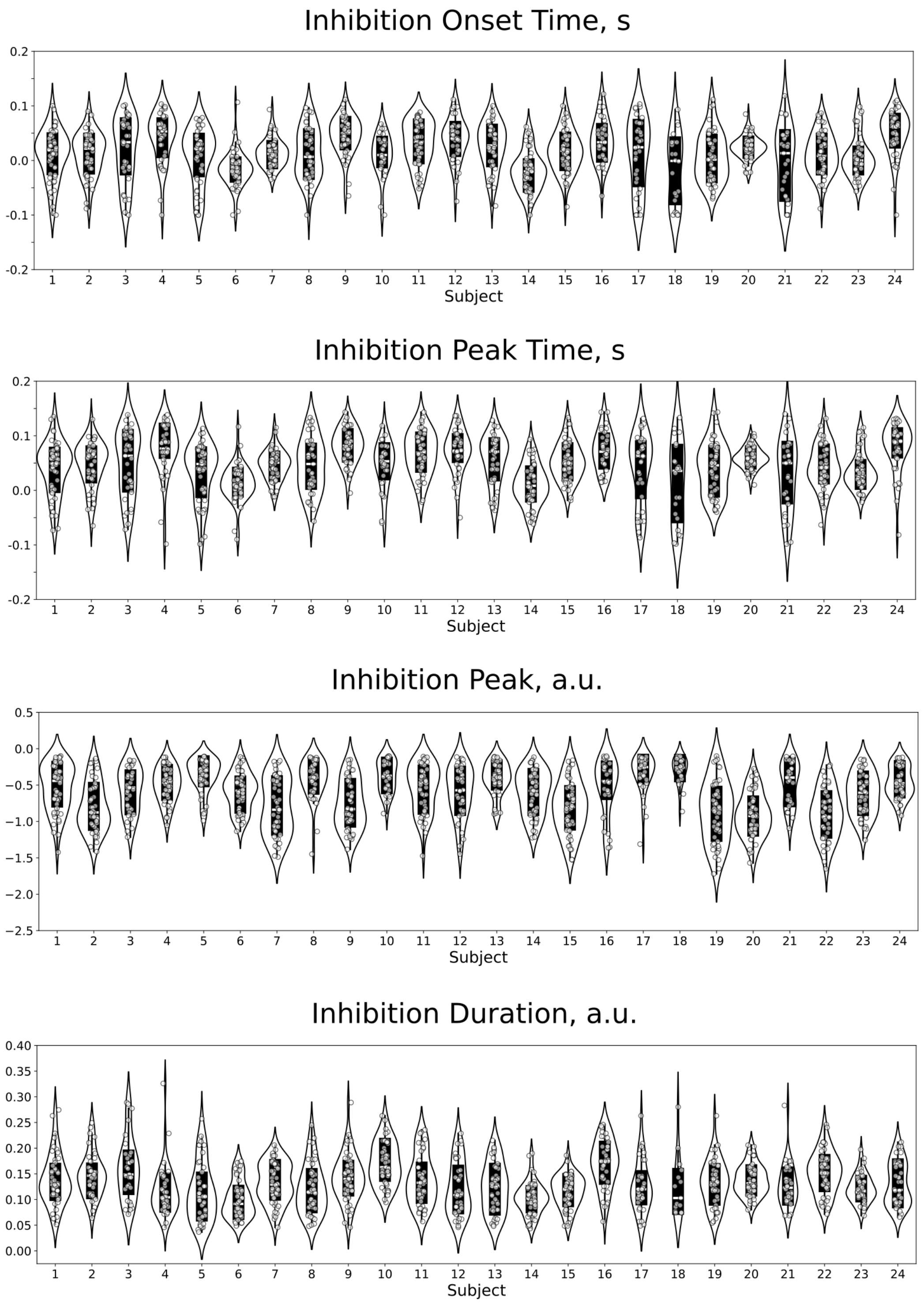


**Figure S5.** Violin plots of *Biceps brachii* inhibition-derived parameters (Inhibition Onset Time, Inhibition Peak Time, Inhibition Peak (strength), Inhibition Duration) for each subject, showing the distribution of values and individual data points.

### Correlation analysis between high-gamma power and *Biceps brachii* inhibition strength

For illustration, we show the correlation between Peak EMG Inhibition and high-gamma (90-130 Hz) power, which was significant in SMA (Fig. 2A), across both hemispheres (Figure S6).

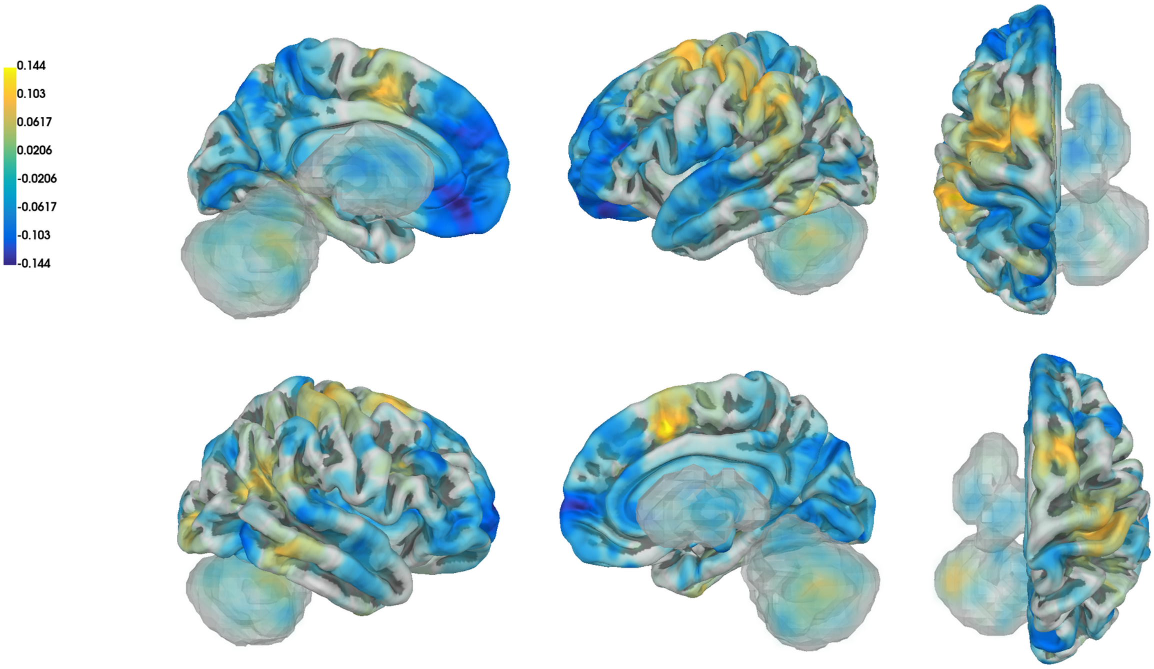


**Figure S6.** The correlation between Peak EMG Inhibition and high-gamma power (90-130 Hz) across both hemispheres.

### GAM: Main effect of *Biceps brachii* inhibition time on the strength of inhibition

Across several GAMs predicting Peak EMG Inhibition from gamma or beta power, with Inhibition Peak Time included to account for temporal dynamics, we consistently observed a significant main effect of Inhibition Peak Time on Peak EMG Inhibition (Table 2). We illustrate this effect using the GAM with high-gamma power (Fig. S7; $F = 12.52$, $p < 0.001$); similar effects were observed across the other models. See Table 2A in the main text for full model details.

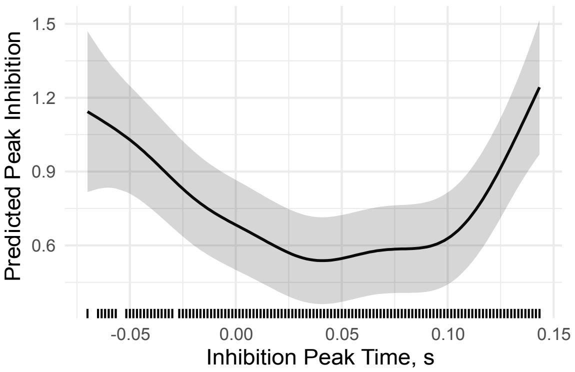


**Figure S7.** Significant main effect of *Biceps brachii* inhibition peak time on the strength of EMG Inhibition (Table 2A). More negative Peak Inhibition values indicate stronger inhibition.

### GAM: Main effect of *Biceps brachii* inhibition time on inhibition duration

In the GAM predicting *Biceps brachii* Inhibition Duration from beta power, with Inhibition Peak Time included to account for temporal dynamics, we observed a significant main effect of Peak Inhibition Time on Inhibition Duration ($F = 4.36$, $p = 0.002$), as illustrated in Figure S8. See details of this model in the main text (Table 2D).

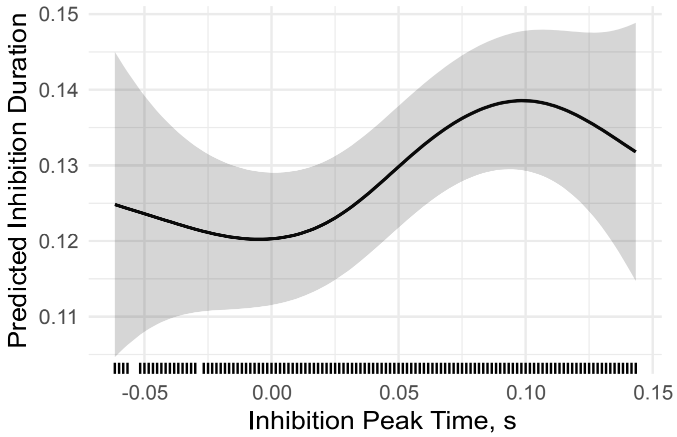


**Figure S8.** Significant main effect of peak *Biceps brachii* inhibition time on inhibition duration (s; see Table 2D).

### GAM: Predicting high-gamma power from 6-30 Hz activity

To test whether the high-gamma suppression linked to stronger anticipatory inhibition (Fig. 2A) is mediated by alpha or beta activity, while accounting for subject-level variability and temporal dynamics, we fitted a GAM predicting high-gamma power (averaged over the 25 ms preceding the inhibition peak) from 6-30 Hz power in the SMA label. The analysis was restricted to anticipatory inhibitions with peaks between -70 and 75 ms relative to unloading, and included 19 participants with a sufficient number of anticipatory trials.

The model included (1) frequency-specific smooth effects of residual 6-30 Hz power (after regressing out broadband 1-130 Hz activity to control for aperiodic contributions), (2) a smooth effect of inhibition peak time to capture nonlinear temporal dynamics, and (3) subject-specific random slopes for inhibition peak time and residual power, and random intercepts, to account for inter-individual differences in both temporal and spectral effects.

Exact model specification:

*Gamma ~ s(Power_res, by=Freq) + s(Inh_peak_time, k=60) +*
*s(Inh_peak_time, Subject, bs="fs") +*
*s(Power_res, Subject, bs = "fs") + s(Subject, bs = "re")*

The model revealed significant frequency-specific effects in the 21-25 Hz range, indicating that beta power predicts high-gamma activity (Fig. S9; Table S2). Notably, a similar significant effect of 21-24 Hz beta power on high-gamma activity was observed when all participants and both anticipatory and reactive inhibitions were included. The model also showed a strong main effect of inhibition peak time on high-gamma power. Significant random intercepts and slopes for both beta power and inhibition peak time indicated substantial inter-individual variability in high-gamma power and in its relationship to beta activity and inhibition timing.

**Table S2.** GAM results on smooth terms predicting high-gamma power on 6-30 Hz power and inhibition peak time.

| Smooth term | edf | Ref.df | F | p-value |
|---|---|---|---|---|
| s(Power_res):Freq6 | 1.000 | 1.000 | 0.100 | 0.752 |
| s(Power_res):Freq7 | 1.000 | 1.000 | 0.160 | 0.689 |
| s(Power_res):Freq8 | 1.000 | 1.000 | 0.178 | 0.673 |
| s(Power_res):Freq9 | 1.000 | 1.000 | 0.441 | 0.507 |
| s(Power_res):Freq10 | 1.000 | 1.000 | 0.348 | 0.555 |
| s(Power_res):Freq11 | 1.000 | 1.000 | 0.054 | 0.816 |
| s(Power_res):Freq12 | 1.000 | 1.000 | 0.012 | 0.913 |

| Smooth term | edf | Ref.df | F | p-value |
|---|---|---|---|---|
| s(Power_res):Freq13 | 1.000 | 1.000 | 0.200 | 0.655 |
| s(Power_res):Freq14 | 1.000 | 1.000 | 0.132 | 0.716 |
| s(Power_res):Freq15 | 1.582 | 1.977 | 0.264 | 0.750 |
| s(Power_res):Freq16 | 1.000 | 1.000 | 0.092 | 0.762 |
| s(Power_res):Freq17 | 1.000 | 1.000 | 0.001 | 0.973 |
| s(Power_res):Freq18 | 1.000 | 1.000 | 0.002 | 0.967 |
| s(Power_res):Freq19 | 1.000 | 1.000 | 0.048 | 0.826 |
| s(Power_res):Freq20 | 1.000 | 1.000 | 1.517 | 0.218 |
| s(Power_res):Freq21 | 1.000 | 1.000 | 6.067 | 0.0138* |
| s(Power_res):Freq22 | 1.000 | 1.000 | 8.038 | 0.0046** |
| s(Power_res):Freq23 | 1.000 | 1.000 | 7.632 | 0.0057** |
| s(Power_res):Freq24 | 1.001 | 1.002 | 7.504 | 0.0061** |
| s(Power_res):Freq25 | 1.000 | 1.000 | 4.415 | 0.0356* |
| s(Power_res):Freq26 | 1.000 | 1.000 | 1.379 | 0.240 |
| s(Power_res):Freq27 | 1.000 | 1.000 | 0.305 | 0.581 |
| s(Power_res):Freq28 | 1.000 | 1.000 | 0.130 | 0.719 |
| s(Power_res):Freq29 | 1.000 | 1.001 | 0.026 | 0.874 |
| s(Power_res):Freq30 | 1.480 | 1.827 | 0.313 | 0.731 |
| s(Inh_peak_time) | 54.804 | 56.798 | 23.824 | <2e−16*** |
| s(Subject) | 10.746 | 18.000 | 1.611 | <2e−16*** |
| s(Inh_peak_time, Subject) | 139.690 | 184.000 | 752.131 | 0.0028** |
| s(Power_res, Subject) | 42.057 | 188.000 | 0.843 | <2e−16*** |

Inh_peak_time: EMG Inhibition Peak Time; Power_res: residual power after regressing out 1-130 Hz broadband activity; s(Inh_peak_time, Subject, bs="fs"), s(Power_res, Subject, bs="fs"): subject-specific factor smooths allowing the effect of each variable to vary across subjects; s(Subject, bs="re"): subject-specific random intercept.

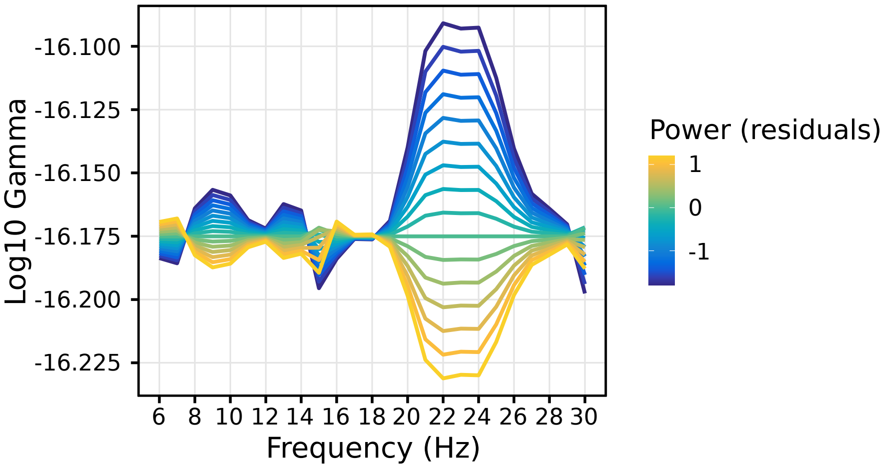


**Figure S9.** Significant main effect of 21-25 Hz power on high-gamma power for data averaged over the 25 ms preceding the *Biceps brachii* inhibition peak.

### Distribution of burst trials across subjects

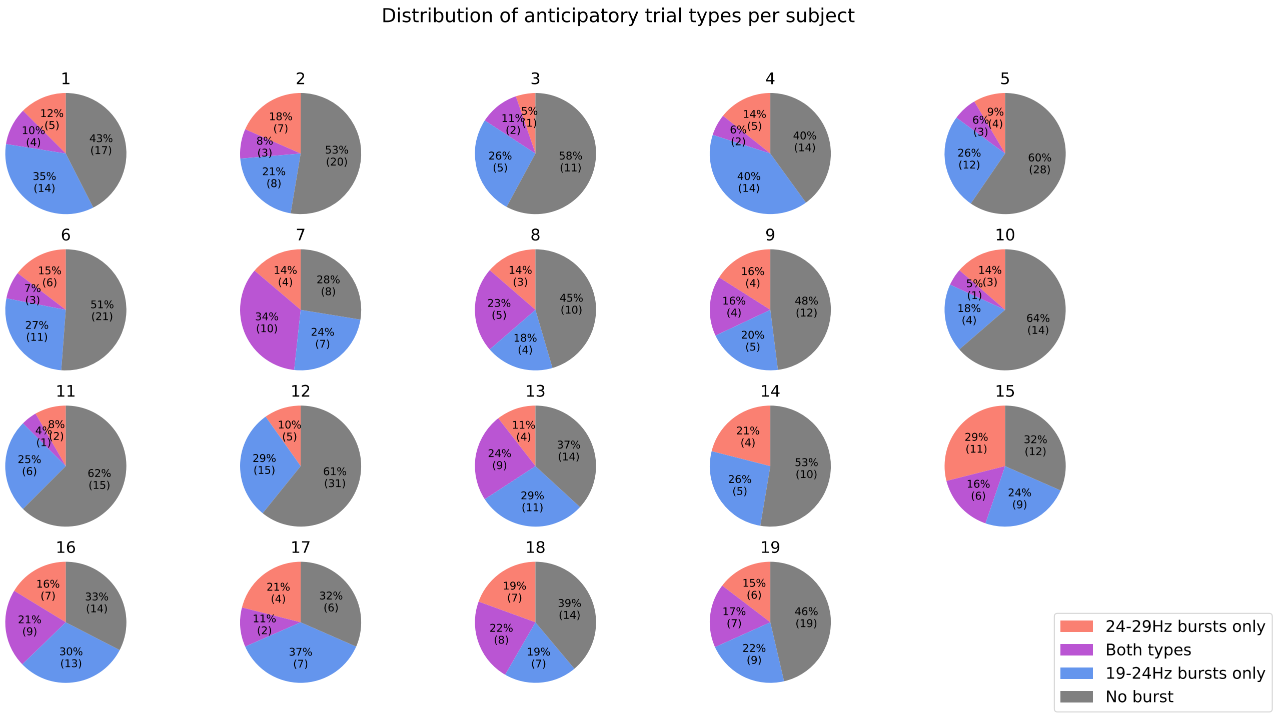


**Figure S10.** Illustration of the absolute number and percentage of trials containing beta bursts in each category (19-24 Hz bursts, 24-29 Hz bursts, or both) for each subject.

### Brain-level analyses on anticipatory *Biceps brachii* inhibition

***High-gamma power modulation around inhibition peak***

In addition to the correlation between high-gamma power and *Biceps brachii* inhibition strength reported in the main manuscript (Fig. 2A), we tested whether high-gamma power in selected cortical and subcortical regions is modulated around the time of *Biceps brachii* inhibition, which would suggest a potential contribution to muscle inhibition.

For this analysis, high-gamma power (90-130 Hz, averaged across frequencies) was aligned to the inhibition peak across trials and compared to baseline activity (-1.6 to -1.3 s). Only trials with inhibition peaks occurring between -70 and 150 ms relative to unloading were included.

A permutation cluster test over the -100 to 50 ms window revealed a significant increase in high-gamma power in ventral PMC/M1, with a peak cluster spanning -36 to 23 ms (Fig. S11). To assess temporal specificity, we fitted a GAM predicting high-gamma power in this region from inhibition peak time, including subject-specific random intercepts and slopes. No significant effect of inhibition time was observed, suggesting that this excitability increase is not specific to the anticipatory period (Table S3, Fig. S11). A significant random intercept indicates inter-individual differences in overall high-gamma power levels across participants.

Exact model specification:

Gamma_m1 ~ s(Inh_peak_time) + s(Subject, Inh_peak_time, bs = "re") + s(Subject, bs = "re")

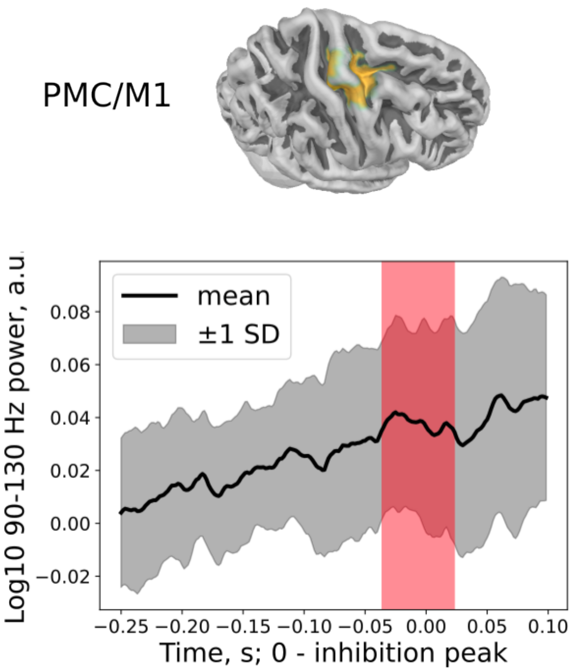


**Figure S11.** PMC/M1 cluster showing increased high-gamma power (90-130 Hz) around inhibition peak. The plot shows the grand-average log10-transformed, baseline-corrected power averaged across cluster vertices; grey shading indicates between-subject standard deviation, and the red shaded area indicates the significant time interval.

**Table S3.** GAM results on smooth terms predicting PMC/M1 high-gamma power from inhibition peak time.

| Smooth term | edf | Ref.df | F | p-value |
|---|---|---|---|---|
| s(Inh_peak_time) | 1.01 | 1.02 | 0.10 | 0.761 |
| s(Subject) | 9.08 | 18.00 | 1.01 | 0.008** |
| s(Subject, Inh_peak_time) | <0.001 | 18.00 | <0.001 | 0.455 |

Inh_peak_time: EMG Inhibition Peak Time; s(Subject, Inh_peak_time, bs="re"): random slope for EMG Inhibition Peak Time by subject; s(Subject, bs="re"): random intercept by subject.

***Correlational analysis***

To test whether cortical or subcortical regions contribute to anticipatory inhibition of the postural *Biceps brachii* via alpha- or beta-band activity, we estimated correlations between Peak EMG Inhibition and periodic power in the 6-30 Hz range across right and left cortical regions, the right basal ganglia, and bilateral cerebellum, taking into account the bimanual nature of the task. We additionally assessed correlations between 6-30 Hz power and EMG Inhibition Duration, suggesting that a broader network may contribute to modulation of anticipatory inhibition duration via alpha/beta activity.

Correlations were computed at each time-frequency point within the -0.15 to 0 s interval relative to the inhibition peak, separately for each source. This earlier time window, compared with that used for the high-gamma correlation with Peak EMG Inhibition (Fig. 2A), was chosen to account for the possibility that some regions exert indirect control that may precede the peak effect. One-tailed permutation cluster tests were conducted separately for the right cerebral cortex, left cerebral cortex, and subcortical regions. These tests assessed whether correlation values were significantly less than zero for Peak EMG Inhibition correlation, indicating that stronger anticipatory muscle inhibition was associated with increased periodic alpha-beta power, and significantly greater than zero for EMG Inhibition Duration correlation, suggesting that higher periodic alpha-beta power was associated with longer inhibition duration.

For Peak EMG Inhibition, a significant negative correlation cluster was observed in the right cerebral cortex, with the maximal effect localized to the right inferior frontal cortex (IFC). The cluster spanned the full -0.15 to 0 s interval and the 14-19 Hz range ($t_{min}$ = -6.42, p = 0.040; Fig. S12A). No significant effects were observed in the left cortex ($t_{min}$ = -6.15, p = 0.21), right cerebellum ($t_{min}$ = -3.98, p = 0.97), left cerebellum ($t_{min}$ = -4.39, p = 0.83), or right basal ganglia ($t_{min}$ = -4.15, p = 0.45). No significant relationships were also observed for the correlation between inhibition duration and 6-30 Hz power in any region, including the right cerebral cortex ($t_{max}$ = 6.67, p = 0.52), left cerebral cortex ($t_{max}$ = 5.69, p = 0.21), right cerebellum ($t_{max}$ = 4.57, p = 0.17), left cerebellum ($t_{max}$ = 3.23, no clusters), or right basal ganglia ($t_{max}$ = 3.52, no clusters).

To explore the temporal dynamics of the relationship between beta power and Peak EMG Inhibition in the IFC cluster, we fitted a GAM predicting Peak EMG Inhibition from 14-19 Hz beta power and Inhibition Peak Time. The model showed a significant main effect of Inhibition Peak Time but no main effect of beta power. However, there was a significant interaction between beta power and Inhibition Peak Time (Table S3), indicating that beta power explains most variance in Peak EMG Inhibition when inhibition occurs in the early anticipatory period (Fig. S12A). Significant random intercepts indicated inter-individual differences in IFC beta power. Together, these results support the correlation analysis and highlight the anticipatory nature of the beta-muscle inhibition relationship in IFC.

**Table S3.** GAM results on smooth terms predicting *Biceps brachii* inhibition strength on 14-19 Hz beta power and inhibition peak time in IFC (Fig. S12A).

| Term | **edf** | **Ref.df** | **F** | **p** |
|---|---|---|---|---|
| *Peak EMG Inhibition ~ IFC Beta + Inhibition peak time + IFC Beta x Inhibition Peak time* | | | | |
| s(Beta_IFC) | 1.000 | 1.000 | 1.450 | 0.2288 |
| s(Inh_peak_time) | 5.263 | 6.375 | 12.867 | < 0.001 *** |
| ti(Beta_IFC, Inh_peak_time) | 1.819 | 2.261 | 4.387 | 0.0124 * |
| s(Subject, Beta_IFC) | 2.359 | 23.000 | 0.117 | 0.2972 |
| s(Subject, Inh_peak_time) | 7.336 | 23.000 | 6.161 | 0.3246 |
| s(Subject) | 21.917 | 23.000 | 46.066 | < 0.001 *** |

Inh_peak_time: EMG Inhibition Peak Time; Beta_IFC: 14-19 Hz beta power; ti(Beta_IFC, Inh_peak_time): tensor product interaction between smooth terms; s(Subject, Inh_peak_time, bs="re"), s(Subject, Beta_IFC, bs="re"): random slopes by subject; s(Subject, bs="re"): random intercept by subject.

***Phase-locking analysis***

To identify brain regions potentially involved in the timing of *Biceps brachii* inhibition, we performed phase-locking analysis, based on the assumption that if a given region contributes to the timing of muscle suppression, the phase of its oscillatory activity at a given frequency should be consistent across trials when aligned to inhibition onset.

To assess phase consistency, we estimated the phase-locking value (PLV) for trials exhibiting anticipatory *Biceps brachii* inhibition, using data centered on the EMG inhibition onset. As a control condition, we computed the average PLV during a baseline period (-1.6 to -1.3 s relative to unloading) from the same trials. The PLV was calculated as previously described (Lachaux et al., 1999).

PLV was estimated in the 6-30 Hz range, suggesting that the phase of alpha- or beta-band activity may be associated with the timing of inhibition. One-tailed permutation cluster tests were performed on the contrast between PLV computed from data aligned to inhibition onset and PLV from a baseline interval to identify brain regions where alpha-beta PLV was significantly higher in the 100 ms preceding inhibition onset.

Although we primarily expected involvement of right-hemispheric cortical regions contralateral to the postural arm, the basal ganglia and cerebellum have also been implicated in motor timing, suggesting that they may ensure coordination between APA and voluntary movements (Andersen & Dalal, 2024; Massion, 1992; Ng et al., 2013), which may be reflected in phase alignment with inhibition onset. Given the close temporal coupling between APA and voluntary movement onset in adults (Hugon et al., 1982; Massion, 1992), PLV was additionally estimated in the left cortex, proposed to relay timing signals to the right hemisphere (Massion, 1992).

Three significant right cortical clusters were identified (Fig. S12B). The largest peaked in PMC, SMA, and M1, spanning the full 100 ms preceding inhibition onset in the 6-7 Hz range ($t_{max}$ = 6.88, p = 0.004). A second cluster in ventral PMC/M1 spanned -95 ms to onset in the 18-29 Hz range ($t_{max}$ = 6.15, p = 0.019), and a third in the precuneus spanned -65 ms to onset in the 12-19 Hz range ($t_{max}$ = 8.96, p = 0.022).

Significant PLV increases were also observed in the right basal ganglia across three clusters: putamen (-73 ms to onset, 6-7 Hz; $t_{max}$ = 6.11, p = 0.014), globus pallidus (-78 ms to onset, 6-7 Hz; $t_{max}$ = 6.40, p = 0.038), and caudate nucleus (-38 ms to onset, 18-29 Hz; $t_{max}$ = 6.65, p = 0.021; Fig.

S12B). The right cerebellum also showed two significant clusters in its lateral part: one spanning the full -100 to 0 ms interval in the 6-19 Hz range ($t_{max} = 8.47$, $p = 0.0002$), and a second spanning -84 to -28 ms in the 17-30 Hz range ($t_{max} = 7.23$, $p = 0.048$; Fig. S12B).

No significant effects were observed in the left cortex ($t_{max} = 5.89$, $p = 0.18$) or left cerebellum ($t_{max} = 5.84$, $p = 0.20$).

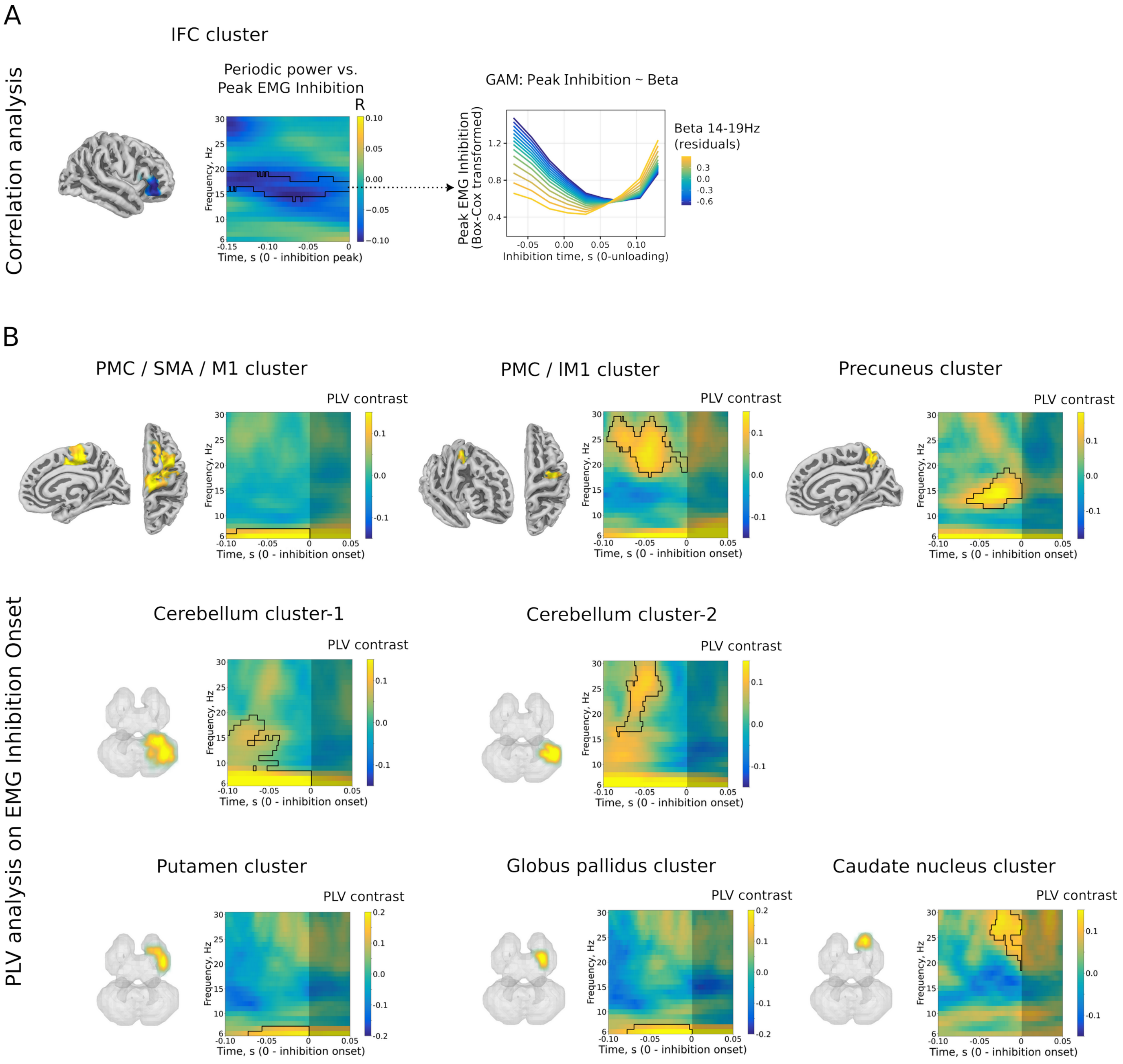


**Figure S12.** Brain-level correlations and phase-locking associated with *Biceps brachii* inhibition strength and onset. **A:** Left panel: right IFC label showing a significant correlation with Peak EMG Inhibition in the beta band and the time-frequency plot showing group-averaged Spearman's R values in the IFC label. Right panel: the corresponding GAM model predicting Peak EMG Inhibition from beta power averaged over the cluster, assessed across Inhibition Peak Time. See Table S3 for full GAM results. **B:** Phase-locking analysis showing significant PLV increases in right cortical and subcortical regions shortly preceding *Biceps brachii* inhibition onset compared with baseline. In panels **A** and **B**, thin black contours indicate significant correlations ($p < 0.05$, corrected). Bright rectangles in panel B indicate the analysis window; shaded areas are shown for visualization purposes.

### Brain-level analysis of lower-beta bursts: subcortical connectivity and beta–high-gamma interactions

We estimated Granger causality between the basal ganglia, cerebellum, and SMA during lower-beta bursts (19-24 Hz) to assess whether subcortical structures contribute to SMA burst dynamics. No directed influence from the right basal ganglia to the SMA was observed ($t_{max} = 3.32$, no clusters). In contrast, both cerebellar hemispheres showed significant directed connectivity to the SMA (Fig. S13): the right cerebellum exhibited clusters spanning -50 to -32 ms at 19-24 Hz ($t_{max} = 4.90$, $p = 0.002$) and -46 to -26 ms at 21-24 Hz ($t_{max} = 4.19$, $p = 0.026$), while the left cerebellum showed a cluster spanning -50 to -30 ms at 20-24 Hz ($t_{max} = 4.37$, $p = 0.032$).

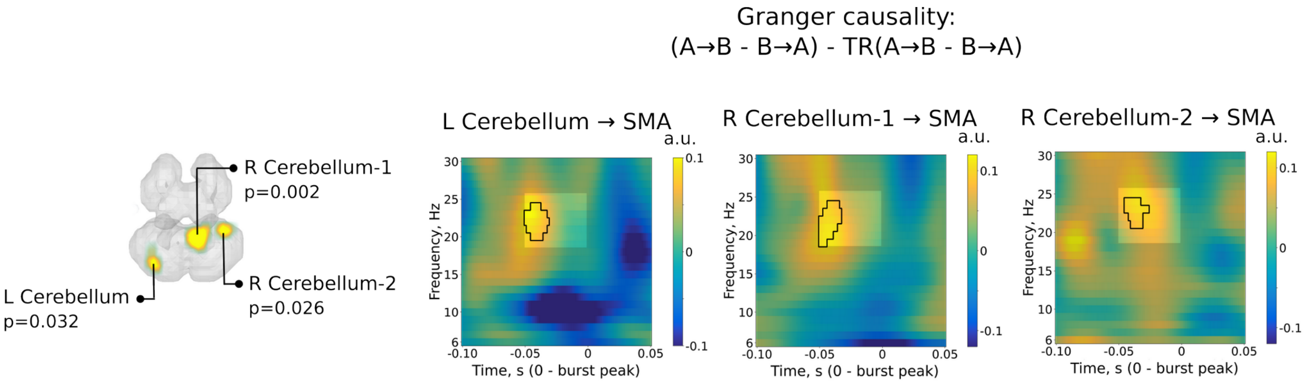


**Figure S13.** Directed connectivity from cerebellar regions to the SMA during lower-beta bursts (19-24 Hz).

### Granger causality estimation between lPFC and PMC/M1

To verify the directional influence from lPFC to PMC/M1 during SMA bursts – both areas that showed directional influence on the SMA during lower-beta (19-24 Hz) bursts – we estimated directed connectivity from lPFC to PMC/M1. In contrast to our previous study in adults (Manyukhina et al., 2026), we did not observe a significant effect in the SMA beta burst range (19-

24 Hz). Instead, the time-frequency distribution of the directed connectivity effect suggests the opposite direction, although the peak effect does not align with the 19-24 Hz beta bursts (Fig. S14).

An exploratory permutation cluster analysis (two-tailed) in the 100 ms before the SMA burst peak, spanning 6-30 Hz, revealed a tendency for directional influence from PMC/M1 to lPFC ($t_{max}$ = 3.06, $p_{min}$ = 0.069).

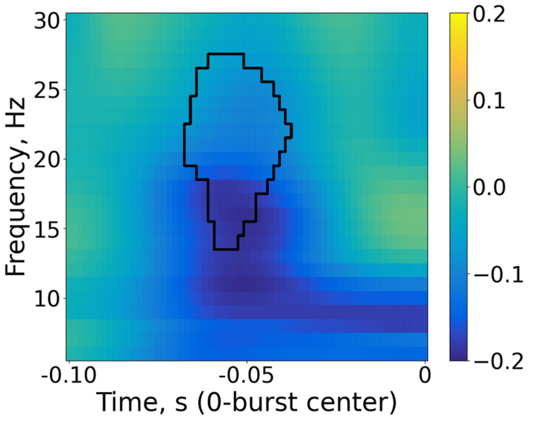


**Figure S14.** Exploratory Granger causality from lPFC to PMC/M1, indicating a tendency ($p < 0.15$) for directional connectivity in the opposite direction to that observed in adults (Manyukhina et al., 2026): from PMC/M1 to lPFC.

### Temporal profiles of identified burst types

For illustrative purposes, we examined the temporal profiles of lower- and higher-beta bursts detected in the SMA label. LCMV timecourses from the four SMA vertices were centered on burst peaks and bandpass-filtered within ±3 Hz around the respective burst frequency range (16-27 Hz for lower-beta; 21-32 Hz for higher-beta bursts). The resulting vertex-wise and label-averaged timecourses for both burst types are shown in Figure S15.

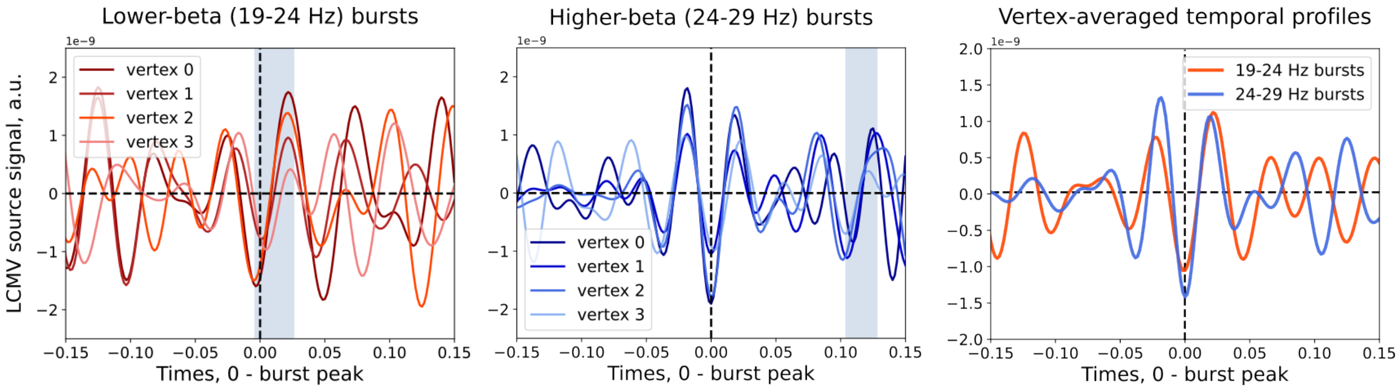

**Figure S15.** Vertex-wise and label-averaged timecourses of lower- and higher-beta bursts in the SMA label. Grey shaded intervals indicate periods of significant high-gamma suppression during each burst type.

### Relationship between burst-related high-gamma power and behavioural measures

#### *GAMs*

In the main manuscript, we assessed the relationship between high-gamma suppression during each burst type and EMG- and elbow rotation-based parameters using GAMs. Model specifications and results for the three GAMs, fitted separately for lower- and higher-beta bursts, are summarized in Table S4 and illustrated in Figure S16.

**Table S4.** GAM results for linear fixed and smooth terms predicting high-gamma suppression during each burst type from: A) Peak EMG Inhibition, EMG Inhibition Peak Time, and EMG Rebound; B) EMG Inhibition Onset and EMG Inhibition Duration; C) Peak Elbow Rotation and Elbow Rotation Decline.

***A.*** *Gamma ~ Inh_ampl + Inh_peak_time + Inh_rebound + s(Subject, Inh_ampl, bs = "re") + s(Subject, Inh_peak_time, bs = "re") + s(Subject, Inh_rebound, bs = "re") + s(Subject, bs = "re")*

***Lower-beta bursts***

| **Parametric term** | **Estimate** | **SE** | **t** | **p** |
|---|---|---|---|---|
| Intercept | -16.13 | 0.106 | -152.76 | <0.001 *** |
| Inh_ampl | 0.168 | 0.085 | 1.98 | 0.048 * |
| Inh_peak_time | 0.831 | 0.665 | 1.25 | 0.212 |
| Inh_rebound | 0.118 | 0.110 | 1.08 | 0.280 |
| **Smooth term** | **edf** | **Ref.df** | **F** | **p** |
| s(Subject, Inh_ampl) | 4.27 | 18 | 13.73 | 0.163 |
| s(Subject, Inh_peak_time) | <0.001 | 18 | 0.00 | 0.876 |
| s(Subject, Inh_rebound | 3.50 | 18 | 0.50 | 0.408 |
| s(Subject) | 16.02 | 18 | 65.34 | <0.001 *** |

***Higher-beta bursts***

***A.*** *Gamma ~ Inh_ampl + Inh_peak_time + Inh_rebound + s(Subject, Inh_ampl, bs = "re") + s(Subject, Inh_peak_time, bs = "re") + s(Subject, Inh_rebound, bs = "re") + s(Subject, bs = "re")*

| **Parametric term** | **Estimate** | **SE** | **t** | **p** |
|---|---|---|---|---|
| Intercept | -16.28 | 0.110 | -148.33 | <0.001 *** |
| Inh_ampl | 0.072 | 0.071 | 1.01 | 0.313 |
| Inh_peak_time | 0.802 | 0.614 | 1.31 | 0.192 |
| Inh_rebound | 0.244 | 0.086 | 2.85 | 0.0047 ** |
| **Smooth term** | **edf** | **Ref.df** | **F** | **p** |
| s(Subject, Inh_ampl) | 3.22 | 18 | 7.65 | 0.363 |
| s(Subject, Inh_peak_time) | <0.001 | 18 | 0.00 | 0.862 |
| s(Subject, Inh_rebound) | <0.001 | 18 | 0.00 | 0.697 |
| s(Subject) | 16.84 | 18 | 56.91 | <0.001 *** |

***B.*** *Gamma ~ Inh_onset_time + Inh_duration + s(Subject, Inh_duration, bs = "re") + s(Subject, Inh_onset_time, bs = "re") + s(Subject, bs = "re")*

***Lower-beta bursts***

| **Parametric term** | **Estimate** | **SE** | **t** | **p** |
|---|---|---|---|---|
| Intercept | -16.20 | 0.123 | -131.37 | <0.001 *** |
| Inh_onset_time | -0.049 | 0.803 | -0.06 | 0.951 |
| Inh_duration | -0.220 | 0.543 | -0.41 | 0.686 |
| **Smooth term** | **edf** | **Ref.df** | **F** | **p** |
| s(Subject, Inh_duration) | <0.001 | 18 | 0.00 | 0.930 |
| s(Subject, Inh_onset_time) | 1.29 | 18 | 0.35 | 0.259 |
| s(Subject) | 17.07 | 18 | 27.23 | <0.001 *** |

***Higher-beta bursts***

| **Parametric term** | **Estimate** | **SE** | **t** | **p** |
|---|---|---|---|---|
| Intercept | -16.27 | 0.115 | -141.39 | <0.001 *** |
| Inh_onset | 1.526 | 0.710 | 2.15 | 0.032 * |
| Inh_duration | 0.641 | 0.563 | 1.14 | 0.256 |
| **Smooth term** | **edf** | **Ref.df** | **F** | **p** |
| s(Subject, Inh_duration) | 5.24 | 18 | 23.61 | 0.308 |

***B.*** *Gamma ~ Inh_onset_time + Inh_duration + s(Subject, Inh_duration, bs = "re") + s(Subject, Inh_onset_time, bs = "re") + s(Subject, bs = "re")*

| | | | | |
|---|---|---|---|---|
| s(Subject, Inh_onset) | <0.001 | 17 | 0.00 | 0.997 |
| s(Subject) | 14.92 | 18 | 54.40 | <0.001 *** |

***C.*** *Gamma ~ PeakElbRot + ElbRotDecl + s(Subject, PeakElbRot, bs = "re") + s(Subject, ElbRotDecl, bs = "re") + s(Subject, bs = "re")*

***Lower-beta bursts***

| **Parametric term** | **Estimate** | **SE** | **t** | **p** |
|---|---|---|---|---|
| Intercept | -16.24 | 0.109 | -149.30 | <0.001 *** |
| PeakElbRot | 2.46 | 5.22 | 0.47 | 0.637 |
| ElbRotDecl | -2.57 | 22.60 | -0.11 | 0.910 |
| **Smooth term** | **edf** | **Ref.df** | **F** | **p** |
| s(Subject, PeakElbRot) | <0.001 | 18 | 0.00 | 0.837 |
| s(Subject, ElbRotDecl) | <0.001 | 18 | 0.00 | 0.786 |
| s(Subject) | 17.08 | 18 | 24.77 | <0.001 *** |

***Higher-beta bursts***

| **Parametric term** | **Estimate** | **SE** | **t** | **p** |
|---|---|---|---|---|
| Intercept | -16.31 | 0.102 | -159.98 | <0.001 *** |
| PeakElbRot | 11.93 | 4.26 | 2.80 | 0.0054 ** |
| ElbRotDecl | 7.42 | 29.23 | 0.25 | 0.800 |
| **Smooth term** | **edf** | **Ref.df** | **F** | **p** |
| s(Subject, PeakElbRot) | <0.001 | 18 | 0.00 | 0.569 |
| s(Subject, ElbRotDecl) | <0.001 | 18 | 0.00 | 0.544 |
| s(Subject) | 17.11 | 18 | 23.06 | <0.001 *** |

PeakElbRot: Peak Elbow Rotation; ElbRotDecl: Elbow Rotation Decline; Inh_ampl: Peak EMG Inhibition; Inh_peak_time: EMG Inhibition Peak time; Inh_rebound: EMG Rebound; Inh_onset_time: EMG Inhibition Onset time; Inh_duration: EMG Inhibition duration; s(Subject, Inh_ampl, bs = "re"): subject-based random slope; s(Subject, bs = "re"): subject-based random intercept.

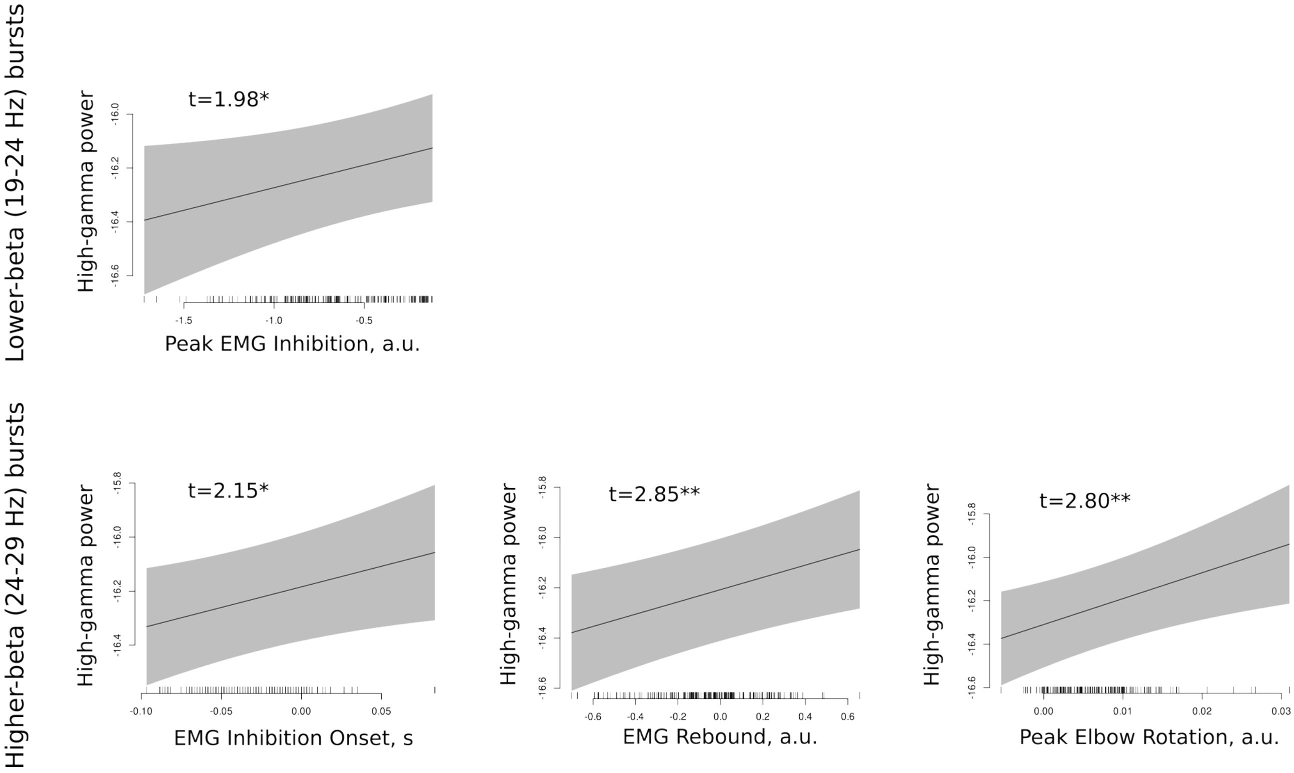


**Figure S16.** Visualization of significant fixed effects from GAMs predicting high-gamma suppression from behavioural measures during lower- and higher-beta bursts (Table S4).

***Spearman's correlation***

To validate the GAM results, Spearman's correlations between the same behavioral measures and high-gamma suppression were estimated for each burst type. As with the GAMs, correlational analyses were restricted to 19 participants with sufficient anticipatory inhibition trials, with two more excluded from the inhibition onset and duration correlations during higher-beta bursts due to insufficient trials.

Consistent with GAM results, greater high-gamma suppression during lower-beta bursts correlated with stronger anticipatory muscle inhibition ($t(18) = 2.8$, $p = 0.011$); no significant correlations with other behavioural parameters were observed. For higher-beta bursts, high-gamma suppression correlated with earlier EMG Inhibition Onset ($t(16) = 2.33$, $p = 0.033$), lower EMG Rebound ($t(18) = 2.59$, $p = 0.018$), and lower Peak Elbow Rotation ($t(18) = 2.27$, $p = 0.036$). Altogether, these correlations replicated the GAM results on the link between high-gamma suppression and behavioral measures (see correlation distributions in Figure S17, middle panels).

To assess a broader temporal effect of high-gamma suppression on forearm dynamics, correlations between high-gamma suppression during each burst type and elbow rotation were computed at each time point from 0 to 750 ms after unloading, followed by one-tailed permutation cluster tests to identify intervals at which stronger high-gamma suppression was associated with reduced elbow rotation.

Significant correlations were found for both burst types (Fig. S17, right panels). For lower-beta bursts, maximal clusters spanned 86-96 ms and 408-750 ms after unloading ($t_{max}$ = 3.05, $p_{min}$ = 0.028), and for higher-beta bursts, 118-134 ms and 310-424 ms ($t_{max}$ = 3.34, $p_{min}$ = 0.023). These results indicate that stronger burst-related high-gamma suppression is associated with forearm stabilization during both the raising and falling phases of elbow rotation, though the timing of the later effects suggests they are likely indirect. Interestingly, the late stabilization associated with lower-beta bursts may reflect recruitment of mechanisms ensuring return of the forearm to its initial position – an effect not present for higher-beta bursts, which instead appear implicated in compensating for forearm destabilization, reflected in suppressing maximal elbow rotation after unloading.

Given the low trial counts per burst type in some participants, which limits the reliability of trial-wise correlation estimates, these correlational analyses should be considered supplementary, with GAMs providing more robust group-level estimates.

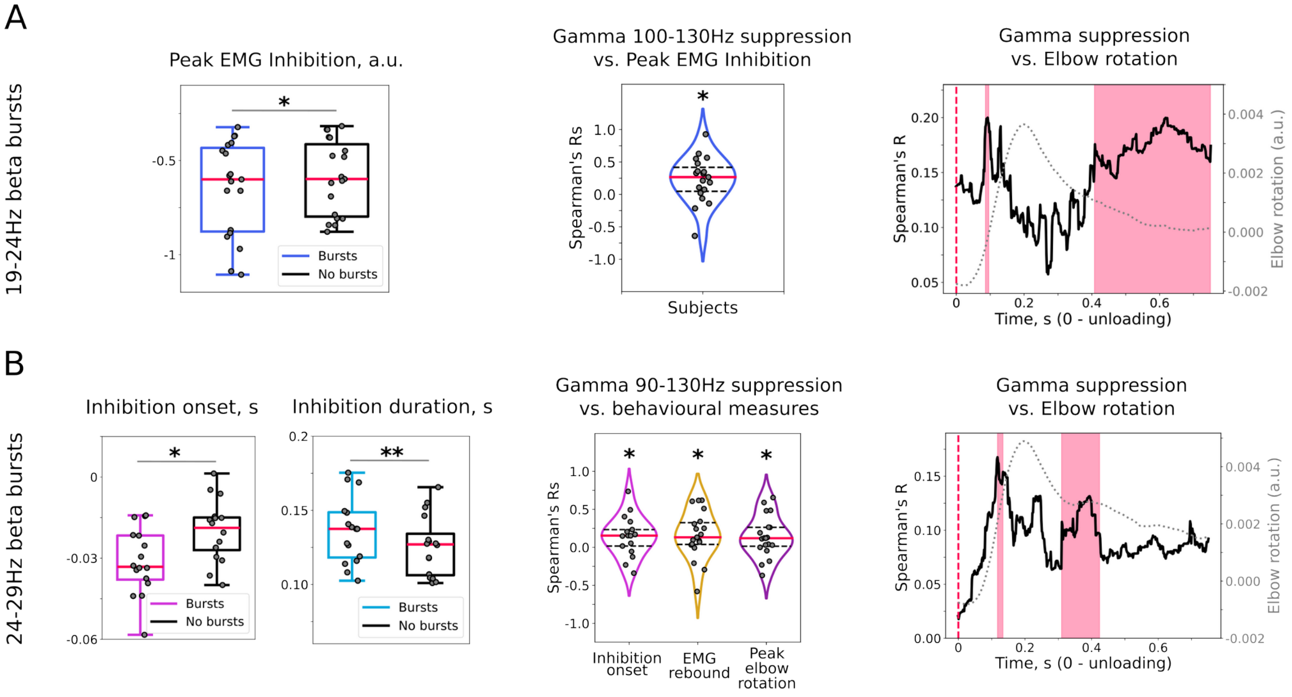


**Figure S17.** Effects of beta bursts and high-gamma suppression on *Biceps brachii* inhibition and elbow rotation measures. Panels **A** and **B** show analogous analyses for the two burst types: 19-24 Hz (**A:** "lower-beta") and 24-29 Hz (**B:** "higher-beta"). Left plots: differences in behavioral measures between burst and no-burst trials. Middle plots: correlations between high-gamma suppression and behavioral measures across burst trials. Right plots: correlations between high-gamma suppression and elbow rotation amplitude over time. Thick black lines show the group-average correlation time course and red-shaded areas indicate significant time spans ($p < 0.05$, corrected). Grey dotted lines represent elbow rotation dynamics, illustrating their alignment with the correlation time course.

# $p<0.15$, * $p<0.05$, ** $p<0.01$

### References


Andersen, L. M., & Dalal, S. S. (2024). The role of the cerebellum in timing. *Current Opinion in Behavioral Sciences*, *59*, 101427. https://doi.org/10.1016/j.cobeha.2024.101427

Hugon, M., Massion, J., & Wiesendanger, M. (1982). Anticipatory postural changes induced by active unloading and comparison with passive unloading in man. *Pflügers Archiv European Journal of Physiology*, *393*(4), 292–296. https://doi.org/10.1007/BF00581412

Lachaux, J.-P., Rodriguez, E., Martinerie, J., & Varela, F. J. (1999). Measuring phase synchrony in brain signals. *Human Brain Mapping*, *8*(4), 194–208. https://doi.org/10.1002/(SICI)1097-0193(1999)8:4<194::AID-HBM4>3.0.CO;2-C

Lotze, M., Erb, M., Flor, H., Huelsmann, E., Godde, B., & Grodd, W. (2000). fMRI Evaluation of Somatotopic Representation in Human Primary Motor Cortex. *NeuroImage*, *11*(5), 473–481. https://doi.org/10.1006/nimg.2000.0556

Manyukhina, V., Abdoun, O., Di Rienzo, F., Barlaam, F., Daligault, S., Delpuech, C., Szul, M., Bonaiuto, J., Bonnefond, M., & Schmitz, C. (2026). Beta bursts in SMA mediate anticipatory muscle inhibition. *Cerebral Cortex*, *36*(6), bhag054. https://doi.org/10.1093/cercor/bhag054

Massion, J. (1992). Movement, posture and equilibrium: Interaction and coordination. *Progress in Neurobiology*, *38*(1), 35–56. https://doi.org/10.1016/0301-0082(92)90034-C

Ng, T. H. B., Sowman, P. F., Brock, J., & Johnson, B. W. (2013). Neuromagnetic imaging reveals timing of volitional and anticipatory motor control in bimanual load lifting. *Behavioural Brain Research*, *247*, 182–192. https://doi.org/10.1016/j.bbr.2013.03.020

Plow, E. B., Arora, P., Pline, M. A., Binenstock, M. T., & Carey, J. R. (2010). Within-limb somatotopy in primary motor cortex – revealed using fMRI. *Cortex*, *46*(3), 310–321. https://doi.org/10.1016/j.cortex.2009.02.024
